# A semi-automated organoid screening method demonstrates epigenetic control of intestinal epithelial differentiation

**DOI:** 10.1101/2020.07.23.217414

**Authors:** Jenny Ostrop, Rosalie Zwiggelaar, Marianne Terndrup Pedersen, François Gerbe, Korbinian Bösl, Håvard T. Lindholm, Alberto Díez-Sánchez, Naveen Parmar, Silke Radetzki, Jens Peter von Kries, Philippe Jay, Kim B. Jensen, Cheryl Arrowsmith, Menno J. Oudhoff

## Abstract

Intestinal organoids are an excellent model to study epithelial biology. Yet, the selection of analytical tools to accurately quantify heterogeneous organoid cultures remains limited. Here, we developed a semi-automated organoid screening method, which we applied to a library of highly specific chemical probes to identify epigenetic regulators of intestinal epithelial biology. The role of epigenetic modifiers in adult stem cell systems, such as the intestinal epithelium, is still undefined. Based on this resource data, we identified several targets that affected epithelial cell differentiation, including HDACs, EP300/CREBBP, LSD1, and type I PRMTs, which were verified by complementary methods. For example, we show that inhibiting type I PRMTs, which leads enhanced epithelial differentiation, blocks the growth of adenoma but not normal organoid cultures. Thus, epigenetic probes are powerful tools to study intestinal epithelial biology and may have therapeutic potential.

## Introduction

The intestinal epithelium, a single layer of cells, faces the challenge of both providing a barrier against pathogens while also being responsible for the uptake of nutrients and water. One of the hallmarks of intestinal epithelium is the rapid turnover of 3-5 days, which is driven by LGR5+ intestinal stem cells (ISCs) that reside at the bottom of crypts. ISCs are continuously dividing and give rise to progenitor cells, which differentiate into specialized intestinal epithelial cell (IEC) lineages such as absorptive enterocytes and secretory lineages such as mucus-producing goblet cells, antimicrobial-producing Paneth cells, hormone-secreting enteroendocrine cells, and chemosensory tuft cells ^1^. The intestinal epithelium exhibits high plasticity in respond to challenges ^2,3^. On the other hand, it is vulnerable to tumorigenesis with colorectal cancer being the second leading cause of cancer deaths world-wide.

The balance between ISC proliferation and IEC differentiation is controlled by pathways including WNT, BMP and NOTCH ^1^. Specific transcription factors, such as ATOH1, are critically required for acquisition of IEC effector lineage identities ^4^. Gene expression is further determined by the chromatin landscape. It is known that epigenetic marks such as methylated DNA and histone tail modifications differ strongly between fetal and adult intestine ^5,6^, and can be altered in intestinal pathologies ^7^. While the requirement of epigenetic modifications for embryonic stem cell differentiation ^8^ and differentiation and maturation of immune cells ^9^ has been extensively studied, their role for maintenance of intestinal homeostasis is debated. Both a permissive chromatin structure and regulation of IEC lineage differentiation by transcription factors, and a control of gene expression patterns by the chromatin states itself have been proposed as conflicting models (extensively reviewed by Elliot et al. ^10^). The classic NOTCH-mediated lateral inhibition model of ISC-to-IEC differentiation has been attributed to a broadly permissive chromatin landscape, supporting the idea of regulation by transcription factors as the most defining factor ^11^. However, other studies suggest that ISC differentiation and the de-differentiation of lineage-defined IECs back to ISCs are mediated by changes in DNA methylation and chromatin accessibility ^3,5,12,13^. Several hundred epigenetic modification enzymes contribute to writing, erasing, and reading the epigenetic code ^14^. Currently, the investigation of the role of epigenetic modifiers in the intestinal epithelium depends mostly on labour-intensive mouse models with conditional genetic deletion, allowing for the examination of one or few epigenetic modifiers at the same time ^15,16^. A higher throughput could be achieved by using organoids to investigate epigenetic effects in the intestinal epithelium ^17^. Curated by the Structural Genomics Consortium, an openly accessible chemical probe library targeting epigenetic modification enzymes with high selectivity and specificity became available recently ^18,19^. Treating organoids with this chemical probe library will enable a direct comparison of the putative requirement of many epigenetic modifiers for epithelial homeostasis or differentiation of IEC lineages.

Heterogeneous organoid cultures are quite sensitive to subtle changes in handling and culture conditions. Therefore, development of quantitative analysis methods for reproducible quantification of a whole organoid population instead of relying on representative example data points is crucial ^20^. Indeed, this has recently led to specialized studies such as using light-sheet microscopy to elegantly define symmetry breaking ^21^, using single-cell RNA sequencing (scRNA-seq) to describe epithelial responses to immune cues ^2,22^, or analysis of single intestinal organoids in microcavity arrays ^23^. However, these techniques are costly and the required instrumentation and data analysis pipelines are not widely available to the research community. Thus, quantitative but cost-efficient tools based on standard laboratory equipment that can be scaled to screen setups need to be established.

Here, we provide a semi-automated organoid quantification method suitable for screening experiments and designed to be used in laboratories with a standard infrastructure. To widely investigate the role of epigenetic modifiers for adult intestinal epithelial homeostasis, we combined this toolbox with a chemical probe library consisting of 39 inhibitors that target epigenetic modification enzymes with high selectivity and specificity ^18^. From this screen dataset, we identified several mediators of IEC biology that we verified with complementary methods. We envision that this resource will be useful for the research community and will lay basis for further mechanistic investigation. Specifically, we find new regulators of organoid size related to ISC frequency, as well as new regulators of IEC differentiation. Finally, we explore the potential of some of these probes for treatment of intestinal cancer by application on intestinal tumor organoids.

## Results

### Development of a toolbox to quantify intestinal organoid growth and cellular composition

A decade after its establishment by Sato et al. ^24^, the use of intestinal organoids has been become a standard in the method repertoire. However, accurately quantifying heterogeneous organoid cultures remains a challenge and the analytical tools available to a broad community, especially for screening purposes, remain limited or labour intensive. We thus initiated a small intestinal (SI) organoid system that, similar to the original work, starts with freshly isolated crypts that self-organize into budding organoids by day 4 (96h after seeding), which can be split and propagated (**Fig. 1a, Supplementary Fig. S1a**). We next designed a setup to daily acquire bright-field z-stack images of the whole extracellular matrix (Matrigel droplet) in a well, followed by automatic segmentation and quantification of all individual organoids based on the open-source tools ImageJ/Fiji and Ilastik (**Fig. 1b**). Based on edge detection in each stack layer, this workflow can be used to robustly quantify organoid size (object area) and classify, e.g. by determining intensity, organoid phenotypes over time (**Fig. 1c, Supplementary Fig. S1b, S1c**). The workflow is robust to changes in morphology, stitching artefacts, and can be adjusted to image data from different automated microscopes. As the object classification by Ilastik is not dependent on deep learning and extensive training data, it can easily be adapted to new phenotypes and changes in experimental conditions.

**Figure 1:**
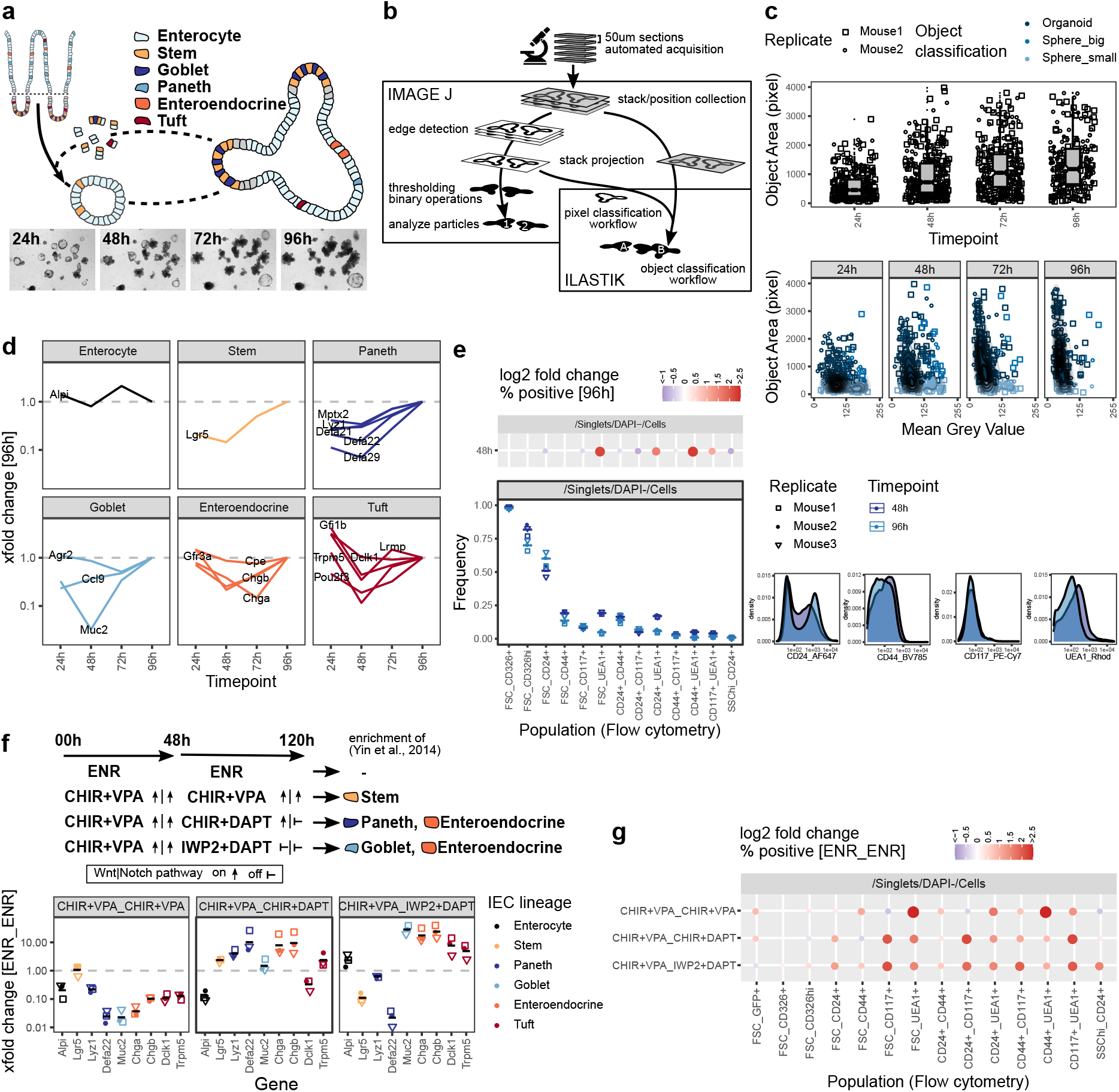
Quantification of intestinal organoid growth and cellular composition. a) Scheme of organoid formation and images of a representative position 24h-96h after seeding. Whole well is shown in **Supplementary Fig. S1a**. b) Scheme of organoid size quantification workflow using open-source tools ImageJ/Fiji and Ilastik. Visual quantification output and ImageJ quantification results are shown in **Supplementary Fig. S1b, S1c**. c) Box plots showing organoid size quantified as object area at 24-96h timepoints (top). Object area vs. object mean grey value (8-bit scale) on a minimum projection of the image stack (bottom). Pooled data from 2 biol replicates, indicated by shape. Each dot represents one organoid. d) mRNA expression of IEC lineage marker genes at 24h-96h timepoints, measured by qRT-PCR. xfold change relative to 96h organoids, median of 3 biol. replicates. e) Flow cytometry of organoids grown for 48h and 96h. Staining of representative replicate (bottom right). Population frequencies in Cells parent gate, 3 biol. replicates, indicated by shape. Mean highlighted (bottom left). Log2 fold change relative to 96h timepoint, median of 3 biol. replicates. Dot size corresponds to absolute log2 fold change (top left). Gating strategy and population frequencies for FSC CD326^hi^ and FSC CD24+ parent gates are shown in **Supplementary Fig. S1d, S1e**. f) Organoids cultured for 48h followed by 72h (48h 72h) with normal culture medium (ENR ENR), or culture medium containing CHIR+VPA CHIR+VPA, CHIR+VPA CHIR+DAPT, or CHIR+VPA IWP2+DAPT to modify IEC composition by interfering with Wnt and Notch signaling pathways as indicated, adapted from Yin et al. ^25^. mRNA expression measured by qRT-PCR. xfold change relative to ENR ENR treatment. 3 biol. replicates, indicated by shape. Mean highlighted. g) Flow cytometry of organoids treated for 48h 72h as indicated, population frequencies normalized to ENR ENR treatment. Log2 fold change, median of 3 biol. replicates. Dot size corresponds to absolute log2 fold change. Gating strategy, representative staining and population frequencies for FSC CD326^hi^ and FSC CD24+ parent gates are shown in **Supplementary Fig. S1d, S1f-S1h**.

In addition to determining organoid size, the cellular composition is of critical interest. We therefore selected transcripts that are specific for individual IEC lineages ^2^, and performed qRT-PCR on these within a 24h-96h time course (**Fig. 1d**). Except for the enterocyte marker *Alpi*, we generally find an increase in lineage-specific gene expression over time cumulating at 96h (**Fig. 1d**). This corresponds to the transition from spheroids, consisting mainly of progenitors, to mature budding organoids that contain more differentiated lineages, as was shown previously ^21^. As a complementary technique to quantify cellular composition on a single-cell level, we conducted flow cytometry of commonly used IEC surface markers (**Fig. 1e, Supplementary Fig. S1d, S1e**). The differences between 48h and 96h organoids were modest (**Fig. 1e**). Of note, we observed that the frequency of *Ulex europaeus* agglutinin 1 (UEA1) positive cells reduced over time, indicating that the population expressing UEA1 on the surface may be progenitor cells that are different from the population of UEA1^bright^ secretory cells commonly detected by immunofluorescence staining of permeabilized tissue (**Fig. 1e**). As a proof of principle, we next tested our approach on organoids with an altered cell composition. Interfering with WNT and NOTCH signaling pathways has previously been established by Yin and colleagues as a method to enrich organoids for stem cells, Paneth cells, goblet cells, or enteroendocrine cells ^25^. WNT and NOTCH pathways are activated or respectively inhibited by treatment with combinations of the glycogen synthase kinase 3 (GSK3) inhibitor CHIR99021 (CHIR), valproic acid (VPA), the porcupine inhibitor IWP2, or the gamma-secretase inhibitor DAPT ^25^ (**Fig. 1f**). Interestingly, we found that incubation with CHIR + VPA followed by IWP2 + DAPT increased the expression of tuft cell marker genes in addition to the previously described effects on goblet cells and enteroendocrine cells (**Fig. 1f**). Drastic effects on the cell composition were reflected by widely altered surface marker expression measured by flow cytometry and resulted in characteristic patterns (**Fig. 1g, Supplementary Fig. S1d, S1f, S1g**). However, we observed that well established flow cytometry gating strategies, such as identifying Paneth cells by a SSC^hi^ CD24+ gate ^26^, did not follow the *Lyz1* gene expression pattern in some conditions (**Fig. 1f, Supplementary Fig. S1h**). Thus, while flow cytometry demonstrates to be very useful to detect changes in the organoid composition, surface marker expression may be influenced by additional factors, such as the growth conditions, and identification of certain cell populations by flow cytometry requires appropriate controls. In summary, we developed an easy-to-use and cost-efficient toolbox for the analysis of (intestinal) organoids that is suitable to detect changes in organoid growth and cell composition.

### Organoid screen of epigenetic modifier inhibitors identifies established drugs targeting cancer growth

Next, we applied our organoid toolbox for screening of a chemical probe library that targets epigenetic modifiers to modulate the epigenome ^18^. Organoids generated from 4 individual mice were grown in the presence of 39 inhibitors, with DMSO vehicle and VPA serving as controls (Fig. 2a, 2b). Samples were imaged daily and expression of 12 transcripts specific for IEC lineages ^2^ was analyzed at the 96h endpoint (Fig. 2a). We observed that some of the probes significantly affected organoid growth as determined by area (Fig. 2c, 2d, Supplementary Fig. S2a-S2c, S3a). Integration of the primary readouts revealed a strong correlation of organoid size and expression of the ISC marker gene *Lgr5* (Fig. 2e). We found three probes that reduced both organoid size and *Lgr5* mRNA expression, namely the pan-Poly (ADP-ribose) polymerase (PARP) inhibitor olaparib and two histone deacetylase (HDAC) inhibitors LAQ824 (Dacinostat), a pan-HDAC inhibitor, and CI-994 (Tacedinaline), an HDAC1-3 & HDAC8 inhibitor (Fig. 2c-2e, Supplementary Fig. S2b, S2c) ^27,28,29^. These findings are in agreement with a study that showed reduced growth and *Lgr5* gene expression but a gain of enterocyte marker expression in CI-994 treated organoids ^30^. Olaparib-treated organoids would sufficiently grow to perform flow cytometry. This allowed us to use *Lgr5*-EGFP expressing reporter organoids to confirm the reduced *Lgr5* gene expression levels. Indeed, we found markedly fewer GFP-high/GFP-positive cells in olaparib-treated compared to control organoids (Fig. 2f, Supplementary Fig. S2d). Finally, we treated Adenomatous polyposis coli (*Apc*) knockout organoids, which are a model for intestinal cancer, with the two HDAC inhibitors and olaparib and found that these probes also limited growth in these tumor cultures, with similar growth reductions compared to WT organoids (Fig. 2g). Together, this is well in line with the design goal of these probes to limit cellular growth to target cancer cells.

**Figure 2:**
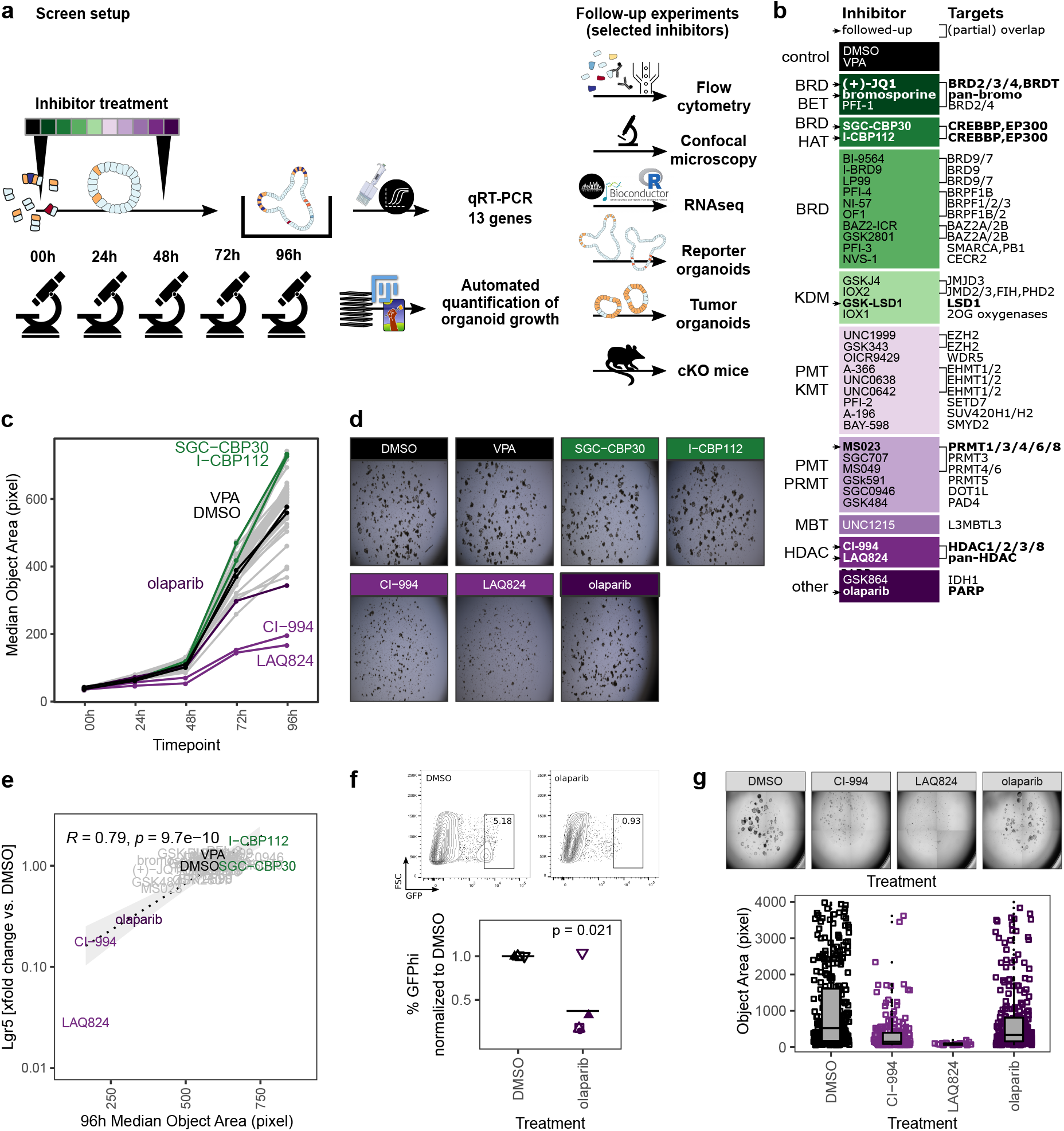
Organoid screen of epigenetics probes identifies established cancer drugs. a) Scheme of screen setup and follow-up experiments. b) Inhibitors used in screen, detailed information is provided in **Supplementary Table 1**. Probes used in follow-up experiments are highlighted. Abbreviations of inhibitor target families (inhibitor class): BRD - Bromodomain, BET - Extra-terminal motif, HAT - Histone acetyltransferase, KDM - Lysine demethylase, PMT - Protein methyltransferase, KMT - Lysine methyltransferase, PRMT - Protein arginine methyltransferase, MBT - Malignant brain tumor, HDAC - Histone deacetylase. c) Median object area of organoids treated with DMSO or inhibitors for 0-96h. Median of 4 biol. replicates. Boxplots for each inhibitor and timepoint are shown in **Supplementary Fig. S2a**. Probes that altered organoid size and were followed-up are highlighted: Controls, CI-994, LAQ824 (HDAC inhibitors), olaparib (PARP inhibitor), SGC-CBP30, I-CBP112 (BRD BET inhibitors). d) Representative replicates, 96h timepoint. e) Correlation of median organoid size and relative *Lgr5* gene expression, median of 4 biol. replicates. Pearson coefficient. f) Frequency of *Lgr5*-EGFP stem cells in reporter organoids treated with DMSO or olaparib for 96h. Gating of representative replicate (top) and percentage of GFP^hi^ cells, normalized to DMSO control. 5 biol. replicates, indicated by shape. Mean highlighted. Minimum 5000 viable cells in parent gate. Paired t-test (bottom). Percentage of total GFP+ cells is shown in **Supplementary Fig. S2d**. g) *Apc*-deficient adenomas treated with DMSO, CI-994, LAQ824, or olaparib for 96h. Representative replicate (top) and quantification of object size in 7/3/3/7 (DMSO/CI-994/LAQ828/olaparib) individual wells (bottom).

### Inhibition of EP300/CREBBP enhances organoid size and *Lgr5* expression

We next focused on probes that increased organoid size (**Fig. 2b-2d**). We found that both SGC-CBP30 and I-CBP112 significantly increased the organoid area (**Fig. S3a**) and this increase was seen in objects that were classified as “Organoids” and thus was not dependent on the occurrence of large spheres (**Fig. 3a, Supplementary Fig. S3b**). In support of the sensitivity of our assay, both probes have the same targets: EP300/CREBBP. E1A Binding Protein P300 (EP300, P300) and Creb-binding protein (CREBBP, CBP) are closely related bromodomain-containing acetyltransferases that serve as transcriptional co-activators for numerous transcription factors ^31,32,33^. Both SCG-CBP30 and I-CBP112 specifically target the bromodomain-binding domain, which thus renders these proteins unable to bind acetylated lysines. The observed increase in organoid size was surprising since both inhibitors have been designed to cause growth restriction in cancer cells ^34,35,36^. Comparing SGC-CBP30/I-CBP112-treated organoids with the DMSO vehicle control, organoid morphology appeared normal, however, we observed a reduction of putative goblet/Paneth cells as determined by cytosolic UEA1 staining (**Fig. 3b, Supplementary Fig. S3c**). We next tested whether these probes would expand the LGR5+ cell population in *Lgr5*-EGFP organoids and found a modest increase upon treatment, either alone or in combination with our positive control CHIR, an activator of canonical WNT signaling (**Fig. 3c, Supplementary Fig. S3d, S3e**). However, incubation with SGC-CBP30 or I-CBP112 could not enhance organoid growth under low EGF concentrations, replace R-Spondin in the culture medium, or overcome treatment with the WNT inhibitor IWP2 (**Supplementary Fig. S3f, S3g**). To determine which genes are under the control of EP300/CREBBP in the intestinal epithelium, we performed mRNA sequencing on untreated vs. I-CBP112 treated organoid cultures. In accordance with a transcriptional co-activator role for EP300/CREBBP, we found 53 genes upregulated and 110 genes downregulated using a log2 fold change cutoff of 0.5 and padjust *≤* 0.01 (**Fig. 3d**). Furthermore, signatures of transcription factors known to interact with either EP300 or CREBBP were negatively enriched (**Supplementary Fig. S3h**). Remarkably, *Lgr5* was the most significantly upregulated gene in our dataset, substantiating our previous results (**Fig. 3d**). In support, gene set enrichment analysis (GSEA) with a LGR5+ stem cell gene set ^37^ showed positive correlation (**Fig. 3e**). The second most significantly upregulated gene was *Egr1* (**Fig. 3d**), which is an inducible transcription factor that is involved in cell proliferation ^38^. The expansion of ISCs or progenitors appears to come at a cost to the differentiation of other cell lineages. We observed reduced UEA1 staining and downregulation of goblet cell markers such as *Muc4* and *Ccl6* following EP300/CREBBP inhibition (**Fig. 3b, 3d**). This is further supported by the negative correlation with secretory cell gene sets by GSEA (**Supplementary Fig. S3i**). Conversely, this is in line with positive enrichment of Gene Ontology biological process (GO:BP) terms such as smoothened signaling pathway and tissue morphogenesis (**Supplementary Fig. S3j, S3k**). Irrespective of the exact mechanism, we demonstrate that the paradoxical increase of organoid size after inhibition of EP300/CREBBP bromodomains may be explained by upregulation of *Lgr5*, *Egr1*, and genes associated with developmental processes, at the cost of IEC differentiation.

**Figure 3:**
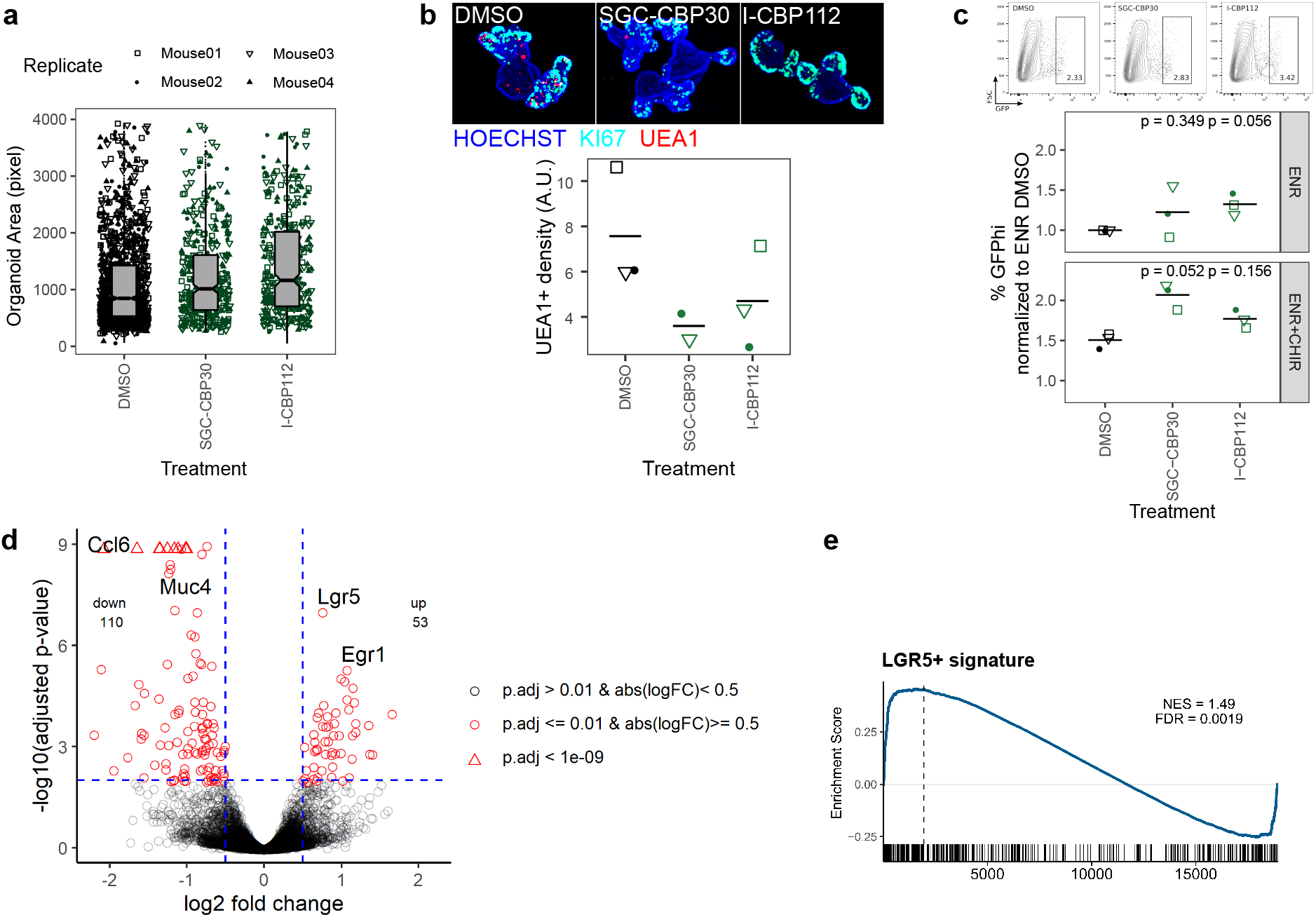
Inhibition of EP300/CREBBP enhances organoid size and *Lgr5* expression. a) Organoids treated with DMSO, SCG-CBP30, or I-CBP112 for 96h. Area of objects classified as “Organoid” by combined ImageJ/Ilastik workflow. 4 biol. replicates, indicated by shape. b) Representative organoids treated with DMSO, SCG-CBP30, or I-CBP112 for 96h. 10x magnification, max. intensity projection. KI67 staining marks crypt regions (top). Density of UEA1+ cells, each value represents the median of ≥5 organoids quantified. 3/2/3 biol. replicates, indicated by shape. Mean highlighted (bottom). Full wells for one representative replicate is shown in **Supplementary Fig. S3c**. c) Frequency of *Lgr5*-EGFP stem cells in reporter organoids grown in ENR or ENR+CHIR and treated with DMSO, SCG-CBP30, or I-CBP112 for 96h, measured by flow cytometry. Gating of representative replicate grown in ENR (top) and percentage of GFP^hi^ cells normalized to ENR DMSO condition of 3 biol. replicates, indicated by shape. Mean highlighted. Paired t-test (bottom). Percentage of total GFP+ cells is shown in **Supplementary Fig. S3d**. d) Volcano plot of mRNA sequencing of untreated vs. I-CBP112 treated organoids, 4 biol replicates per group. Selected genes are highlighted. e) mRNA sequencing of untreated vs. I-CBP112 treated organoids. GSEA of LGR5+ stem cell signature from Munñoz et al. ^37^ (GSE33949).

### GSK-LSD1 broadly affects IEC composition

So far, we have used organoid size as a probe selection criteria. Additionally, we performed qRT-PCR on 12 genes associated with specific cell lineages ^2^. We found that, after our positive control VPA, treatment with GSK-LSD1 leads to the largest perturbation of the IEC lineage marker profile as determined by calculating the euclidean distance of the gene expression xfold changes relative to DMSO treatment (**Fig. 4a, Supplementary Fig. S4a, S4f-S4k**). Flow cytometry screening of inhibitor treated organoids showed primarily moderate changes in surface marker expressions (**Supplementary Fig. S4b, S4c**). Although GSK484 and SGC0946 caused the most perturbation in surface marker populations, they showed little effect by qRT-PCR and thus we did not pursue these probes further (**Supplementary Fig. S4d, S4e**). Treatment with GSK-LSD1 markedly reduced gene expression of Paneth and goblet cell markers, but caused an increase in enteroendocrine and tuft cell marker genes, particularly *Gfi1b* (**Fig. 4b**). This supports our recent work in which we found that Lysine-specific Demethylase 1A (LSD1, KDM1A) is required for Paneth cell differentiation and contributes to goblet cell differentiation ^39,40^. Paneth cells are commonly gated as SSC^hi^ CD24+ population in flow cytometry ^26^. In line with a strong reduction of the Paneth cell marker genes *Lyz1* and *Defa22*, we find this population significantly reduced in GSK-LSD1 treated organoids (**Fig. 4b, 4c**). Furthermore, we observed that the pattern of CD24+ expressing cells in GSK-LSD1 treated organoids differs from control organoids in flow cytometry, with increase of a SSC^lo^ CD24^hi^ population (**Fig. 4c**). This pattern change was even more pronounced in SI crypt IECs from *Villin*-Cre+ *Lsd1* ^f/f^ mice, which conditionally lack *Lsd1* in IECs, compared to wild type (WT) littermates (**Fig. 4d**). Similar gating has previously been associated with enteroendocrine cells and their progenitors^26^, which thus fits with our previous observation that enteroendocrine progenitors such as *Neurod1* and *Neurog3* are upregulated in *Villin*-Cre+ *Lsd1* ^f/f^ mice ^39^. However, upon performing intracellular flow cytometry staining for the canonical tuft cell marker DCLK1, we found that also a DCLK1^hi^ population fell within this gate and is increased in *Lsd1*-deficient crypts (**Fig. 4e**). In support, there was a modest yet significant increase of DCLK1+ cells in duodenal tissue sections as well as colon sections from *Villin*-Cre+ *Lsd1* ^f/f^ mice compared to WT littermates (**Fig. 4f, Supplementary Fig. S4l**). Together, this example highlights that the epigenetic probe library contains inhibitors that are able to completely mimic the phenotype that is seen upon genetic deletion *in vivo*.

**Figure 4:**
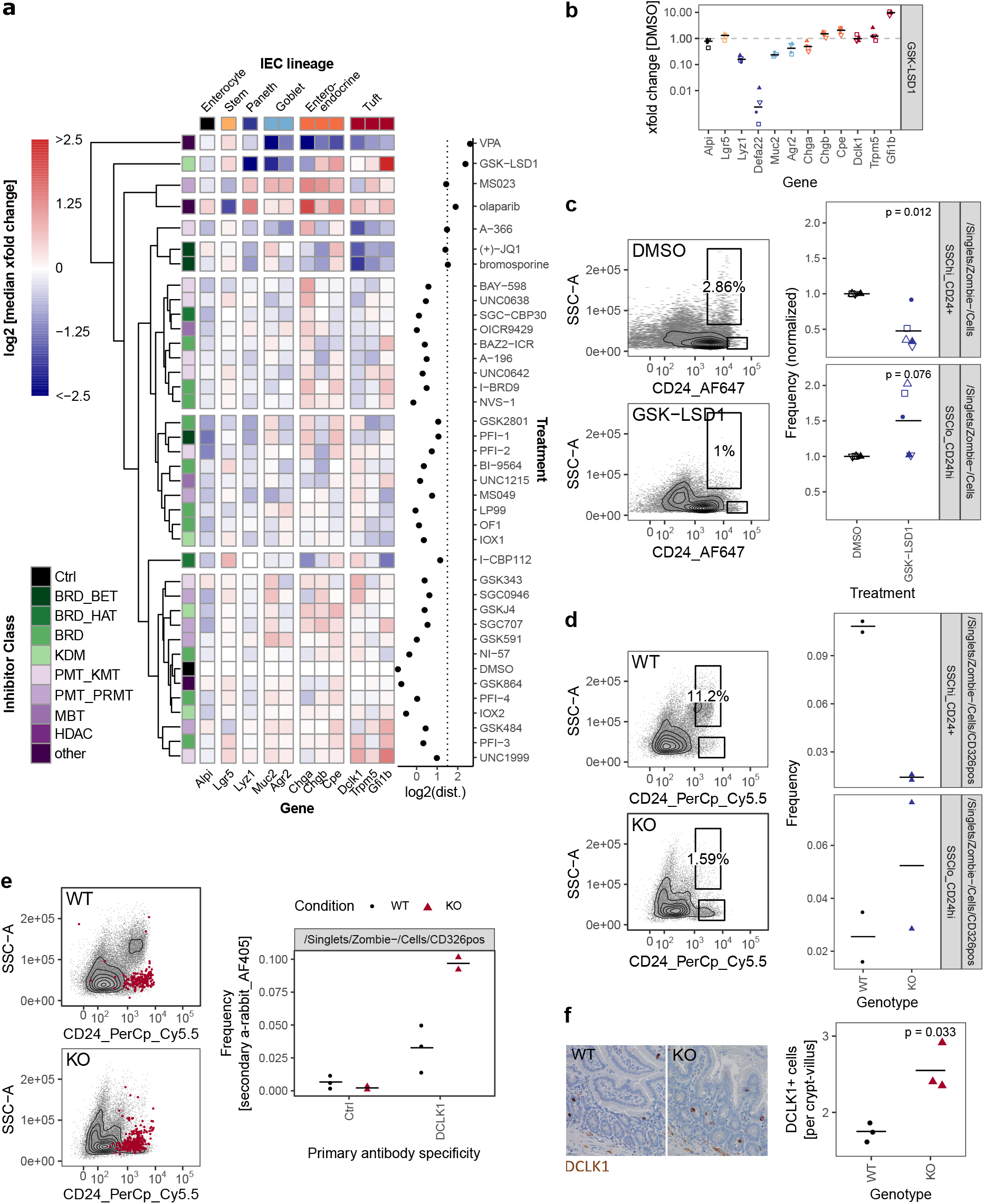
GSK-LSD1 broadly affects IEC composition. a) Gene expression of organoids treated with DMSO or inhibitors for 96h, measured by qRT-PCR. Color scale represents log2 of median of xfold change relative to DMSO-treated organoids of 4 biol. replicates. IEC lineage marker genes are indicated on x-axis. Inhibitor class is indicated on y-axis. Clustering tree is based on euclidean distance. Log2 of the euclidean distance (“perturbation”) is indicated in the right panel, the line at x=1.5 indicates inhibitors that were followed up in further experiments. Samples treated with HDAC inhibitors and gene *Defa22* and were excluded from the analysis. Euclidean distance including *Defa22* is shown in **Supplementary Fig. S4a**. b) Gene expression of organoids treated with GSK-LSD1 for 96h measured by qRT-PCR. xfold change relative to DMSO-treated organoids. 4 biol. replicates, indicated by shape. Median highlighted. c) Flow cytometry of organoids treated with DMSO or GSK-LSD1 for 96h. Gating of representative replicates (left) and normalized frequencies of SSC^hi^ CD24+ and SSC^lo^ CD24^hi^ populations of 5 biol. replicates, indicated by shape. Mean highlighted. Paired t-test (right). d) Flow cytometry of small intestinal crypts isolated from *Villin*-Cre+ *Lsd1* ^fl/fl^ (KO) mice with intestine-specific deletion of *Lsd1* or wild type (WT) littermates. Gating of representative replicates (left) and frequencies of SSC^hi^ CD24+ and SSC^lo^ CD24^hi^ populations of 2/2 mice (right). e) Frequency of DCLK1+ cells measured by intracellular flow cytometry in small intestinal crypts from WT and KO mice. Overlay of positive cells for secondary anti-rabbit staining, representative replicate (left). Quantification of intracellular staining with rabbit anti-DCLK1 primary antibody or control in small intestinal crypts isolated from WT or KO mice. 3/2 mice. Minimum 7000 viable cells in parent gate (right). DCLK1+ cells per crypt-villus pair in duodenum of WT and KO mice. Immunohistochemistry staining of tissue sections. Representative staining (left) and quantification in 3/3 mice, mean highlighted. Unpaired t-test (right).

### BET inhibition reduces relative abundance of tuft cells

Secretory cell lineage differentiation, such as goblet and Paneth cells, is well studied and is generally thought to involve NOTCH-mediated lateral inhibition. Tuft cell differentiation, however, is less defined. Therefore, we next focused on the BRD/BET inhibitors (+)-JQ1 and bromosporine in our marker gene expression dataset (**Fig. 4a**) as treatment with these led to a strong reduction of tuft cell marker genes *Dclk1*, *Trpm5*, and *Gfi1b* (**Fig. 5a**). Organoids treated with these probes also had altered expression in some of the other IEC lineage marker genes, but the downregulation of tuft cell marker genes was consistent and prominent (**Supplementary Fig. S5a**). Although probe A-366, an inhibitor of Euchromatic histone-lysine N-methyltransferase 1 and 2 (EHMT1/2, GLP/G9A) also reduced tuft cell marker genes, two other EHMT1/2 inhibitors, UNC0638 and UNC0642, did not (**Fig. 4a, Supplementary Fig. S4k**). (+)-JQ1 inhibits Bromodomain-containing protein 2 (BRD2), BRD3, BRD4, and BRDT while bromosporine is a pan-bromodomain inhibitor. In our hands, these two probes did not affect overall organoid growth in the 96h course of the screen experiment (**Supplementary Fig. S2a, S5b**), but (+)-JQ1 treatment affected organoid morphology when inhibitor treatment was continued after passaging (**Fig. 5b**). Others have reported that (+)-JQ1 treatment strongly reduced the efficiency of crypts to form organoids ^41^. Tuft cell quantification after treatment of *Hpgds2*-tdTomato reporter organoids indeed confirmed a complete lack of tuft cell differentiation in organoids treated with either (+)-JQ1 or bromosporine (**Fig. 5b, 5c**), suggesting that BRD proteins are necessary for the tuft cell lineage. BRD2, BRD3, BRD4, and BRDT are mutual targets of (+)-JQ1 and bromosporine, of which BRDT is not expressed in SI crytps or organoids (**Supplementary Fig. S5c**). Interestingly, the BRD2/4 inhibitor PFI-1 did not cause marked changes in tuft cell marker gene expression in our screen (**Supplementary Fig. S5d**). To investigate the role of specific BRDs in tuft cell differentiation in future studies may be worthwile.

**Figure 5:**
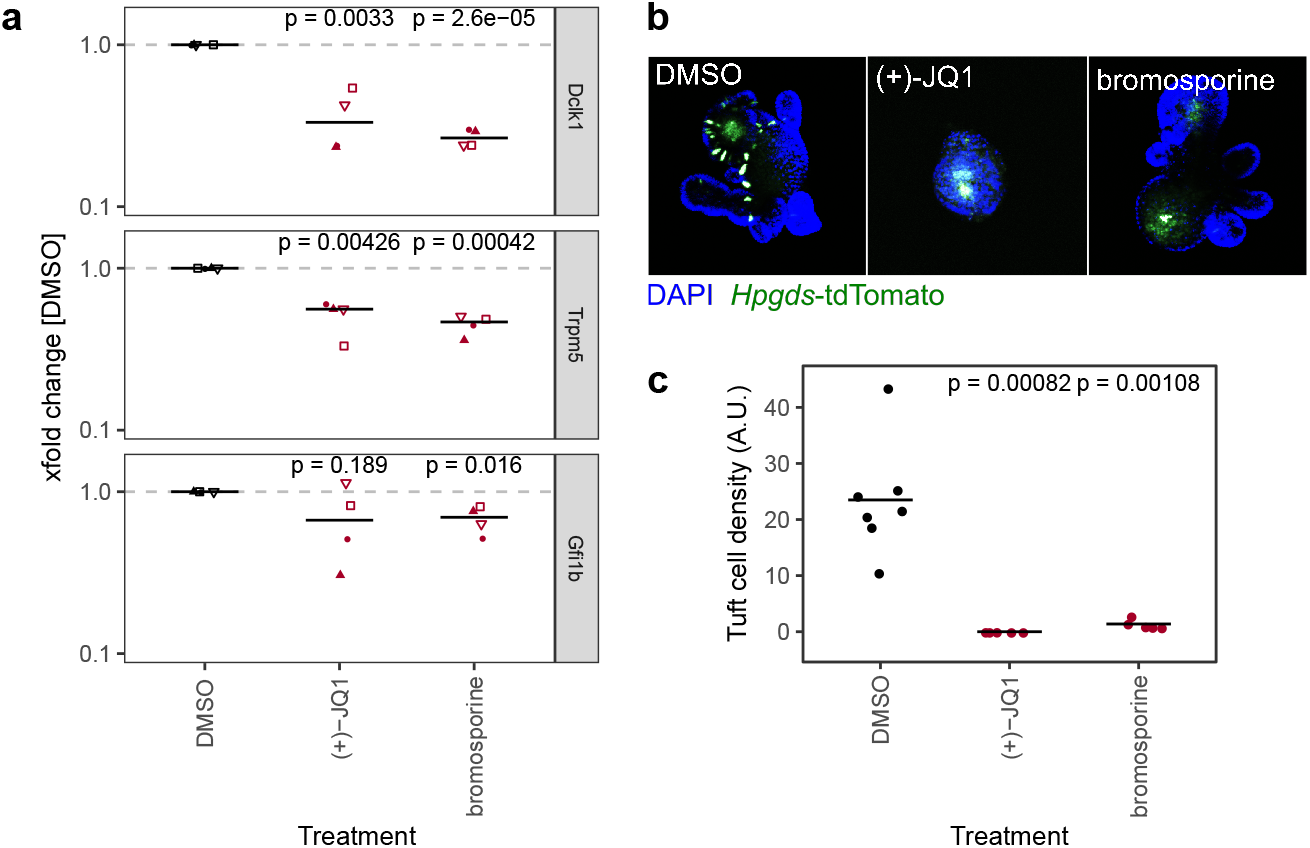
BET inhibition reduces relative abundance of tuft cells. a) Gene expression of tuft cell marker genes of organoids treated with DMSO, (+)-JQ1, or bromosporine for 96h, measured by qRT-PCR. xfold change relative to DMSO-treated organoids. 4 biol. replicates, indicated by shape. Median highlighted. b) Representative *Hpgds*-tdTomato tuft cell reporter organoids treated with DMSO, (+)-JQ1, or bromosporine for 8 days with one passage. 10x magnification, max. intensity projection. c) Tuft cell density in *Hpgds*-tdTomato organoids treated with DMSO, (+)-JQ1, or bromosporine for 8 days. Each dot represents one organoid.

### Inhibition of type I PRMTs results in higher relative abundance of secretory cells and prevents growth of tumor organoids

So far, we focused on inhibitors that caused reduced IEC differentiation. However, two probes stood out because they increased the expression of genes associated with Paneth-, goblet-, and enteroendocrine cells (**Fig. 4a**). Of these two, the pan-PARP inhibitor olaparib also had a marked effect on median organoid size and abundance of LGR5+ stem cells (**Fig. 2b-2e**). The other probe is MS023, which is an inhibitor of type 1 protein arginine methyltransferases (PRMTs) such as PRMT1, PRMT3, PRMT4 (CARM1), and PRMT8^42^ (**Fig. 6a**). Of note, two other PRMT inhibitors in our probe library, SGC707 and MS049 that inhibit PRMT3 and PRMT4/PRMT6 respectively, did not cause similar effects (**Supplementary Fig. 6a**). Although MS023-treated organoids were moderately yet signficantly smaller than control organoids and *Lgr5* gene expression was reduced, frequency of *Lgr5*-EGFP stem cells was not significantly affected, and organoids treated for 96h would renew normally after splitting (**Supplementary Fig. S6b-S6d**). The upregulation of secretory cell marker genes by the inhibitors was reflected by relative cell abundance of the respective lineages in MS023-treated versus control organoids. SSC^hi^ CD24+ Paneth cells appeared more frequent in MS023 treated organoids in our flow cytometry screen (**Fig. 6b, Supplementary Fig. S4c**), and quantification of MUC2+ goblet cells showed a trend in the same direction (**Fig. 6c**). Furthermore, we treated enteroendocrine cell reporter organoids with MS023 and found an increased frequency of *Neurog3*-RFP+ cells compared to the DMSO control (**Fig. 6d**). To get a more detailed overview of how MS023 affects organoids, we performed mRNA sequencing of untreated vs. MS023 treated organoids. We found 462 genes upregulated and 457 genes downregulated with a log2 fold change cutoff of 0.5 and padjust *≤* 0.01 (**Fig. 6e**). Importantly, GSEA of cell-lineage specific gene sets confirmed that MS023-treated organoids have a transcriptome that is enriched for secretory cell lineages (**Fig. 6f**). However, GSEA also indicated an enrichment for genes associated with enterocytes (**Fig. 6f**). Thus, rather than specifically affecting secretory cells, differentiation of all IEC cell lineages seems to be increased in MS023 treated organoids, potentially at the cost of progenitor cells. This is in agreement with positive enrichment of GO:BP terms related to nutrient uptake and response to microbials (**Supplementary Fig. S6e, S6f**), which are associated with mature enterocytes and Paneth cells respectively. DNA repair, which is a well established function of type I PRMTs ^43^, was among the negatively correlated GO:BP terms (**Supplementary Fig. S6e, S6g**). We found that PRMT1 was the type I PRMT with the highest gene expression level in SI crypts and organoids (**Supplementary Fig. S6h**). Enhanced PRMT levels are found in various malignancies and high PRMT1 expression is negatively correlated with survival in colon cancer ^44,45^. Furthermore, *Prmt1* gene expression was highest in ISC, transit-amplifying cells, and early enterocyte progenitors compared to fully differentiated lineages in a published IEC scRNA-seq dataset ^2^ (**Supplementary Fig. S6i**). We therefore hypothesize that inhibition of type I PRMTs leads to maturation of IECs, which aligns with the observation that differentiated cells have lower *Prmt1* levels. To test if PRMT type I inhibition could hence be used therapeutically to force progenitors, such as those found in WNT-driven tumors, to mature or differentiate, we treated *Apc*-deficient organoids with MS023 or the PRMT1-specific inhibitor TC-E5003. MS023 treated adenomas were smaller and darker than adenomas treated with DMSO control, and TC-E5003 treatment almost completely hindered their growth (**Fig. 6g, Supplementary Fig. S6j**). Yet, these probes did not cause growth inhibition of wild type organoids nor did they reduce cell viability (**Fig. 6h, Supplementary Fig. S6k, S6l**). In summary, we show that inhibition of type I PRMT leads to more differentiated organoids and has the potential to hinder proliferation in intestinal tumor organoids, making it an attractive candidate to pursue in future studies.

**Figure 6:**
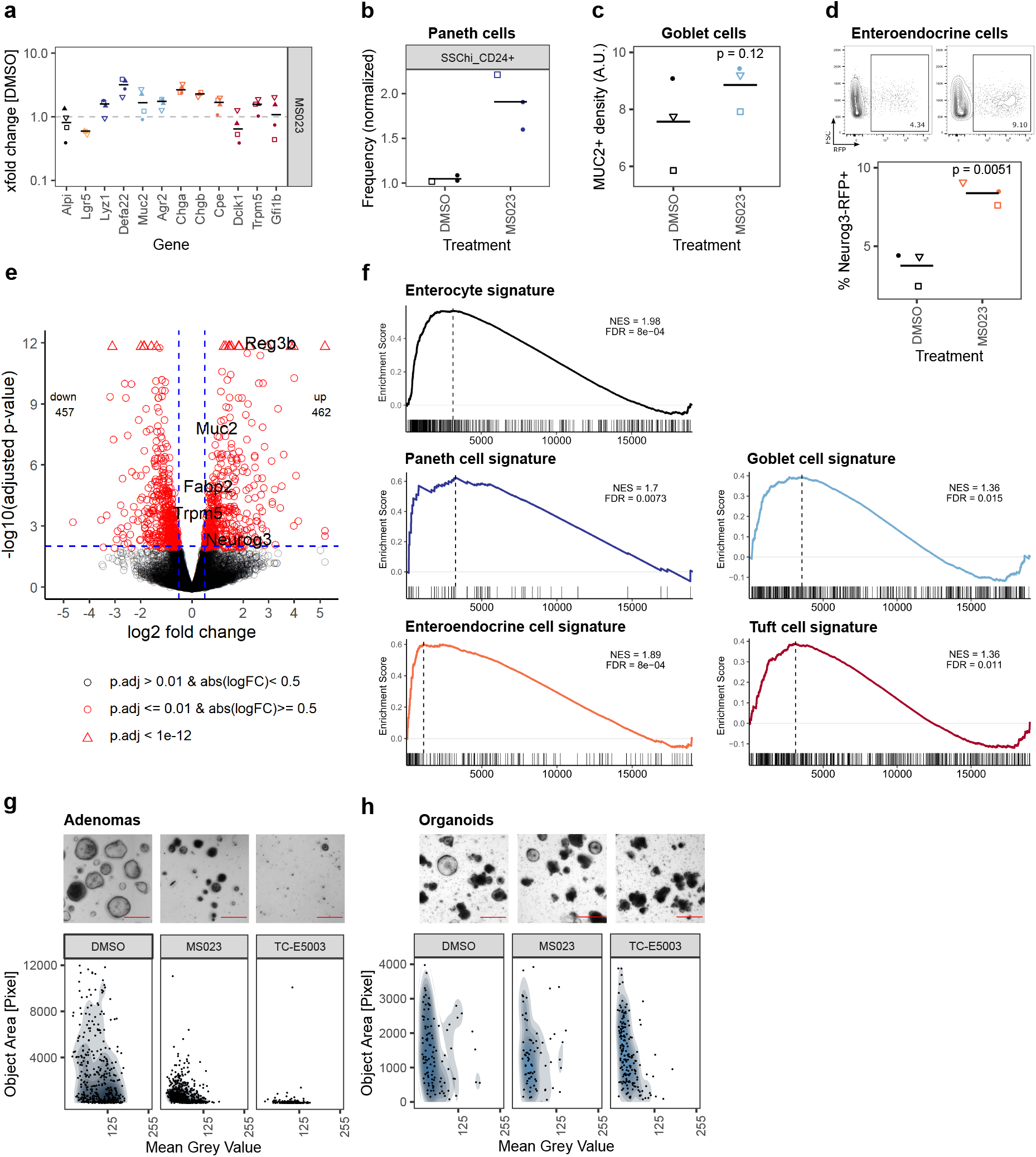
Inhibition of type I PRMTs leads to more mature organoids and prevents adenoma growth. a) Gene expression of organoids treated with MS023 for 96h measured by qRT-PCR. xfold change relative to DMSO-treated organoids. 4 biol. replicates, indicated by shape. Median highlighted. b) Frequency of SSC^hi^ CD24+ Paneth cells in organoids treated with MS023 for 96h, normalized to DMSO treatment, measured in flow cytometry screen (**Supplementary Fig. S4c**). 3 wells/2 biol. replicates, indicated by shape. Mean highlighted. c) Density of MUC2+ goblet cells in organoids treated with DMSO or MS023 for 96h. Median of 3 biol. replicates, indicated by shape. Each value is the median of 4-11 organoids quantified. Paired t-test. d) Frequency of enteroendocrine/enteroendocrine progenitor *Neurog3*-RFP+ cells in reporter organoids treated with DMSO or MS023 for 96h, measured by flow cytometry. Representative gating and quantification in 3 biol. replicates, indicated by shape. Paired t-test. e) Volcano plot of mRNA sequencing of untreated vs. MS023 treated organoids, 3 biol replicates per group. Selected genes are highlighted. f) mRNA sequencing of untreated vs. MS023 treated organoids. GSEA for Paneth cell, goblet cell, enteroendocrine cell, and tuft cell signatures from Haber et al. ^2^ (GSE92332). g) *Apc*-deficient adenomas treated with DMSO, MS023, or TC-E5003 for 96h. Representative well (top, full well shown in **Supplementary Fig. S6j**, scale bar shows 500*μ*m) and quantification of organoid size and mean grey value, 7 individual wells per condition (bottom). h) Organoids treated with DMSO, MS023, or TC-E5003 for 96h. Representative replicate (top, full well shown in **Supplementary Fig. S6k**, scale bar shows 500*μ*m) and quantification of organoid size and mean grey value, 3 biol. replicates per condition (bottom).

## Discussion

Working with heterogeneous organoid cultures is challenging with respect to reproducibility and quantification. Our toolkit, which we present in this article enables reproducible results across biological replicates using standard equipment and is suitable for screening setups. We established a quantification workflow that is based on the open source tools ImageJ and Ilastik, which is a simple yet robust alternative to recent stand-alone software options ^46,47^ and could easily be adapted to different tissue organoids. In addition, qRT-PCR and flow cytometry of IEC lineages is sufficiently sensitive for initial screening and was subsequently confirmed by additional methods such as reporter organoids. We used this screening setup to test a set of 39 chemical probes targeting epigenetic modifiers and identified probes that strongly affected organoid size or IEC lineage composition. These new regulators of intestinal epithelial biology are highly interesting candidates for further mechanistic studies.

Probes targeting EP300/CREBBP were designed as cancer therapeutics ^36^. Thus, we were surprised to find that inhibition of P300/CREBBP led to an increase of organoid size, which was supported by an expansion of LGR5+ cells and reduction of differentiation (**Fig. 3**). EP300/CREBBP mediate acetylation of histone H3K27 at enhancer elements and promoters, and can act as a transcriptional co-activator with numerous transcription factors ^31,32,48,33,49^. In support of a general activating role for EP300/CREBBP, we found that the majority of genes altered by I-CBP112-treatment were down-regulated (**Fig. 3d**), and many of these genes are established targets of EP300/CREBBP-associated transcription factors (**Fig. 3e**). EP300/CREBBP is a well established co-activator of signaling cascades that control cell proliferation, including WNT, NF*κ*B, or MYB signaling. Although we cannot rule out that altering these interactions might contribute to enhanced organoid growth, transcriptional signatures associated with *β*-Catenin (*Ctnnb1*), NF*κ*B-subunit RelA, or MYB were downregulated after treatment with the EP300/CREBBP inhibitor I-CBP112 (**Supplementary Fig. S3h**). It is difficult to separate the epigenetic modifier (H3K27 acetylation) from the transcriptional co-activator role of EP300/CREBBP, especially since a recent study showed a central role for the bromodomain and HAT domain also for the EP300/CREBBP transcription factor binding capacity ^50^. Nevertheless, underlining the critical role of the bromodomain, *plt6*-mice that carry a mutation in the EP300KIX domain, which specifically prevents interaction of EP300 with the transcription factor MYB, displayed reduced cell proliferation in the intestine ^51^. The EP300/CREBBP bromodomain is critically required for H3K27 acetylation at enhancer elements, a mark of active enhancers, and its inhibition leads to reduced expression of enhancer-proximal genes ^52,33^. In the adult small intestine, Sheaffer et al. described a gain of H3K27Ac at dynamically methylated enhancer sites in differentiated IECs but not LGR5+ ISC ^12^. Furthermore, Kazakevych et al. found that H3K27Ac positive distal elements were a good indicator for cell identity and differentiation status whereas genes positively regulating proliferation were transcribed in most IEC types ^5^. EP300/CREBBP has previously been shown to be required for differentiation of embryonic stem cells, muscle cells and adipocytes ^53,54,55^. In turn, Ebrahimi et al. recently described that EP300/CREBBP maintains transcription of fibroblast-specific somatic genes and that EP300/CREBBP bromodomain inhibition can promote cellular reprogramming to pluripotency ^56^, accompanied by decrease in promoter- and enhancer-associated H3K27 acetylation ^56^. We provide evidence that EP300/CREBBP inhibition in the intestinal epithelium can promote proliferation rather than preventing it. It appears plausible that EP300/CREBBP bromodomain activity is critically required to enable transcription of IEC differentiation genes and that in its absence the intestinal epithelium remains immature, accompanied by an enhanced proliferative capacity.

We recently demonstrated a central role of LSD1 in Paneth and goblet cell differentiation and maturation ^39,40^. Here, we confirm the critical role of LSD1 for IEC lineage differentiation in an unbiased screen and in addition provide indications that instead of Paneth/goblet cells there is an expansion of DCLK1+ tuft cells that is associated with a CD24^high^ SSC^low^ population by flow cytometry (**Fig. 4**). In contrast, we find that treatment with the BRD/BET inhibitors (+)JQ-1 and bromosporine completely blocks tuft cell differentiation (**Fig. 5**). Tuft cells are important mediators of intestinal type 2 immunity ^57,58^. Our work matches observations of two studies that found that inhibition of the BET bromodomain *in vivo* abolished tuft cells ^41,59^. Using a different BRD/BET probe, Nakagawa et al. described that the absence of tuft cells was due to blockade of transit-amplifying cells as their intermediate progenitors ^59^. While Nakagawa et al. also found a reduction of enteroendocrine cells, another study described an increase of pancreatic NEUROG3+ enteroendocrine progenitors following (+)JQ-1 treatment ^60^. Our findings could be the foundation of using these compounds to modulate immune responses, especially when a type 2 response is unfavourable.

Type I PRMT inhibition with MS023 led to a more differentiated intestinal epithelium without major loss of LGR5+ stem cells (**Fig. 6, Supplementary Fig. S6b**). PRMT1 was the most highly expressed type I PRMT and is higher expressed in ISCs and progenitors compared to differentiated cells (**Supplementary Fig. S6h, S6i**). An evolutionary conserved role of PRMT1 in the adult intestine has been proposed earlier as endogenous PRMT1 knockdown reduces the adult ISC population in *Xenopus* and zebrafish, while transgenic PRMT1 overexpression leads to an increase of ISCs ^61,62^. Furthermore, our observation is very similar to findings of enriched PRMT1 in epidermis progenitors, required for maintenance of this population ^63^. Bao et al. proposed that PRMT1 is both involved in the maintenance of progenitor/proliferative genes as well as the repression of ‘differentiation’ genes ^63^. In agreement with the latter, we found increase of all differentiated IEC lineages after treatment with MS023 (**Fig. 6a-6f**). PRMT1 has a wide substrate specificity and mediates both arginine methylation of histones such as H4R3, and non-histone proteins ^64,65^. Elevated PRMT1 expression is found in several cancer types and is associated with poor prognosis and chemoinsensitivity ^66,67^ and pharmacological PRMT inhibitors have recently gained interest as drug candidates for cancer treatment ^43,45^. Targeting cancer stem cells (CSCs) in the gut comes with the challenge that following ablation of LGR5+ CSCs, LGR5-cells have the potential to de-differentiate to CSCs ^68^. Therefore, forcing differentiation of cancer cells could be an attractive treatment strategy. Indeed, we found that PRMT type I inhibition with MS023 and PRMT1-specific inhibition with TC-E5003 severely impaired growth of *Apc*-deficient tumor organoids but not normal organoids (**Fig. 6g, 6h**). A conditional *Prmt1*-deficient mouse was recently generated ^69^. Crossing these mice with intestine-specific *Villin*-Cre or tumor-developing *Apc*^min^ mice, could be an elegant way to further study the role of PRMT1 in IEC differentiation and maturation and to investigate the therapeutic potential of PRMT1 inhibition for the treatment of intestinal cancer.

Highly permissive chromatin and transcriptional control of IEC fate, as well as gene regulation by differential chromatin states have been discussed as opposing models of intestinal epithelial differentiation regulation ^10^. Testing a library of highly selective inhibitors targeting more than 20 epigenetic modification enzymes/enzyme families, only two HDAC-inhibitors prevented organoid growth (**Fig. 2**) and the majority of the tested probes did not alter organoid growth or IEC composition. However, we found that few compounds resulted in pronounced changes and these were associated with generally less (EP300/CREBBP, LSD1 inhibition) or more (PRMT type I inhibition) epithelial differentiation. We therefore propose that epigenetic modifiers control the degree of intestinal epithelial differentiation in general, rather than affecting specific cell lineage fate. Whether this parallels the postnatal maturation of the fetal intestinal epithelium remains to be elucidated. Of note, the epigenetic modifiers identified to affect IEC differentiation in our screen share the capacity to both modify histones and to interact with multiple transcription factors. Thus, these molecules could be a key link connecting the epigenetic and the transcriptional layers of gene regulation in the intestinal epithelium. Indeed, work by others supports a model of tightly intertwined epigenetic and transcriptional control and shifting between permissive and dynamic chromatin on a local instead of a global scale. By integrating the investigation of gene expression, open chromatin, and DNA hydroxymethylation in IEC populations with differential expression levels of the transcription factor SOX9, recent elegant work by Raab et al. identified either highly permissive or dynamic chromatin states at given loci relative to transcription factor binding ^70^. EP300 has been described to potentiate SOX9-dependent transcription ^71^ and *Sox9*-deficient intestinal epithelium fails to mature ^72^. Mapping of EP300-binding sites was recently utilized to identify transcriptional networks in specialized cell types in the placenta ^73^, inspiring further investigation of epigenetic modifier-aided transcription in different IEC lineages.

To summarize, we developed a resource that allows to compare the requirement of various epigenetic modifiers for intestinal epithelial renewal and IEC differentiation. Our results indicate that some epigenetic modifiers with the capacity to both mediate histone modifications and act as transcriptional co-regulators control the balance between an undifferentiated/differentiated epithelial state. Thereby, they lay basis for a fine-tuned transcriptional regulation and rapid adjustment upon injury or pathogenic challenge.

## Methods

### Epigenetic modifier inhibitors

The epigenetic modifier inhibitors in the screen experiment were part of the Structural Genomics Consortium Epigenetic Chemical Probes Collection as of March 2016. Probes were reconstituted in DMSO and used at the recommended concentration as listed in Supplementary Table 1. 1mM valproic acid (VPA) was included as positive control. DMSO vehicle control was matched to the highest concentration used per experiment, maximal 10*μ*M. PRMT1-specific inhibitor TC-E5003 (Santa Cruz Biotechnology, # sc397056) was included in follow-up experiments and used at 50*μ*M, equivalent to 10*μ*M DMSO.

### Mice

C57BL/6JRj wild type (Janvier labs), *Lgr5*-EGFP-IRES-CreERT2 (Jackson Laboratories, stock no: 008875), *Villin*-Cre ^74^ (kind gift from Sylvie Robine), *Lsd1* ^f/f^ ^75^ (kind gift from Stuart Orkin), and *Apc*^15lox^ (Jackson Laboratories, stock no: 029275) mice were housed under specific-pathogen free conditions at the Comparative Medicine Core Facility (CoMed), Norwegian University of Science and Technology, Norway. For the flow cytometry screening experiment, organoids were generated from C57BL/6 mice housed at the Leibniz-Forschungsinstitut fuÖr Molekulare Pharmakologie, Germany. *Hpgds*-tdTomato mice ^76^ were housed at University of Montpellier, France. *Neurog3*-RFP mice ^77^ (kind gift from Anne Grapin-Botton) were housed at University of Copenhagen, Denmark. Experiments were performed following the respective legislation on animal protection, were approved by the local governmental animal care committee, and were in accordance with the European Convention for the Protection of Vertebrate Animals used for Experimental and other Scientific purposes.

### Small intestinal crypt isolation

Small intestinal crypts were isolated as described previously ^78^. The proximal half of the small intestine was rinsed, opened longitudinally, cut to small pieces after villi and mucus were scraped off, washed with PBS until the solution was clear, and incubated in 2mM EDTA/PBS for 30min at 4^*◦*^C with gentle rocking. Fragments were subsequently washed with PBS and the crypt fraction was typically collected from wash 2-5. All centrifugation steps were carried out at 300 *×* g.

### Organoid culture

Organoids were generated by seeding ca. 250-500 small intestinal crypts in a 50*μ*l droplet of cold Matrigel (Corning #734-1101) into the middle of a pre-warmed 24-well plate. Matrigel was solidified by incubation at 37^*◦*^C for 5-15min and 500*μ*l culture medium added. Basal culture medium (“ENR”) consisted of advanced DMEM F12 (Gibco) supplemented with 1x Penicillin-Streptomycin (Sigma-Aldrich), 10mM HEPES, 2mM Glutamax, 1x B-27 supplement, 1x N2 supplement, (all Gibco) 500mM N-Acetylcysteine (Sigma-Aldrich), 50ng/ml recombinant EGF (Thermo Fisher Scientific), 10% conditioned medium from a cell line producing Noggin (kind gift from Hans Clevers), and 20% conditioned medium from a cell line producing R-Spondin-1 (kind gift from Calvin Kuo). ENR culture medium was replaced every 2-3 days. Organoids were passaged at 1:3-1:4 ratio by disruption with rigorous pipetting almost to single cells. Organoid fragments were centrifuged at 300 *×* g, resuspended in 40-50*μ*l cold Matrigel per well, and plated on pre-warmed 24-well plates. Organoids derived from different mice or a repetition at least one passage apart are considered biological replicates. Technical replicates, i.e. separate wells, were carried out in some experiments and were pooled for analysis.

### Altering IEC lineage composition in organoids

Protocols to alter the IEC composition in organoids have been described previously ^25,79,80^. Organoids were grown for 48h in ENR or ENR + 3*μ*M CHIR99021 (Sigma-Aldrich) and 1mM valproic acid (VPA). Then, media was replaced by ENR, ENR + 3*μ*M CHIR and 1mM VPA, ENR + 3*μ*M CHIR and 10*μ*M DAPT, or ENR + 10*μ*M DAPT and 2*μ*M IWP2. VPA, DAPT, IWP2 were purchased from Cayman Chemicals. Organoids were harvested 72h after media change.

### Organoid screen with Epigenetic Chemical Probes library

Organoids of four biological replicates were passaged to nearly single cells at 1:4 ratio as described above and seeded in 40*μ*l Matrigel droplets in 24-well plates. 250*μ*l/well ENR were added immediately after solidification and 250*μ*l/well ENR + probes at 2x working concentration (see Supplementary Table 1) were added within 30min. For each biological replicate DMSO vehicle controls were carried out in quadruplicates. Media was replaced after 48h. Organoids bright-field images were acquired daily on an EVOS2 microscope and after 96h RNA was harvested.

### Reporter organoids

*Lgr5*-EGFP organoids were generated as described above from *Lgr5*-EGFP-IRES-CreERT2 mice and maintained for no longer than 3 weeks. Organoids were grown in ENR or ENR + 3*μ*M CHIR99021 (Sigma-Aldrich) as indicated and *Lgr5*-EGFP+ cells were quantified using a BD LSRII flow cytometer (Becton Dickinson) as percentage of viable cells. Tuft cell reporter organoids were generated from *Hpgds*-tdTomato mice (expressing tdTomato under the *Hpgds* promoter) as described above. *Hpgds*-tdTomato+ cells were quantified by confocal microscopy on an Axio Imager Z1 microscope (Zeiss) as number of cells relative to the organoid area after z-stack projection, determined by nuclear staining. Enteroendocrine cell reporter organoids were derived from the proximal small intestine of *Neurog3*-RFP mice (expressing RFP under the *Neurog3* promoter) and cultured as described above using recombinant murine Noggin (100ng/ml, Peprotech) and 10% R-Spondin conditioned medium. *Neurog3*-RFP+ cells were quantified using a BD FACSAria III flow cytometer (Becton Dickinson) as percentage of viable cells.

### Modified organoid growth conditions with EP300/CREBBP inhibition

*Lgr5*-EGFP organoids were grown in ENR or ENR + 3*μ*M CHIR for 96h. Media was replaced after 48h. To investigate low growth factor conditions, wild type organoids were grown in ENR or ENR with 1% R-Spondin, ENR with 5ng/ml EGF, or ENR + 2*μ*M IWP2 for 192h. Media was replaced every 48h.

### Splitting of organoids after type I PRMT inhibition

Organoids were treated with DMSO or MS023 for 96h, passaged to nearly single cells as described above, and cultured in ENR for additional 96h. Media was replaced every 48h.

### Generation of APC-deficient adenomas

Eight week old *Apc*^15lox^ *× Lgr5*-EGFP-IRES-CreERT2 mice were administered 2mg Tamoxifen in corn oil (both Sigma-Aldrich) for 5 consecutive days. Adenomatous polyps developed over the course of a month (ethically approved by the Norwegian Food Safety Authority, FOTS ID: 15888). To generate adenoma organoids, the small intestine was rinsed with PBS, opened longitudinally, polyps were excised, cut into small pieces, and washed in PBS. Next, 5 ml TrypLE express (Thermo Fisher Scientific) was added and incubated for 30min at 37^*◦*^C while pipetting every 5-10min. After incubation, single cells were obtained by passing the supernatant through a 40*μ*m strainer. Single cells were plated in 50*μ*l cold Matrigel on a pre-warmed 24-well plate, and cultured in basal culture medium lacking R-Spondin-1 (“EN”). EN culture medium was replaced every 2-3 days.

### Organoid growth quantification

Organoid bright-field images were acquired on an EVOS2 microscope (Thermo Fisher Scientific) with 2x magnification. At the starting point of the experiment, for each plate an automation setup was generated to acquire z-stacks with 50*μ*m spacing either of a single position or 2-4 tiled images covering height and most area of the Matrigel dome for each well. This automation setup was reused at consecutive timepoints. A custom ImageJ/Fiji v1.52n ^81,82^ macro was used to collect single positions and layers for each well, to save a stack (ImageJ bright-field stack) and projections, and to perform a simple organoid segmentation (“ImageJ workflow”). For the segmentation, a Sobel edge detector was applied to each z-stack layer (ImageJ edge stack), a standard deviation z-projection of the edge stack was generated, and particle analysis with optional manual correction was performed after several binary operations and thresholding. For an improved segmentation that is robust to stitching artefacts, allows to filter out debris and organoid clusters and to distinguish different organoid phenotypes, the ImageJ workflow was combined with the interactive machine learning software Ilastik v1.3.2^83^ (“combined workflow”). Training data was taken from the analyzed experiment and excluded from further analysis. In a first step, pixel classification on an intensity summary projection of the ImageJ edge stack was used to separate between background and object outlines. The generated pixel prediction maps were then used as input in a second step of object classification together with minimum projections of the ImageJ bright-field stack. Routinely, the following label classes were used: Organoid, big sphere, small sphere, cluster, debris, background mislabelled as organoid, air bubble, edges of well plate. Objects classfied in the latter three object classes were excluded from all timepoints, objects classfied as debris or cluster were excluded from 72h and 96h timepoints. Representative images were arranged using GNU R packages magick and ggimage.

### RNA isolation, quantitative RT-PCR and analysis

To harvest RNA, organoids in the Matrigel dome were dissolved in 250*μ*l RNA-solv reagent (Omega Bio-Tek). RNA was isolated using Direct-zol-96 RNA or Direct-zol MiniPrep kit (Zymo Research) according to the manufacturer’s instructions, including DNAse digestion. cDNA was transcribed using High-Capacity RNA-to-cDNA Kit (Applied Biosystems) according to the manufacturer’s instructions. RNA quality and concentration was assessed on an NanoDrop-1000 instrument (NanoDrop). Samples were handled in 96-well plates and transferred with multichannel pipettes. qRT-PCR was carried out in technical duplicates in 384-well plates on a QuantStudio 5 instrument (Thermo Fisher Scientific) using 2x Perfecta ROX,UNG Fast Mix (Quanta Biosciences) and 5ng cDNA per reaction in a total volume of 12*μ*l. Primer-probe combinations were selected based on the Universal Probe Library System (Roche) and are listed in Supplementary table 2, primers were purchased from Sigma-Aldrich. *Hprt* was used as housekeeping gene. ∆CT values were calculated as ∆CT = CT(housekeeping gene) - CT(gene of interest) (such that higher values indicate higher relative expression); ∆∆CT values referred to the calibrator as indicated, and fold change was calculated as 2^∆∆CT^. Target gene “perturbation” was calculated as euclidean distance of the log2 median fold change using GNU R package pheatmap. *Defa22* gene expression was below the detection limit for some samples and was therefore omitted from the euclidean distance ranking but is provided in the supplementary information.

### Flow cytometry

To obtain single cells, Matrigel in 1-3 wells was disrupted by pipetting, well content was transferred to an Eppendorf tube, centrifuged at 300× g, and supernatant removed. Then, organoids were incubated with 300*μ*l TrypLE express (Thermo Fisher Scientific) for 37^*◦*^C for 30min and pipetted up/down with a 1000*μ* pipet tip prior to and after the incubation. Single cells were stained with Zombie Aqua (Biolegend, 1:1000 in PBS) for 15min at room temperature (RT) for live-dead exclusion. If DAPI instead of Zombie Aqua staining was used for live-dead exclusion, it was added it during the last washing step (1:1000). Samples were incubated with antibody conjugates against CD326-BV605, CD24-PerCp-Cy5.5 or AF647, CD44-BV785, CD117-PE-Cy7 (all Biolegend, see Supplementary Table 3 for detailed list, 1:200 in PBS + 2% fetal calf serum (FCS)), and Ulex Europaeus Agglutinin (UEA)1-Rhodamine (2*μ*g/ml, Vector Laboratories #RL-1062-2) for 20min at 4^*◦*^C. For intracellular staining, samples were subsequently fixed with 2% paraformaldehyde (PFA) for 15min, and incubated with or without rabbit anti-DCLK1 (Abcam #ab31704, 1:500 in PBS/2% FCS/0.05% Saponin) for 1h at 4^*◦*^C, followed by incubation with Goat anti-Rabbit IgG-AF405 (Invitrogen, 1:1000 in PBS + 2% FCS + 0.05% Saponin). Samples were analyzed on a BD LSRII instrument (Becton Dickinson) equipped with 405nm, 488nm, 561nm, 647nm laser lines. Single fluorochrome stainings of cells and compensation particles (BD CompBead, Becton Dickinson) were included in each experiment. For analysis, FlowJo software v10.6.2 and GNU R/Bioconductor v3.6.3/v3.10 packages flowCore, CytoML/flowWorkspace, ggcyto, flowViz were used ^84^. If not indicated otherwise, only samples with more than 10000 viable cells in the parent gate were included.

### Flow cytometry screening

For the flow cytometry screening experiment, organoids were grown in 96well glass-bottom plates (Cellvis) that were pre-cooled and held on ice during seeding. Organoid fragments in Matrigel (50*μ*l/well) were distributed in pre-cooled plates with an automated pipette, then plates were transferred to a rotary plate shaker for 30sec at 150rpm, before the Matrigel was solidified at 37 ^*◦*^C. With help of a Viaflo 96-channel pipette (Integra Biosciences) 200*μ*l/well ENR without or with inhibitors were added of which 100*μ*l were replaced daily during the 96h time course. To obtain single cells, culture media was removed, 100*μ*l/well TrypLE express (Thermo Fisher Scientific) added and Matrigel disrupted by repeated pipetting with a multichannel pipette. Staining with Zombie Aqua, CD326-BV421, CD24-PerCp-Cy5.5, CD44-AF647, CD117-PE-Cy7 (all Biolegend, see Supplementary Table 3 for detailed list), and UEA1-FITC (Invitrogen) was carried out as described above. Samples were run on a MACSQuant X instrument (Miltenyi Biotec) equipped with 405nm, 488nm, 647nm laser lines and analyzed as described above. Euclidean distance clustering tree of normalized median population frequencies was generated with GNU R package ggtree ^85^.

### Confocal microscopy and quantification

For immunofluorescence staining, organoids were grown in 30*μ*l/well Matrigel droplets in a 8-well microscopy chamber (Ibidi) that was pre-warmed for seeding. After 96h incubation, the organoids were fixed in 4% paraformaldehyde and 2% sucrose for 30min at RT, washed, and permeabilized with 0.2% Triton-X100 in PBS. Free aldehyde groups were blocked using 100mM glycine, followed by blocking buffer (1% BSA, 2% NGS diluted in 0.2% Triton-X100 in PBS) for 1h at RT. The organoids were incubated overnight at 4^*◦*^C with a primary antibody against KI67 (Invitrogen #MA5-14520; 1:200) or MUC2 (Santa Cruz #sc-15334; 1:200) in blocking buffer, followed by three washes with slight agitation. Next, the organoids were incubated with Goat anti-Rabbit IgG-AF488 (Invitrogen, 1:500), UEA1-Rhodamine (Vector Laboratories #RL-1062-2, 2*μ*g/ml), and Hoechst 33342 overnight at 4^*◦*^C. After washing, the organoids were mounted using Fluoromount G (Thermo Fisher Scientific), and visualized using a LSM880 confocal microscope (Zeiss). UEA1/MUC2-positive cells were manually counted for *≥*5 organoids per biological replicate in a middle plane of a z-stack. Cell numbers are reported relative to the area of the z-stack projection of each organoid, determined by nuclear staining.

### Immunohistochemistry & Immunofluorescence staining of paraffin-embedded tissue and quantification

Immediately after euthanizing mice, the intestinal tissues were removed, washed with PBS, fixed in 4% formaldehyde for 48-72h at RT, and embedded in paraffin wax. Staining was carried out on 4*μ*m paraffin sections. The sections were rehydrated and treated with 3% hydrogen peroxide for 10min at RT. Antigens were retrieved by boiling the slides in citrate buffer (pH6) in a microwave for 15min. For immunohistochemistry staining of duodenum sections, the sections were incubated overnight at 4^*◦*^C with anti-DCLK1 (Abcam #ab31704; 1:1500) in TBS + 0.025% Tween 20 + 1% BSA. Specific binding was detected with Envision-HRP (Dako) and DAB (Dako) and images were acquired on a EVOS2 microscope (Thermo Fisher Scientific) with 10x magnification. DCLK1+ cells were quantified for *≥*30 crypt-villus pairs per mouse. Representative images were acquired on a Eclipse Ci-L microscope (Nikon) with 20x magnification. For immunofluorescence staining of colon sections, slides were blocked with PBS + 1% BSA + 2% goat serum + 0.2% Triton X-100 for 1h at RT and incubated overnight at 4^*◦*^C with anti-DCLK1 antibody (Abcam #ab31704; 1:250) in PBS + 1% BSA +1% goat serum + 0.05% Tween Specific binding was detected with Goat anti-Rabbit IgG-AF488 (Invitrogen, 1:1000) for 1h at 37 ^*◦*^C while nuclei were stained with DAPI (1:1000). Slides were mounted with Fluoromount G (Thermo Fisher Scientific) and images were acquired on a LSM880 confocal microscope (Zeiss) with 20x magnification. DCLK1+ cells were quantified for *≥*50 crypts per mouse.

### mRNA sequencing

Organoid RNA was isolated and treated with DNAse with Quick-RNA Micro prep kit (Zymo Research) according to manufacturer’s instructions. RNA integrity numbers were found to be *≥*7. For the I-CBP112 inhibitor study, library preparation was done using the Illumina TruSeq Stranded protocol. Library concentrations were quantified with the Qubit Fluorometric Quantitation system (Life Technologies) and the size distribution was assessed using a 2100 Bioanalyzer automated electrophoresis system (Agilent). For sequencing, samples were diluted and pooled into NGS libraries in equimolar amounts and sequenced at 75bp single-read chemistry on an Illumina NS500 MO flow-cell on a Ilumina NextSeq 500 instrument (Illumina) by the Genomics core facility (GCF, NTNU). For the MS023 study, library preparation was done using the NEB Next Ultra RNA Library Prep Kit with poly(A) mRNA enrichment and samples were sequenced at 150X2 bp paired-end chemistry on a Illumina NovaSeq 6000 instrument by Novogene (UK) Co.

### mRNA sequencing analysis

Read quality was assessed using FastQC v0.11.8, reads were aligned with STAR v2.7.3a to the Mus musculus genome build mm10, and MultiQC v1.7 was used to summarize logs from STAR and FastQC ^86,87,88^. The number of reads that uniquely aligned to the exon region of each gene in GENCODE annotation M18 of the mouse genome was then counted using featureCounts v1.6.4^89,90^. Genes that had a total count less than 10 were filtered out. Differential expression was then determined with GNU R/Bioconductor v3.6.1/v3.10 package DESeq2 v1.26.0 using default settings and shrunken log2foldchange was calculated with the apeglm method ^91,92^. GSEA enrichment was performed using GNU R/Bioconductor v3.6.3/v3.10 package ClusterProfiler v3.14.3 by shrunken log2 fold change and with the shrunken log2 fold change as weights using 10000 permutations ^93^. Gensets for celltype signatures were assembled based on single-cell and bulk RNA-Sequencing data from sorted samples based on datasets by Haber et al. ^2^ (GSE92332) and Munoz et al. ^37^ (GSE33949). Transcription factors interacting with murine or human EP300 or CREBBP were retrieved from protein-protein interactions with an minimum medium experimental confidence level (≥0.4) from STRING-DB v11^94^. Genesets regulated for the mouse and human version of these transcription factors were retrieved from TRRUST v2^95^. For human genesets, murine orthologue genes retrieved from Ensembl GRCh38.p13 through GNU R/Bioconductor v3.6.3/v3.10 package biomaRt v2.42.1^96^ were used for enrichment. Genesets for characterization of Biological Process were directly obtained from the Gene Ontology Consortium ^97^.

### Data processing and statistical analysis

Data was processed and statistical analysis was carried out with GNU R v3.6.3 using the packages tidyverse and ggpubr ^98^. Pearson correlation coefficient, and paired or unpaired t-test were calculated as indicated, assuming normal distribution. Median or mean are shown as indicated. In boxplots, the box represent the 25%, 50% and 75% percentiles and whiskers represent 1.5 *×* IQR.

## Data availability

The Imaging data from the initial screen was deposited to the Image Data Resource ^99^ (https://idr.openmicroscopy.org) under accession number idr0092. The Ilastik projects and respective training data of the initial screen organoid segmentation were deposited to Zenodo under https://doi.org/10.5281/zenodo.4311473. The qRT-PCR data from the initial screen was deposited along with processed data from follow-up experiments to BioStudies database at EMBL-EBI ^100^ (https://www.ebi.ac.uk/biostudies) under accession number S-BSST447. RNA-seq data were deposited in the ArrayExpress database at EMBL-EBI ^101^ (https://www.ebi.ac.uk/arrayexpress) under accession number E-MTAB-9290 (I-CBP112-treated samples) and E-MTAB-9291 (MS023-treated samples).

## Code availability

The ImageJ script used for organoid segmentation is available from https://github.com/jennyostrop/Fiji_organoid_brightfield_processing and deposited under https://doi.org/10.5281/zenodo.3951126.

## Acknowledgements

We thank the imaging (CMIC), animal care (CoMed), and genomics core (GCF) facilities at NTNU for assistance in this work. Funding of this work was provided by the Norwegian Research Council (‘Young Research Talent’ 274760 to MJO and Centre of Excellence grant 223255/F50) and the Norwegian Cancer Society (182767 to MJO). The flow cytometry screen was carried out at Leibniz-Forschungsinstitut för Molekulare Pharmakologie (FMP) Screening Unit, Berlin, Germany with funding from H2020 CORBEL (PID6031 to JO). ELIXIR Norway and the Centre for Digital Life Norway, funded by the Norwegian Research Council (270068 and 248810/O70) assisted with data management. We thank Dr. Sneha Pinto for critical reading of the manuscript.

## Author contributions

JO, MJO designed the study. JO, RZ, MTP, FG, HTL, AD, NP, SR, MJO performed experiments. JO, RZ, MTP, FG, KB, HTL analyzed data. JO, KB, HTL curated data. CA provided critical materials. PJ, KBJ, CA, MJO supervised or provided critical insight. JO, RZ, MJO wrote the manuscript with subsequent input from other authors.

## Supplementary information

Supplementary Figures S1-S6

Supplementary Table 1: Inhibitors and concentrations used

Supplementary Table 2: qRT-PCR primers and probes used

Supplementary Table 3: Materials, reagents, and software used

## References

1. Helmuth Gehart and Hans Clevers. Tales from the crypt: New insights into intestinal stem cells. Nat Rev Gastroenterol Hepatol, 16(1):19–34, January 2019. ISSN 1759-5053. doi: 10.1038/s41575-018-0081-y.

2. Adam L. Haber, Moshe Biton, Noga Rogel, Rebecca H. Herbst, Karthik Shekhar, Christopher Smillie, Grace Burgin, Toni M. Delorey, Michael R. Howitt, Yarden Katz, Itay Tirosh, Semir Beyaz, Danielle Dionne, Mei Zhang, Raktima Raychowdhury, Wendy S. Garrett, Orit Rozenblatt-Rosen, Hai Ning Shi, Omer Yilmaz, Ramnik J. Xavier, and Aviv Regev. A single-cell survey of the small intestinal epithelium. Nature, 551 (7680):333–339, November 2017. ISSN 0028-0836, 1476-4687. doi: 10.1038/nature24489.

3. Unmesh Jadhav, Madhurima Saxena, Nicholas K. O’Neill, Assieh Saadatpour, Guo-Cheng Yuan, Zachary Herbert, Kazutaka Murata, and Ramesh A. Shivdasani. Dynamic Reorganization of Chromatin Accessibility Signatures during Dedifferentiation of Secretory Precursors into Lgr5+ Intestinal Stem Cells. Cell Stem Cell, 21(1):65–77.e5, July 2017. ISSN 1875-9777. doi: 10.1016/j.stem.2017.05.001.

4. Qi Yang, Nessan A. Bermingham, Milton J. Finegold, and Huda Y. Zoghbi. Requirement of Math1 for Secretory Cell Lineage Commitment in the Mouse Intestine. Science, 294(5549):2155–2158, December 2001. ISSN 0036-8075, 1095-9203. doi: 10.1126/science.1065718.

5. Juri Kazakevych, Sergi Sayols, Berith Messner, Christina Krienke, and Natalia Soshnikova. Dynamic changes in chromatin states during specification and differentiation of adult intestinal stem cells. Nucleic Acids Res, 45(10):5770–5784, June 2017. ISSN 0305-1048. doi: 10.1093/nar/gkx167.

6. Unmesh Jadhav, Alessia Cavazza, Kushal K. Banerjee, Huafeng Xie, Nicholas K. O’Neill, Veronica Saenz-Vash, Zachary Herbert, Shariq Madha, Stuart H. Orkin, Huili Zhai, and Ramesh A. Shivdasani. Extensive Recovery of Embryonic Enhancer and Gene Memory Stored in Hypomethylated Enhancer DNA. Molecular Cell, 74(3):542–554.e5, May 2019. ISSN 1097-2765. doi: 10.1016/j.molcel.2019.02.024.

7. Greeshma Ray and Michelle S. Longworth. Epigenetics, DNA Organization, and Inflammatory Bowel Disease. Inflamm. Bowel Dis., 25(2):235–247, January 2019. ISSN 1536-4844. doi: 10.1093/ibd/izy330.

8. Yaser Atlasi and Hendrik G. Stunnenberg. The interplay of epigenetic marks during stem cell differentiation and development. Nat. Rev. Genet., 18(11):643–658, November 2017. ISSN 1471-0064. doi: 10.1038/nrg.2017.57.

9. Damiana Álvarez-Errico, Roser Vento-Tormo, Michael Sieweke, and Esteban Ballestar. Epigenetic control of myeloid cell differentiation, identity and function. Nat. Rev. Immunol., 15(1):7–17, January 2015. ISSN 1474-1741. doi: 10.1038/nri3777.

10. Ellen N. Elliott and Klaus H. Kaestner. Epigenetic regulation of the intestinal epithelium. Cell. Mol. Life Sci., 72(21):4139–4156, November 2015. ISSN 1420-9071. doi: 10.1007/s00018-015-1997-9.

11. Tae-Hee Kim, Fugen Li, Isabel Ferreiro-Neira, Li-Lun Ho, Annouck Luyten, Kodandaramireddy Nalapareddy, Henry Long, Michael Verzi, and Ramesh A. Shivdasani. Broadly permissive intestinal chromatin underlies lateral inhibition and cell plasticity. Nature, 506(7489):511–515, February 2014. ISSN 1476-4687. doi: 10.1038/nature12903.

12. Karyn L. Sheaffer, Rinho Kim, Reina Aoki, Ellen N. Elliott, Jonathan Schug, Lukas Burger, Dirk Schübeler, and Klaus H. Kaestner. DNA methylation is required for the control of stem cell differentiation in the small intestine. Genes Dev., 28(6):652–664, March 2014. ISSN 1549-5477. doi: 10.1101/gad.230318.113.

13. Lucas TJ Kaaij, Marc van de Wetering, Fang Fang, Benjamin Decato, Antoine Molaro, Harmen JG van de Werken, Johan H. van Es, Jurian Schuijers, Elzo de Wit, Wouter de Laat, Gregory J. Hannon, Hans C. Clevers, Andrew D. Smith, and René F. Ketting. DNA methylation dynamics during intestinal stem cell differentiation reveals enhancers driving gene expression in the villus. Genome Biology, 14(5):R50, May 2013. ISSN 1474-760X. doi: 10.1186/gb-2013-14-5-r50.

14. Cheryl H. Arrowsmith, Chas Bountra, Paul V. Fish, Kevin Lee, and Matthieu Schapira. Epigenetic protein families: A new frontier for drug discovery. Nature Reviews Drug Discovery, 11(5):384–400, May 2012. ISSN 1474-1784. doi: 10.1038/nrd3674.

15. Martijn A. J. Koppens, Gergana Bounova, Gaetano Gargiulo, Ellen Tanger, Hans Janssen, Paulien Cornelissen-Steijger, Marleen Blom, Ji-Ying Song, Lodewyk F. A. Wessels, and Maarten van Lohuizen. Deletion of Polycomb Repressive Complex 2 From Mouse Intestine Causes Loss of Stem Cells. Gastroenterology, 151(4):684–697.e12, October 2016. ISSN 0016-5085. doi: 10.1053/j.gastro.2016.06.020.

16. Alexis Gonneaud, Naomie Turgeon, François Boudreau, Nathalie Perreault, Nathalie Rivard, and Claude Asselin. Distinct Roles for Intestinal Epithelial Cell-Specific Hdac1 and Hdac2 in the Regulation of Murine Intestinal Homeostasis. J. Cell. Physiol., 231(2):436–448, February 2016. ISSN 1097-4652. doi: 10.1002/jcp.25090.

17. Judith Kraiczy and Matthias Zilbauer. Intestinal Epithelial Organoids as Tools to Study Epigenetics in Gut Health and Disease, 2019.

18. Sebastian Scheer, Suzanne Ackloo, Tiago S. Medina, Matthieu Schapira, Fengling Li, Jennifer A. Ward, Andrew M. Lewis, Jeffrey P. Northrop, Paul L. Richardson, H. Ümit Kaniskan, Yudao Shen, Jing Liu, David Smil, David McLeod, Carlos A. Zepeda-Velazquez, Minkui Luo, Jian Jin, Dalia Barsyte-Lovejoy, Kilian V. M. Huber, Daniel D. De Carvalho,Masoud Vedadi, Colby Zaph, Peter J. Brown, and Cheryl H. Arrowsmith. A chemical biology toolbox to study protein methyltransferases and epigenetic signaling. Nat Commun, 10(1):1–14, January 2019. ISSN 2041-1723. doi: 10.1038/s41467-018-07905-4.

19. Suzanne Ackloo, Peter J. Brown, and Susanne Müller. Chemical probes targeting epigenetic proteins: Applications beyond oncology. Epigenetics, 12(5):378–400, May 2017. ISSN 1559-2308. doi: 10.1080/15592294.2017.1279371.

20. Meritxell Huch, Juergen A. Knoblich, Matthias P. Lutolf, Alfonso Martinez-Arias. The hope and the hype of organoid research. Development, 144(6):938–941, March 2017. ISSN 0950-1991, 1477-9129. doi: 10.1242/dev.150201.

21. Denise Serra, Urs Mayr, Andrea Boni, Ilya Lukonin, Markus Rempfler, Ludivine Challet Meylan, Michael B. Stadler, Petr Strnad, Panagiotis Papasaikas, Dario Vischi, Annick Waldt, Guglielmo Roma, and Prisca Liberali. Self-organization and symmetry breaking in intestinal organoid development. Nature, 569(7754): 66–72, May 2019. ISSN 1476-4687. doi: 10.1038/s41586-019-1146-y.

22. Moshe Biton, Adam L. Haber, Noga Rogel, Grace Burgin, Semir Beyaz, Alexandra Schnell, Orr Ashenberg, Chien-Wen Su, Christopher Smillie, Karthik Shekhar, Zuojia Chen, Chuan Wu, Jose Ordovas-Montanes, David Alvarez, Rebecca H. Herbst, Mei Zhang, Itay Tirosh, Danielle Dionne, Lan T. Nguyen, Michael E. Xifaras, Alex K. Shalek, Ulrich H. von Andrian, Daniel B. Graham, Orit Rozenblatt-Rosen, Hai Ning Shi, Vijay Kuchroo, Omer H. Yilmaz, Aviv Regev, and Ramnik J. Xavier. T Helper Cell Cytokines Modulate Intestinal Stem Cell Renewal and Differentiation. Cell, 175(5):1307–1320.e22, November 2018. ISSN 0092-8674. doi: 10.1016/j.cell.2018.10.008.

23. Nathalie Brandenberg, Sylke Hoehnel, Fabien Kuttler, Krisztian Homicsko, Camilla Ceroni, Till Ringel, Nikolce Gjorevski, Gerald Schwank, George Coukos, Gerardo Turcatti, and Matthias P. Lutolf. High-throughput automated organoid culture via stem-cell aggregation in microcavity arrays. Nature Biomedical Engineering, pages 1–12, June 2020. ISSN 2157-846X. doi: 10.1038/s41551-020-0565-2.

24. Toshiro Sato, Robert G. Vries, Hugo J. Snippert, Marc van de Wetering, Nick Barker, Daniel E. Stange, Johan H. van Es, Arie Abo, Pekka Kujala, Peter J. Peters, and Hans Clevers. Single Lgr5 stem cells build crypt-villus structures in vitro without a mesenchymal niche. Nature, 459(7244):262–265, May 2009. ISSN 1476-4687. doi: 10.1038/nature07935.

25. Xiaolei Yin, Henner F. Farin, Johan H. van Es, Hans Clevers, Robert Langer, and Jeffrey M. Karp. Niche-independent high-purity cultures of Lgr5 + intestinal stem cells and their progeny. Nature Methods, 11(1): 106–112, January 2014. ISSN 1548-7105. doi: 10.1038/nmeth.2737.

26. Toshiro Sato, Johan H. van Es, Hugo J. Snippert, Daniel E. Stange, Robert G. Vries, Maaike van den Born, Nick Barker, Noah F. Shroyer, Marc van de Wetering, and Hans Clevers. Paneth cells constitute the niche for Lgr5 stem cells in intestinal crypts. Nature, 469(7330):415–418, January 2011. ISSN 1476-4687. doi: 10.1038/nature09637.

27. J. Mateo, C. J. Lord, V. Serra, A. Tutt, J. Balmaña, M. Castroviejo-Bermejo, C. Cruz, A. Oaknin, S. B. Kaye, and J. S. de Bono. A decade of clinical development of PARP inhibitors in perspective. Annals of Oncology, 30(9):1437–1447, September 2019. ISSN 0923-7534. doi: 10.1093/annonc/mdz192.

28. E. Weisberg, L. Catley, J. Kujawa, P. Atadja, S. Remiszewski, P. Fuerst, C. Cavazza, K. Anderson, and J. D. Griffin. Histone deacetylase inhibitor NVP-LAQ824 has significant activity against myeloid leukemia cells in vitro and in vivo. Leukemia, 18(12):1951–1963, December 2004. ISSN 1476-5551. doi: 10.1038/sj.leu.2403519.

29. Thomas Beckers, Carmen Burkhardt, Heike Wieland, Petra Gimmnich, Thomas Ciossek, Thomas Maier, and Karl Sanders. Distinct pharmacological properties of second generation HDAC inhibitors with the benzamide or hydroxamate head group. International Journal of Cancer, 121(5):1138–1148, 2007. ISSN 1097-0215. doi: 10.1002/ijc.22751.

30. Alexis Gonneaud, Christine Jones, Naomie Turgeon, Dominique Lévesque, Claude Asselin, Françcois Boudreau, and François-Michel Boisvert. A SILAC-Based Method for Quantitative Proteomic Analysis of Intestinal Organoids. Scientific Reports, 6(1):38195, November 2016. ISSN 2045-2322. doi: 10.1038/srep38195.

31. Richard H. Goodman and Sarah Smolik. CBP/p300 in cell growth, transformation, and development. Genes Dev., 14(13):1553–1577, July 2000. ISSN 0890-9369, 1549-5477. doi: 10.1101/gad.14.13.1553.

32. Yolande F. M. Ramos, Matthew S. Hestand, Matty Verlaan, Elise Krabbendam, Yavuz Ariyurek, Michiel van Galen, Hans van Dam, Gert-Jan B. van Ommen, Johan T. den Dunnen, Alt Zantema, and Peter A. C. ^’^t Hoen. Genome-wide assessment of differential roles for p300 and CBP in transcription regulation. Nucleic Acids Res, 38(16):5396–5408, September 2010. ISSN 0305-1048. doi: 10.1093/nar/gkq184.

33. Ryan Raisner, Samir Kharbanda, Lingyan Jin, Edwin Jeng, Emily Chan, Mark Merchant, Peter M. Haverty, Russell Bainer, Tommy Cheung, David Arnott, E. Megan Flynn, F. Anthony Romero, Steven Magnuson, and Karen E. Gascoigne. Enhancer Activity Requires CBP/P300 Bromodomain-Dependent Histone H3K27 Acetylation. Cell Reports, 24(7):1722–1729, August 2018. ISSN 2211-1247. doi: 10.1016/j.celrep.2018.07.041.

34. Duncan A. Hay, Oleg Fedorov, Sarah Martin, Dean C. Singleton, Cynthia Tallant, Christopher Wells, Sarah Picaud, Martin Philpott, Octovia P. Monteiro, Catherine M. Rogers, Stuart J. Conway, Timothy P. C. Rooney, Anthony Tumber, Clarence Yapp, Panagis Filippakopoulos, Mark E. Bunnage, Susanne Müller, Stefan Knapp, Christopher J. Schofield, and Paul E. Brennan. Discovery and Optimization of Small-Molecule Ligands for the CBP/p300 Bromodomains. J. Am. Chem. Soc., 136(26):9308–9319, July 2014. ISSN 0002-7863. doi: 10.1021/ja412434f.

35. Sarah Picaud, Oleg Fedorov, Angeliki Thanasopoulou, Katharina Leonards, Katherine Jones, Julia Meier, Heidi Olzscha, Octovia Monteiro, Sarah Martin, Martin Philpott, Anthony Tumber, Panagis Filippakopoulos, Clarence Yapp, Christopher Wells, Ka Hing Che, Andrew Bannister, Samuel Robson, Umesh Kumar, Nigel Parr, Kevin Lee, Dave Lugo, Philip Jeffrey, Simon Taylor, Matteo L. Vecellio, Chas Bountra, Paul E. Brennan, Alison O’Mahony, Sharlene Velichko, Susanne Müller, Duncan Hay, Danette L. Daniels, Marjeta Urh, Nicholas B. La Thangue, Tony Kouzarides, Rab Prinjha, Jürg Schwaller, and Stefan Knapp. Generation of a selective small molecule inhibitor of the CBP/p300 bromodomain for leukemia therapy. Cancer Res, 75(23):5106–5119, December 2015. ISSN 0008-5472. doi: 10.1158/0008-5472.CAN-15-0236.

36. Narsis Attar and Siavash K. Kurdistani. Exploitation of EP300 and CREBBP Lysine Acetyltransferases by Cancer. Cold Spring Harb Perspect Med, 7(3), March 2017. ISSN 2157-1422. doi: 10.1101/cshperspect.a026534.

37. Javier Muñoz, Daniel E. Stange, Arnout G. Schepers, Marc van de Wetering, Bon-Kyoung Koo, Shalev Itzkovitz, Richard Volckmann, Kevin S. Kung, Jan Koster, Sorina Radulescu, Kevin Myant, Rogier Versteeg, Owen J. Sansom, Johan H. van Es, Nick Barker, Alexander van Oudenaarden, Shabaz Mohammed, Albert J. R. Heck, and Hans Clevers. The Lgr5 intestinal stem cell signature: Robust expression of proposed quiescent ‘+4’ cell markers. EMBO J., 31(14):3079–3091, June 2012. ISSN 1460-2075. doi: 10.1038/emboj.2012.166.

38. Delphine Gitenay and Véronique T. Baron. Is EGR1 a potential target for prostate cancer therapy? Future Oncol, 5(7):993–1003, September 2009. ISSN 1744-8301. doi: 10.2217/fon.09.67.

39. Rosalie T. Zwiggelaar, Havard T. Lindholm, Madeleine Fosslie, Marianne Terndrup Pedersen, Yuki Ohta, Alberto Díez-Sáanchez, Mara Martín-Alonso, Jenny Ostrop, Mami Matano, Naveen Parmar, Emilie Kvaløy, Roos R. Spanjers, Kamran Nazmi, Morten Rye, Finn Drabløs, Cheryl Arrowsmith, John Arne Dahl, Kim B. Jensen, Toshiro Sato, and Menno J. Oudhoff. LSD1 represses a neonatal/reparative gene program in adult intestinal epithelium. Science Advances, 6(37):eabc0367, September 2020. ISSN 2375-2548. doi: 10.1126/sciadv.abc0367.

40. Naveen Parmar, Kyle Burrows, Håvard T. Lindholm, Rosalie T. Zwiggelaar, Mara Martín-Alonso, Madeleine Fosslie, Bruce Vallance, John Arne Dahl, Colby Zaph, and Menno J. Oudhoff. Intestinal-epithelial LSD1 controls cytoskeletal-mediated cell identity including goblet cell effector responses required for gut inflammatory and infectious diseases. bioRxiv, page 2020.07.09.186114, July 2020. doi: 10.1101/2020.07.09.186114.

41. Jessica E. Bolden, Nilgun Tasdemir, Lukas E. Dow, Johan H. van Es, John E. Wilkinson, Zhen Zhao, Hans Clevers, and Scott W. Lowe. Inducible In Vivo Silencing of Brd4 Identifies Potential Toxicities of Sustained BET Protein Inhibition. Cell Reports, 8(6):1919–1929, September 2014. ISSN 2211-1247. doi: 10.1016/j.celrep.2014.08.025.

42. Mohammad S. Eram, Yudao Shen, Magdalena Szewczyk, Hong Wu, Guillermo Senisterra, Fengling Li, Kyle V. Butler, H. Ümit Kaniskan, Brandon A. Speed, Carlo Dela Seña, Aiping Dong, Hong Zeng, Matthieu Schapira, Peter J. Brown, Cheryl H. Arrowsmith, Dalia Barsyte-Lovejoy, Jing Liu, Masoud Vedadi, and Jian Jin. A Potent, Selective, and Cell-Active Inhibitor of Human Type I Protein Arginine Methyltransferases. ACS Chem. Biol., 11(3):772–781, March 2016. ISSN 1554-8937. doi: 10.1021/acschembio.5b00839.

43. Ernesto Guccione and Stéphane Richard. The regulation, functions and clinical relevance of arginine methylation. Nat. Rev. Mol. Cell Biol., 20(10):642–657, October 2019. ISSN 1471-0080. doi: 10.1038/s41580-019-0155-x.

44. K. Mathioudaki, A. Papadokostopoulou, A. Scorilas, D. Xynopoulos, N. Agnanti, and M. Talieri. The PRMT1 gene expression pattern in colon cancer. British Journal of Cancer, 99(12):2094–2099, December 2008. ISSN 1532-1827. doi: 10.1038/sj.bjc.6604807.

45. James Jarrold and Clare C. Davies. PRMTs and Arginine Methylation: Cancer’s Best-Kept Secret? Trends in Molecular Medicine, 25(11):993–1009, November 2019. ISSN 1471-4914. doi: 10.1016/j.molmed.2019.05.007.

46. Timothy Kassis, Victor Hernandez-Gordillo, Ronit Langer, and Linda G. Griffith. OrgaQuant: Human Intestinal Organoid Localization and Quantification Using Deep Convolutional Neural Networks. Sci Rep, 9(1):12479, August 2019. ISSN 2045-2322. doi: 10.1038/s41598-019-48874-y.

47. Michael A. Borten, Sameer S. Bajikar, Nobuo Sasaki, Hans Clevers, and Kevin A. Janes. Automated brightfield morphometry of 3D organoid populations by OrganoSeg. Sci Rep, 8(1):5319, March 2018. ISSN 2045-2322. doi: 10.1038/s41598-017-18815-8.

48. Feng Wang, Christopher B. Marshall, and Mitsuhiko Ikura. Transcriptional/epigenetic regulator CBP/p300 in tumorigenesis: Structural and functional versatility in target recognition. Cell. Mol. Life Sci., 70(21): 3989–4008, November 2013. ISSN 1420-9071. doi: 10.1007/s00018-012-1254-4.

49. Brian T. Weinert, Takeo Narita, Shankha Satpathy, Balaji Srinivasan, Bogi K. Hansen, Christian Schölz, William B. Hamilton, Beth E. Zucconi, Wesley W. Wang, Wenshe R. Liu, Joshua M. Brickman, Edward A. Kesicki, Albert Lai, Kenneth D. Bromberg, Philip A. Cole, and Chunaram Choudhary. Time-Resolved Analysis Reveals Rapid Dynamics and Broad Scope of the CBP/p300 Acetylome. Cell, 174(1):231–244.e12, June 2018. ISSN 0092-8674. doi: 10.1016/j.cell.2018.04.033.

50. Esther Ortega, Srinivasan Rengachari, Ziad Ibrahim, Naghmeh Hoghoughi, Jonathan Gaucher, Alex S. Holehouse, Saadi Khochbin, and Daniel Panne. Transcription factor dimerization activates the p300 acetyl-transferase. Nature, 562(7728):538–544, October 2018. ISSN 1476-4687. doi: 10.1038/s41586-018-0621-1.

51. S Sampurno, A Bijenhof, D Cheasley, H Xu, S Robine, D Hilton, W S Alexander, L Pereira, T Mantamadiotis, J Malaterre, and R G Ramsay. The Myb-p300-CREB axis modulates intestine homeostasis, radiosensitivity and tumorigenesis. Cell Death Dis, 4(4):e605, April 2013. ISSN 2041-4889. doi: 10.1038/cddis.2013.119.

52. Menno P. Creyghton, Albert W. Cheng, G. Grant Welstead, Tristan Kooistra, Bryce W. Carey, Eveline J. Steine, Jacob Hanna, Michael A. Lodato, Garrett M. Frampton, Phillip A. Sharp, Laurie A. Boyer, Richard A. Young, and Rudolf Jaenisch. Histone H3K27ac separates active from poised enhancers and predicts developmental state. Proc. Natl. Acad. Sci. U.S.A., 107(50):21931–21936, December 2010. ISSN 1091-6490. doi: 10.1073/pnas.1016071107.

53. Xiaomin Zhong and Ying Jin. Critical Roles of Coactivator p300 in Mouse Embryonic Stem Cell Differentiation and Nanog Expression. J Biol Chem, 284(14):9168–9175, April 2009. ISSN 0021-9258. doi: 10.1074/jbc.M805562200.

54. Pier Lorenzo Puri, Maria Laura Avantaggiati, Clara Balsano, Nianli Sang, Adolf Graessmann, Antonio Giordano, and Massimo Levrero. P300 is required for MyoD-dependent cell cycle arrest and muscle-specific gene transcription. The EMBO Journal, 16(2):369–383, January 1997. ISSN 0261-4189. doi: 10.1093/emboj/16.2.369.

55. Maria Namwanje, Longhua Liu, Michelle Chan, Nikki Aaron, Michael J Kraakman, and Li Qiang. The depot-specific and essential roles of CBP/p300 in regulating adipose plasticity. J Endocrinol, 240(2):257–269, February 2019. ISSN 0022-0795. doi: 10.1530/JOE-18-0361.

56. Ayyub Ebrahimi, Kenan Sevinç, Gülben Gürhan Sevinç, Adam P. Cribbs, Martin Philpott, Fırat Uyulur, Tunçc Morova, James E. Dunford, Sencer Göklemez, SÇule Arı, Udo Oppermann, and Tamer T. Önder. Bromodomain inhibition of the coactivators CBP/EP300 facilitate cellular reprogramming. Nat. Chem. Biol., 15(5):519–528, May 2019. ISSN 1552-4469. doi: 10.1038/s41589-019-0264-z.

57. François Gerbe, Emmanuelle Sidot, Danielle J. Smyth, Makoto Ohmoto, Ichiro Matsumoto, Valérie Dardalhon, Pierre Cesses, Laure Garnier, Marie Pouzolles, Bénédicte Brulin, Marco Bruschi, Yvonne Harcus, Valérie S. Zimmermann, Naomi Taylor, Rick M. Maizels, and Philippe Jay. Intestinal epithelial tuft cells initiate type 2 mucosal immunity to helminth parasites. Nature, 529(7585):226–230, January 2016. ISSN 1476-4687. doi: 10.1038/nature16527.

58. Jakob von Moltke, Ming Ji, Hong-Erh Liang, and Richard M. Locksley. Tuft-cell-derived IL-25 regulates an intestinal ILC2-epithelial response circuit. Nature, 529(7585):221–225, January 2016. ISSN 1476-4687. doi: 10.1038/nature16161.

59. Akifumi Nakagawa, Curtis E. Adams, Yinshi Huang, Sulaiman R. Hamarneh, Wei Liu, Kate N. Von Alt, Mari Mino-Kenudson, Richard A. Hodin, Keith D. Lillemoe, Carlos Fernández-del Castillo, Andrew L. Warshaw, and Andrew S. Liss. Selective and reversible suppression of intestinal stem cell differentiation by pharmacological inhibition of BET bromodomains. Scientific Reports, 6(1):20390, February 2016. ISSN 2045-2322. doi: 10.1038/srep20390.

60. Lukas Huijbregts, Maja Borup Kjær Petersen, Claire Berthault, Mattias Hansson, Virginie Aiello, Latif Rachdi, Anne Grapin-Botton, Christian Honore, and Raphael Scharfmann. Bromodomain and Extra Terminal Protein Inhibitors Promote Pancreatic Endocrine Cell Fate. Diabetes, 68(4):761–773, April 2019. ISSN 0012-1797, 1939-327X. doi: 10.2337/db18-0224.

61. Hiroki Matsuda and Yun-Bo Shi. An essential and evolutionarily conserved role of protein arginine methyl-transferase 1 for adult intestinal stem cells during postembryonic development. Stem Cells, 28(11):2073–2083, November 2010. ISSN 1066-5099. doi: 10.1002/stem.529.

62. Atsuko Ishizuya-Oka and Yun-Bo Shi. Evolutionary insights into postembryonic development of adult intestinal stem cells. Cell Biosci, 1:37, November 2011. ISSN 2045-3701. doi: 10.1186/2045-3701-1-37.

63. Xiaomin Bao, Zurab Siprashvili, Brian J. Zarnegar, Rajani M. Shenoy, Eon J. Rios, Natalie Nady, Kun Qu, Angela Mah, Daniel E. Webster, Adam J. Rubin, Glenn G. Wozniak, Shiying Tao, Joanna Wysocka, and Paul A. Khavari. CSNK1a1 Regulates PRMT1 to Maintain the Progenitor State in Self-Renewing Somatic Tissue. Dev. Cell, 43(2):227–239.e5, October 2017. ISSN 1878-1551. doi: 10.1016/j.devcel.2017.08.021.

64. Suming Huang, Michael Litt, and Gary Felsenfeld. Methylation of histone H4 by arginine methyltransferase PRMT1 is essential in vivo for many subsequent histone modifications. Genes Dev., 19(16):1885–1893, August 2005. ISSN 0890-9369. doi: 10.1101/gad.1333905.

65. Stephanie M. Lehman, Hongshan Chen, Emmanuel S. Burgos, Maxim Maron, Sitaram Gayatri, Edward Nieves, Dina L. Bai, Simone Sidoli, Varun Gupta, Matthew R. Marunde, James R. Bone, Zu-Wen Sun, Mark T. Bedford, Jeffrey Shabanowitz, Donald F. Hunt, and David Shechter. Transcriptomic and proteomic regulation through abundant, dynamic, and independent arginine methylation by Type I and Type II PRMTs. bioRxiv, page 2020.06.23.167601, June 2020. doi: 10.1101/2020.06.23.167601.

66. Bolag Altan, Takehiko Yokobori, Munenori Ide, Erito Mochiki, Yoshitaka Toyomasu, Norimichi Kogure, Akiharu Kimura, Keigo Hara, Tuya Bai, Pinjie Bao, Masaki Suzuki, Kyoichi Ogata, Takayuki Asao, Masahiko Nishiyama, Tetsunari Oyama, and Hiroyuki Kuwano. Nuclear PRMT1 expression is associated with poor prognosis and chemosensitivity in gastric cancer patients. Gastric Cancer, 19(3):789–797, July 2016. ISSN 1436-3305. doi: 10.1007/s10120-015-0551-7.

67. Daniele Musiani, Roberto Giambruno, Enrico Massignani, Marica Rosaria Ippolito, Marianna Maniaci, Sriganesh Jammula, Daria Manganaro, Alessandro Cuomo, Luciano Nicosia, Diego Pasini, and Tiziana Bonaldi. PRMT1 Is Recruited via DNA-PK to Chromatin Where It Sustains the Senescence-Associated Secretory Phenotype in Response to Cisplatin. Cell Reports, 30(4):1208–1222.e9, January 2020. ISSN 2211-1247. doi: 10.1016/j.celrep.2019.12.061.

68. RG Morgan, E Mortensson, and AC Williams. Targeting LGR5 in Colorectal Cancer: Therapeutic gold or too plastic? Br J Cancer, 118(11):1410–1418, May 2018. ISSN 0007-0920. doi: 10.1038/s41416-018-0118-6.

69. Seri Choi, Hyeon-Ju Jeong, Hyebeen Kim, Dahee Choi, Sung-Chun Cho, Je Kyung Seong, Seung-Hoi Koo, and Jong-Sun Kang. Skeletal muscle-specific Prmt1 deletion causes muscle atrophy via deregulation of the PRMT6-FOXO3 axis. Autophagy, 15(6):1069–1081, June 2019. ISSN 1554-8627. doi: 10.1080/15548627.2019.1569931.

70. Jesse R. Raab, Deepthi Y. Tulasi, Kortney E. Wager, Jeremy M. Morowitz, Scott T. Magness, and Adam D. Gracz. Quantitative classification of chromatin dynamics reveals regulators of intestinal stem cell differentiation. Development, 147(1), January 2020. ISSN 0950-1991, 1477–9129. doi: 10.1242/dev.181966.

71. Takayuki Furumatsu, Masanao Tsuda, Kenji Yoshida, Noboru Taniguchi, Tatsuo Ito, Megumi Hashimoto, Takashi Ito, and Hiroshi Asahara. Sox9 and p300 cooperatively regulate chromatin-mediated transcription. J. Biol. Chem., 280(42):35203–35208, October 2005. ISSN 0021-9258. doi: 10.1074/jbc.M502409200.

72. Pauline Bastide, Charbel Darido, Julie Pannequin, Ralf Kist, Sylvie Robine, Christiane Marty-Double, Frédéric Bibeau, Gerd Scherer, Dominique Joubert, Frédéric Hollande, Philippe Blache, and Philippe Jay. Sox9 regulates cell proliferation and is required for Paneth cell differentiation in the intestinal epithelium. J Cell Biol, 178(4):635–648, August 2007. ISSN 0021-9525. doi: 10.1083/jcb.200704152.

73. Bum-Kyu Lee, Yujin Jang, Mijeong Kim, Lucy LeBlanc, Catherine Rhee, Jiwoon Lee, Samuel Beck, Wenwen Shen, and Jonghwan Kim. Super-enhancer-guided mapping of regulatory networks controlling mouse trophoblast stem cells. Nature Communications, 10(1):4749, October 2019. ISSN 2041-1723. doi: 10.1038/s41467-019-12720-6.

74. Fatima el Marjou, Klaus-Peter Janssen, Benny Hung-Junn Chang, Mei Li, Valérie Hindie, Lawrence Chan, Daniel Louvard, Pierre Chambon, Daniel Metzger, and Sylvie Robine. Tissue-specific and inducible Cre-mediated recombination in the gut epithelium. Genesis, 39(3):186–193, July 2004. ISSN 1526-954X. doi: 10.1002/gene.20042.

75. Marc A Kerenyi, Zhen Shao, Yu-Jung Hsu, Guoji Guo, Sidinh Luc, Kassandra O’Brien, Yuko Fujiwara, Cong Peng, Minh Nguyen, and Stuart H Orkin. Histone demethylase Lsd1 represses hematopoietic stem and progenitor cell signatures during blood cell maturation. eLife, 2:e00633, June 2013. ISSN 2050-084X. doi: 10.7554/eLife.00633.

76. Chamutal Bornstein, Shir Nevo, Amir Giladi, Noam Kadouri, Marie Pouzolles, François Gerbe, Eyal David, Alice Machado, Anna Chuprin, Beáta Tîoth, Ori Goldberg, Shalev Itzkovitz, Naomi Taylor, Philippe Jay, Valérie S. Zimmermann, Jakub Abramson, and Ido Amit. Single-cell mapping of the thymic stroma identifies IL-25-producing tuft epithelial cells. Nature, 559(7715):622–626, July 2018. ISSN 1476-4687. doi: 10.1038/s41586-018-0346-1.

77. Yung Hae Kim, Hjalte List Larsen, Pau Rué, Laurence A. Lemaire, Jorge Ferrer, and Anne Grapin-Botton. Cell cycle-dependent differentiation dynamics balances growth and endocrine differentiation in the pancreas. PLoS Biol., 13(3):e1002111, March 2015. ISSN 1545-7885. doi: 10.1371/journal.pbio.1002111.

78. Toshiro Sato and Hans Clevers. Primary Mouse Small Intestinal Epithelial Cell Cultures. In Scott H. Randell and M. Leslie Fulcher, editors, Epithelial Cell Culture Protocols: Second Edition, Methods in Molecular Biology, pages 319–328. 2013.

79. Kunihiro Kishida, Sarah C. Pearce, Shiyan Yu, Nan Gao, and Ronaldo P. Ferraris. Nutrient sensing by absorptive and secretory progenies of small intestinal stem cells. American Journal of Physiology-Gastrointestinal and Liver Physiology, 312(6):G592–G605, March 2017. ISSN 0193-1857. doi: 10.1152/ajpgi.00416.2016.

80. Lisa Luu, Zoe J. Matthews, Stuart D. Armstrong, Penelope P. Powell, Tom Wileman, Jonathan M. Wastling, and Janine L. Coombes. Proteomic Profiling of Enteroid Cultures Skewed toward Development of Specific Epithelial Lineages. PROTEOMICS, 18(16):1800132, 2018. ISSN 1615-9861. doi: 10.1002/pmic.201800132.

81. Johannes Schindelin, Ignacio Arganda-Carreras, Erwin Frise, Verena Kaynig, Mark Longair, Tobias Pietzsch, Stephan Preibisch, Curtis Rueden, Stephan Saalfeld, Benjamin Schmid, Jean-Yves Tinevez, Daniel James White, Volker Hartenstein, Kevin Eliceiri, Pavel Tomancak, and Albert Cardona. Fiji: An open-source platform for biological-image analysis. Nature Methods, 9(7):676–682, July 2012. ISSN 1548-7105. doi: 10.1038/nmeth.2019.

82. Johannes Schindelin, Curtis T. Rueden, Mark C. Hiner, and Kevin W. Eliceiri. The ImageJ ecosystem: An open platform for biomedical image analysis. Molecular Reproduction and Development, 82(7-8):518–529, 2015. ISSN 1098-2795. doi: 10.1002/mrd.22489.

83. Stuart Berg, Dominik Kutra, Thorben Kroeger, Christoph N. Straehle, Bernhard X. Kausler, Carsten Haubold, Martin Schiegg, Janez Ales, Thorsten Beier, Markus Rudy, Kemal Eren, Jaime I. Cervantes, Buote Xu, Fynn Beuttenmueller, Adrian Wolny, Chong Zhang, Ullrich Koethe, Fred A. Hamprecht, and Anna Kreshuk. Ilastik: Interactive machine learning for (bio)image analysis. Nature Methods, 16(12): 1226–1232, December 2019. ISSN 1548-7105. doi: 10.1038/s41592-019-0582-9.

84. Phu Van, Wenxin Jiang, Raphael Gottardo, and Greg Finak. ggCyto: Next generation open-source visualization software for cytometry. Bioinformatics, 34(22):3951–3953, November 2018. ISSN 1367-4803. doi: 10.1093/bioinformatics/bty441.

85. Guangchuang Yu, Tommy Tsan-Yuk Lam, Huachen Zhu, and Yi Guan. Two Methods for Mapping and Visualizing Associated Data on Phylogeny Using Ggtree. Mol Biol Evol, 35(12):3041–3043, December 2018. ISSN 0737-4038. doi: 10.1093/molbev/msy194.

86. Alexander Dobin, Carrie A. Davis, Felix Schlesinger, Jorg Drenkow, Chris Zaleski, Sonali Jha, Philippe Batut, Mark Chaisson, and Thomas R. Gingeras. STAR: Ultrafast universal RNA-seq aligner. Bioinformatics, 29(1):15–21, January 2013. ISSN 1367-4803. doi: 10.1093/bioinformatics/bts635.

87. Philip Ewels, Måns Magnusson, Sverker Lundin, and Max Käller. MultiQC: Summarize analysis results for multiple tools and samples in a single report. Bioinformatics, 32(19):3047–3048, October 2016. ISSN 1367-4803. doi: 10.1093/bioinformatics/btw354.

88. Richard Mark Leggett, Ricardo Humberto Ramirez-Gonzalez, Bernardo Clavijo, Darren Waite, and Robert Paul Davey. Sequencing quality assessment tools to enable data-driven informatics for high throughput genomics. Front. Genet., 4, 2013. ISSN 1664-8021. doi: 10.3389/fgene.2013.00288.

89. Adam Frankish, Mark Diekhans, Anne-Maud Ferreira, Rory Johnson, Irwin Jungreis, Jane Loveland, Jonathan M. Mudge, Cristina Sisu, James Wright, Joel Armstrong, If Barnes, Andrew Berry, Alexandra Bignell, Silvia Carbonell Sala, Jacqueline Chrast, Fiona Cunningham, Tomás Di Domenico, Sarah Donaldson, Ian T. Fiddes, Carlos Garcîıa Girón, Jose Manuel Gonzalez, Tiago Grego, Matthew Hardy, Thibaut Hourlier, Toby Hunt, Osagie G. Izuogu, Julien Lagarde, Fergal J. Martin, Laura Martîınez, Shamika Mo-hanan, Paul Muir, Fabio C. P. Navarro, Anne Parker, Baikang Pei, Fernando Pozo, Magali Ruffier, Bianca M. Schmitt, Eloise Stapleton, Marie-Marthe Suner, Irina Sycheva, Barbara Uszczynska-Ratajczak, Jinuri Xu, Andrew Yates, Daniel Zerbino, Yan Zhang, Bronwen Aken, Jyoti S. Choudhary, Mark Gerstein, Roderic Guigó, Tim J. P. Hubbard, Manolis Kellis, Benedict Paten, Alexandre Reymond, Michael L. Tress, and Paul Flicek. GENCODE reference annotation for the human and mouse genomes. Nucleic Acids Res, 47(D1): D766–D773, January 2019. ISSN 0305-1048. doi: 10.1093/nar/gky955.

90. Yang Liao, Gordon K. Smyth, and Wei Shi. featureCounts: An efficient general purpose program for assigning sequence reads to genomic features. Bioinformatics, 30(7):923–930, April 2014. ISSN 1367-4811. doi: 10.1093/bioinformatics/btt656.

91. Michael I. Love, Wolfgang Huber, and Simon Anders. Moderated estimation of fold change and dispersion for RNA-seq data with DESeq2. Genome Biology, 15(12):550, December 2014. ISSN 1474-760X. doi: 10.1186/s13059-014-0550-8.

92. Anqi Zhu, Joseph G. Ibrahim, and Michael I. Love. Heavy-tailed prior distributions for sequence count data: Removing the noise and preserving large differences. Bioinformatics, 35(12):2084–2092, June 2019. ISSN 1367-4803. doi: 10.1093/bioinformatics/bty895.

93. Guangchuang Yu, Li-Gen Wang, Yanyan Han, and Qing-Yu He. clusterProfiler: An R Package for Comparing Biological Themes Among Gene Clusters. OMICS: A Journal of Integrative Biology, 16(5):284–287, March 2012. doi: 10.1089/omi.2011.0118.

94. Damian Szklarczyk, Annika L. Gable, David Lyon, Alexander Junge, Stefan Wyder, Jaime Huerta-Cepas, Milan Simonovic, Nadezhda T. Doncheva, John H. Morris, Peer Bork, Lars J. Jensen, and Christian von Mering. STRING v11: Protein–protein association networks with increased coverage, supporting functional discovery in genome-wide experimental datasets. Nucleic Acids Res, 47(D1):D607–D613, January 2019. ISSN 0305-1048. doi: 10.1093/nar/gky1131.

95. Heonjong Han, Jae-Won Cho, Sangyoung Lee, Ayoung Yun, Hyojin Kim, Dasom Bae, Sunmo Yang, Chan Yeong Kim, Muyoung Lee, Eunbeen Kim, Sungho Lee, Byunghee Kang, Dabin Jeong, Yaeji Kim, Hyeon-Nae Jeon, Haein Jung, Sunhwee Nam, Michael Chung, Jong-Hoon Kim, and Insuk Lee. TRRUST v2: An expanded reference database of human and mouse transcriptional regulatory interactions. Nucleic Acids Res, 46(D1):D380–D386, January 2018. ISSN 0305-1048. doi: 10.1093/nar/gkx1013.

96. Steffen Durinck, Yves Moreau, Arek Kasprzyk, Sean Davis, Bart De Moor, Alvis Brazma, and Wolfgang Huber. BioMart and Bioconductor: A powerful link between biological databases and microarray data analysis. Bioinformatics, 21(16):3439–3440, August 2005. ISSN 1367-4803. doi: 10.1093/bioinformatics/bti525.

97. The Gene Ontology Consortium. The Gene Ontology Resource: 20 years and still GOing strong. Nucleic Acids Res., 47(D1):D330–D338, January 2019. ISSN 1362-4962. doi: 10.1093/nar/gky1055.

98. Hadley Wickham, Mara Averick, Jennifer Bryan, Winston Chang, Lucy D’Agostino McGowan, Romain François, Garrett Grolemund, Alex Hayes, Lionel Henry, Jim Hester, Max Kuhn, Thomas Lin Pedersen, Evan Miller, Stephan Milton Bache, Kirill Möller, Jeroen Ooms, David Robinson, Dana Paige Seidel, Vitalie Spinu, Kohske Takahashi, Davis Vaughan, Claus Wilke, Kara Woo, and Hiroaki Yutani. Welcome to the Tidyverse. Journal of Open Source Software, 4(43):1686, November 2019. ISSN 2475-9066. doi: 10.21105/joss.01686.

99. Eleanor Williams, Josh Moore, Simon W. Li, Gabriella Rustici, Aleksandra Tarkowska, Anatole Chessel, Simone Leo, Bálint Antal, Richard K. Ferguson, Ugis Sarkans, Alvis Brazma, Rafael E. Carazo Salas, and Jason R. Swedlow. Image Data Resource: A bioimage data integration and publication platform. Nature Methods, 14(8):775–781, August 2017. ISSN 1548-7105. doi: 10.1038/nmeth.4326.

100. Ugis Sarkans, Mikhail Gostev, Awais Athar, Ehsan Behrangi, Olga Melnichuk, Ahmed Ali, Jasmine Minguet, Juan Camillo Rada, Catherine Snow, Andrew Tikhonov, Alvis Brazma, and Johanna McEntyre. The Bio-studies database—one stop shop for all data supporting a life sciences study. Nucleic Acids Res, 46(D1): D1266–D1270, January 2018. ISSN 460. doi: 10.1093/nar/gkx965.

101. Awais Athar, Anja Föllgrabe, Nancy George, Haider Iqbal, Laura Huerta, Ahmed Ali, Catherine Snow, Nuno A. Fonseca, Robert Petryszak, Irene Papatheodorou, Ugis Sarkans, and Alvis Brazma. ArrayExpress update – from bulk to single-cell expression data. Nucleic Acids Res, 47(D1):D711–D715, January 2019. ISSN 470. doi: 10.1093/nar/gky964.

