## Supplementary Figures for "A semi-automated organoid screening method demonstrates epigenetic control of intestinal epithelial differentiation"

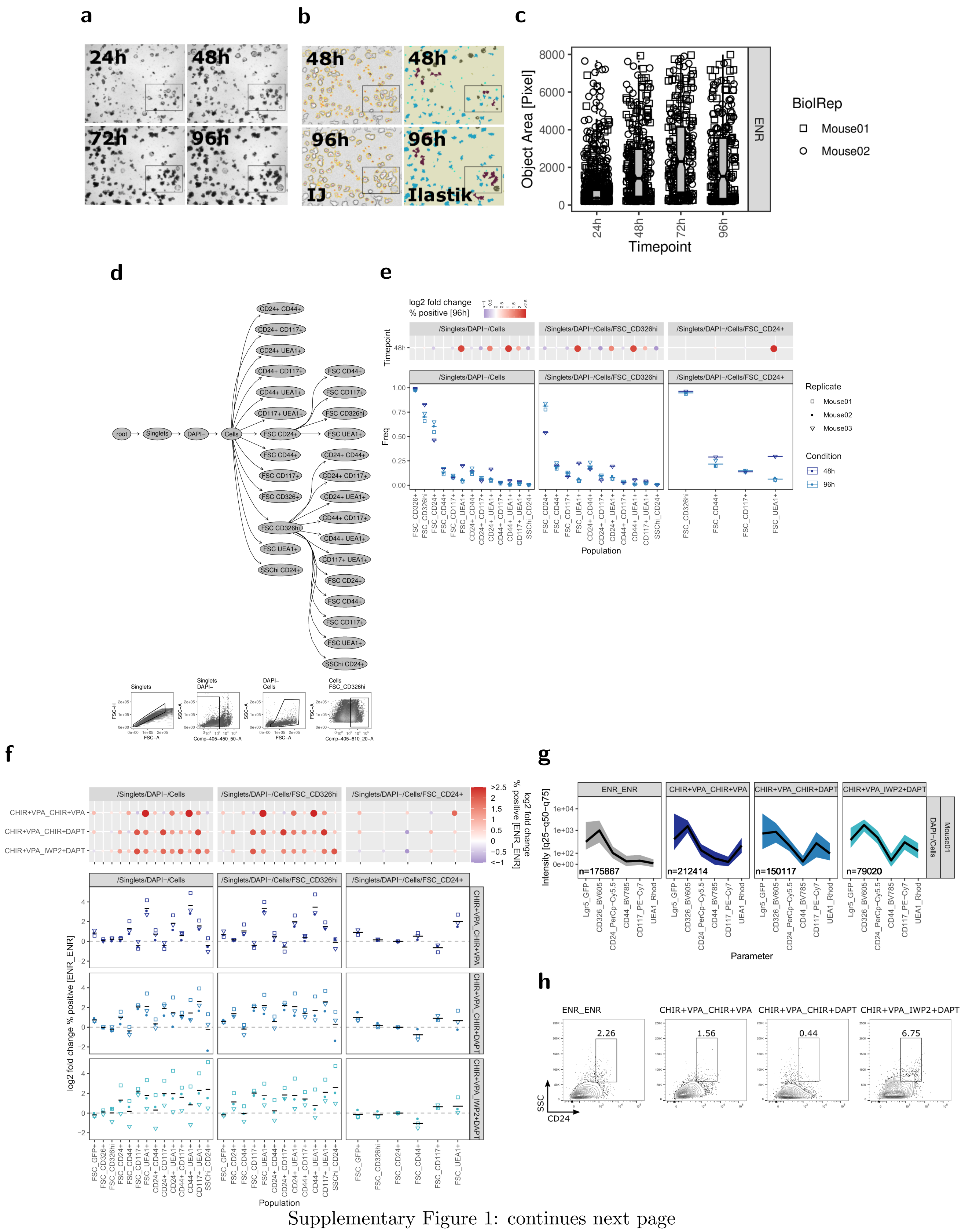

Supplementary Figure 1: continues next page

Supplementary Figure 1:
a) Organoid growth 24-96h, representative well from experiment quantified in **Fig. 1c**. The area shown in **Fig. 1a** is marked.
b) Organoid segmentation with ImageJ/Fiji (”IJ”, left) and combined ImageJ/Fiji and Ilastik (”Ilastik”, right) workflow, 48h and 96h timepoints. The ImageJ segmentation sufficiently identifies objects while the combined workflow is more accurate and can distinguish phenotypes such as organoid, sphere, cluster, or debris. c) Quantification result with ImageJ workflow. Pooled data from 2 biol. replicates.
d) Flow cytometry gating strategy.
e) Flow cytometry of organoids grown for 48h and 96h. Population frequencies of flow cytometry staining in Cells, Cells/CD326$^{hi}$ and Cells/CD24+ parent gate, 3 biol. replicates, indicated by shape, mean highlighted (bottom). Log2 fold change normalized to 96h timepoint, visualized as dot plot, dot size corresponds to absolute log2 fold change (top).
f) Flow cytometry of organoids cultured for 48h followed by 72h (48h_72h) with normal culture medium (ENR_ENR), or culture medium containing CHIR+VPA_CHIR+VPA, CHIR+VPA_CHIR+DAPT, or CHIR+VPA_IWP2+DAPT to modify IEC composition. Population frequencies in Cells, Cells/CD326$^{hi}$ and Cells/CD24+ parent gate. Log2 fold change normalized to ENR_ENR treatment. 3 biol. replicates, indicated by shape, mean highlighted (bottom) and visualized as dot plot, dot size corresponds to absolute log2 fold change (top).
g) Flow cytometry of organoids cultured for 48h followed by 72h (48h_72h) as indicated. Staining of representative replicate, fluorescence intensity median and q25 and q75 percentile. Cell count in Cells parent gate is indicated.
h) Flow cytometry of organoids cultured for 48h followed by 72h (48h_72h) as indicated. SSC vs. CD24_PerCp-Cy5.5 staining in Cells parent gate, frequency in SSC$^{hi}$_CD24+ gate is indicated.

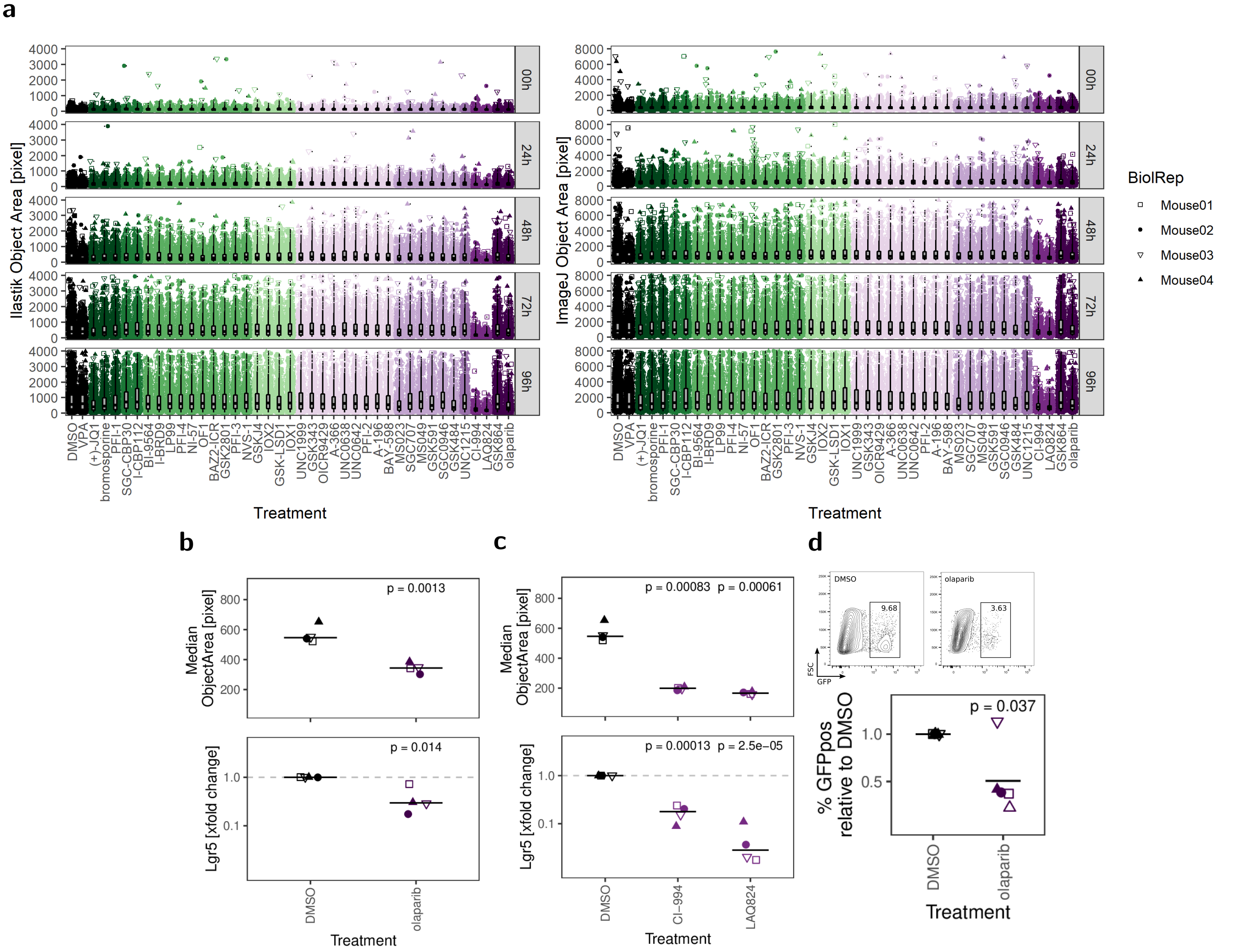


Supplementary Figure 2:
a) Segmentation results of combined ImageJ/Ilastik (left) and ImageJ (right) workflow of organoids treated with DMSO or inhibitors for 0-96h. 4 biol. replicates.
b) Median organoid area and relative *Lgr5* gene expression in DMSO or olaparib-treated organoids. 4 biol. replicates, indicated by shape. Median highlighted. Paired t-test.
c) Median organoid area and relative *Lgr5* gene expression in DMSO, CI-994, or LAQ824-treated organoids. 4 biol. replicates. Median highlighted. Paired t-test.
d) Frequency of *Lgr5*-EGFP stem cells in reporter organoids treated with DMSO or olaparib, measured by flow cytometry. Gating of representative replicate (top) and percentage of GFP+ cells normalized to DMSO condition of 5 biol. replicates, indicated by shape. Mean highlighted. Paired t-test (bottom). Minimum 5000 viable cells in parent gate.

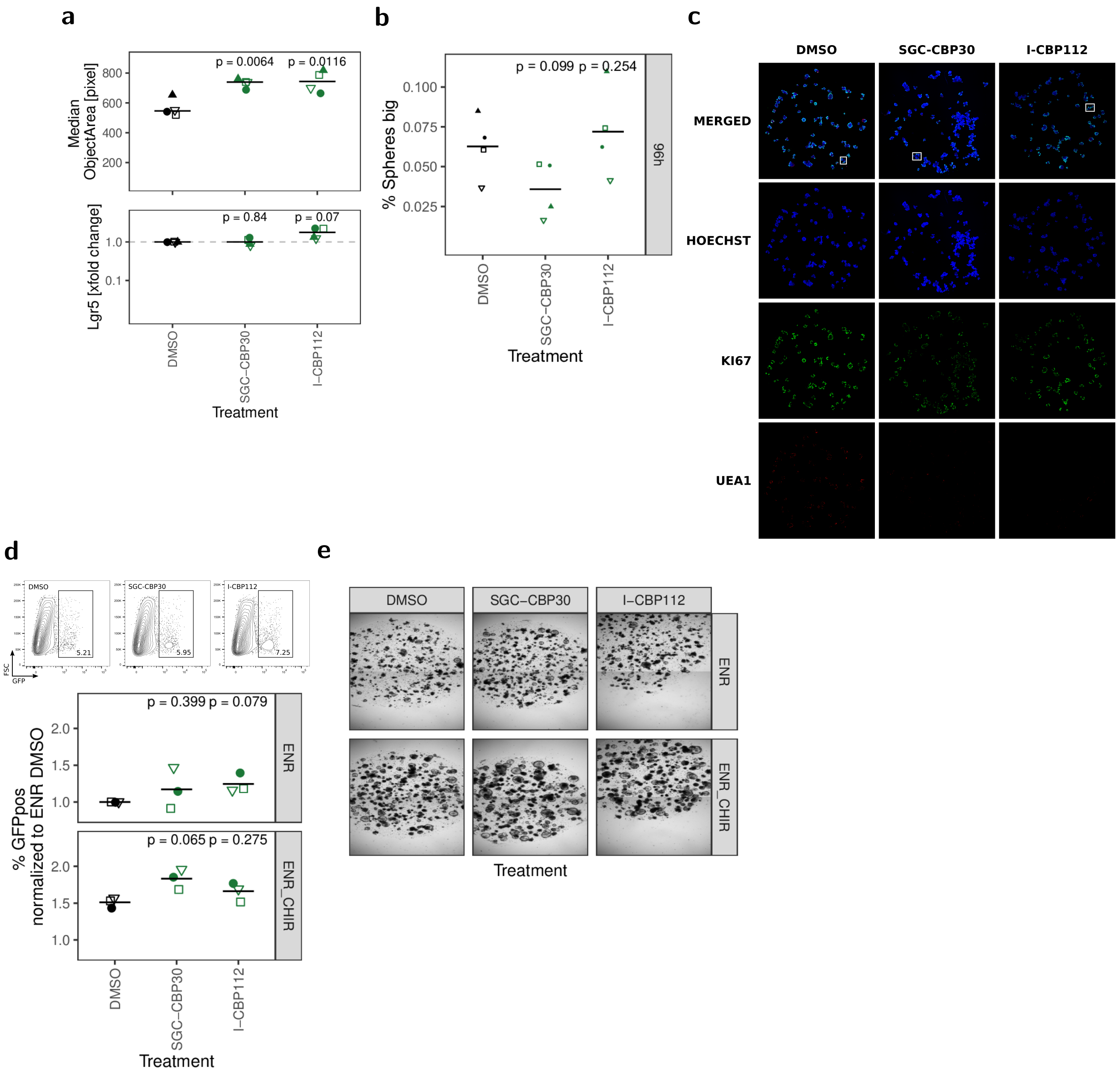


Supplementary Figure 3: continues next page

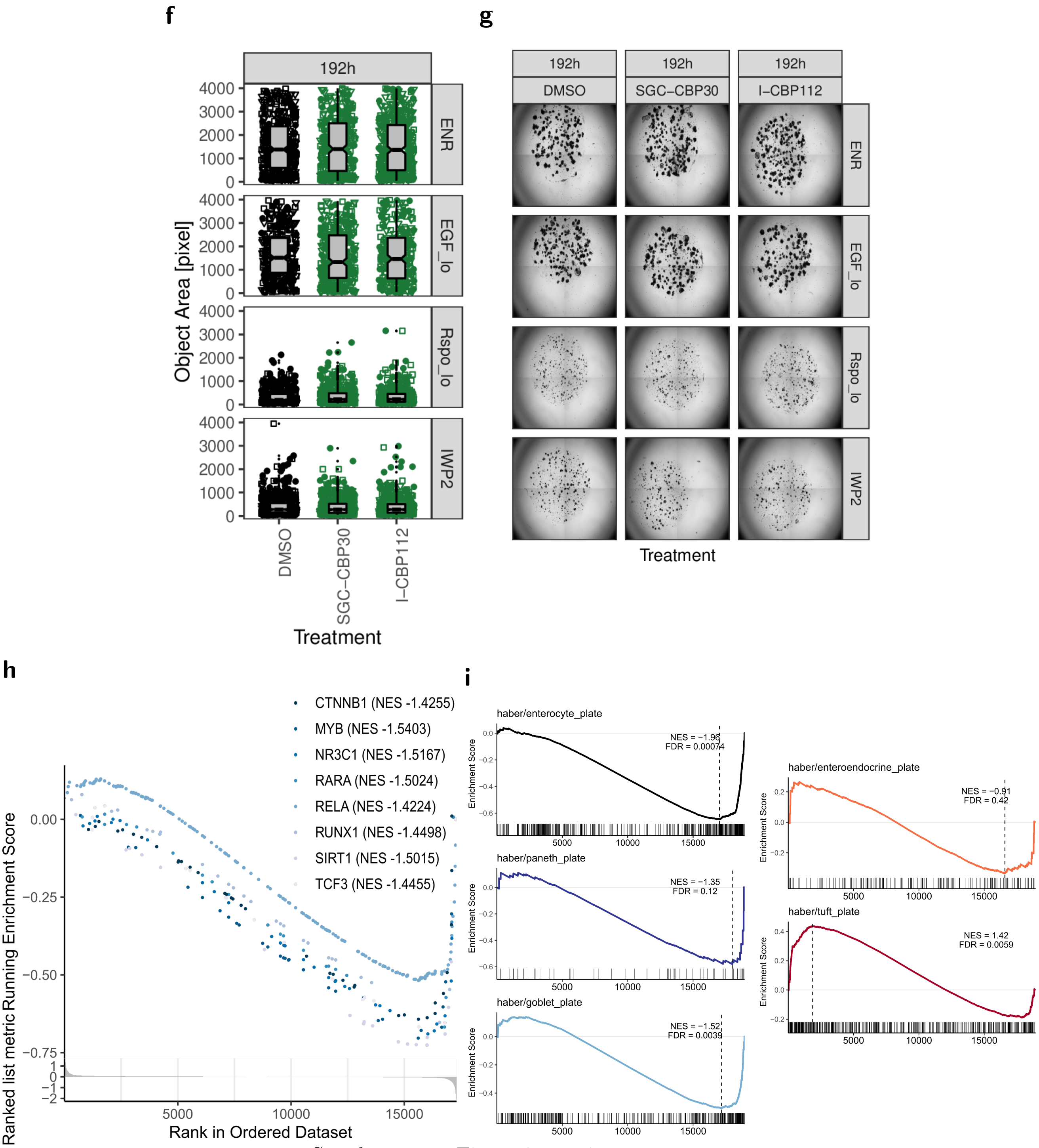


Supplementary Figure 3: continues next page

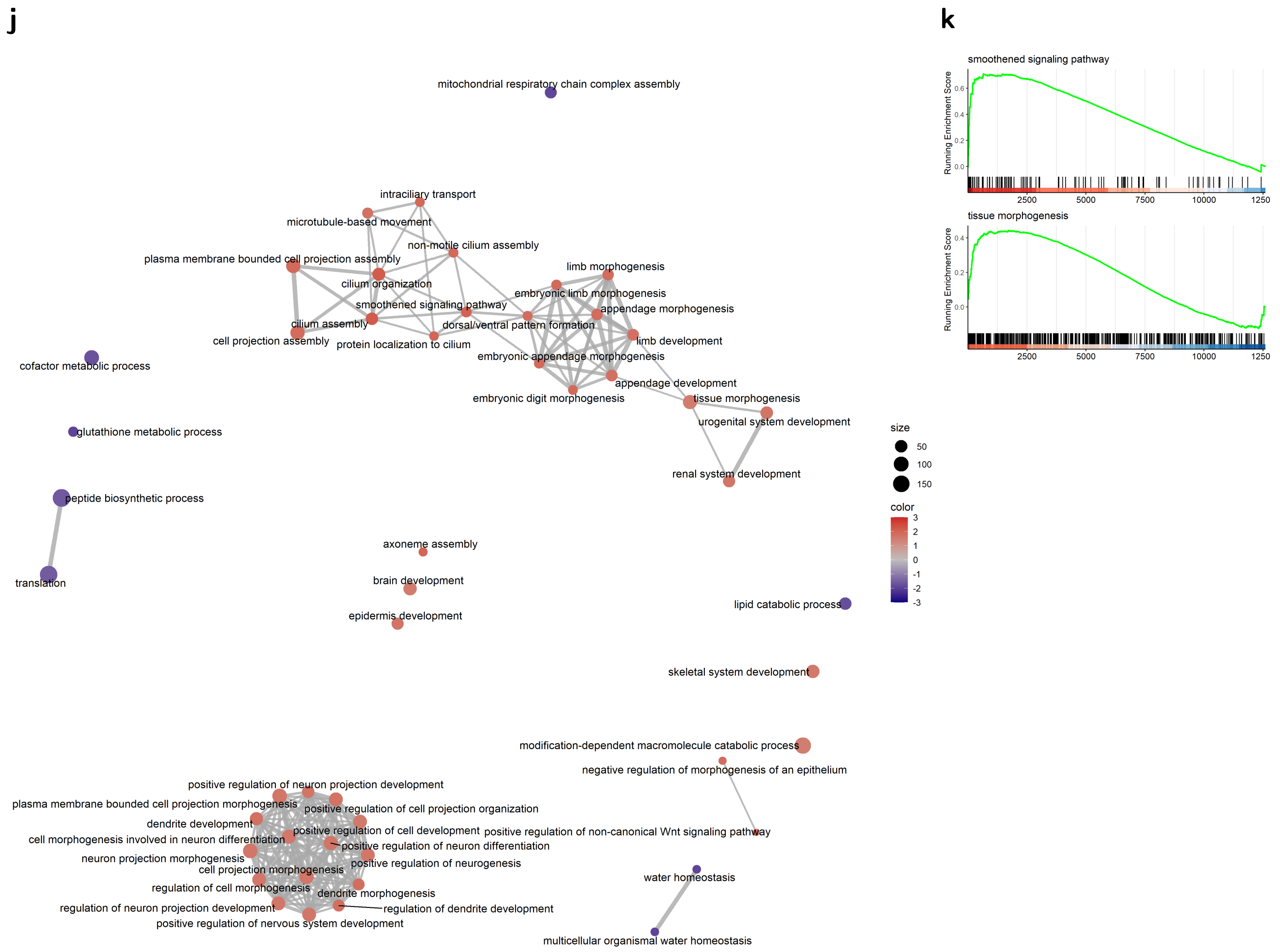


Supplementary Figure 3:

a) Median organoid area and relative *Lgr5* gene expression in DMSO, SCG-CBP30, or I-CBP112 treated organoids. 4 biol. replicates, indicated by shape. Median highlighted. Paired t-test.

b) Organoids treated with DMSO, SGC-CBP30, or I-CBP112 for 96h. Percentage of organoids classified as big spheres in combined ImageJ/Ilastik segmentation workflow. 4 biol. replicates, indicated by shape. Mean highlighted. Paired t-test.

c) Confocal microscopy of DMSO, SCG-CBP30, or I-CBP112 treated organoids. 10x magnification, max. intensity projection. Full well, the organoids shown in Fig. 2b are marked.

d) Frequency of *Lgr5*-EGFP stem cells in reporter organoids grown in ENR or ENR + CHIR and treated with DMSO, SCG-CBP30, or I-CBP112, measured by flow cytometry. Gating of representative replicate (top) and percentage of GFP+ cells normalized to ENR DMSO condition of 3 biol. replicates, indicated by shape. Mean highlighted. Paired t-test (bottom).

e) Representative replicate of *Lgr5*-EGFP reporter organoids grown in ENR or ENR + CHIR and treated with DMSO, SCG-CBP30, or I-CBP112 for 96h.

f) Object area of organoids grown in ENR, ENR + low R-Spondin (1%), ENR + low EGF (1ng/ml), or ENR + IWP2 (Wnt pathway inhibitor) for 192h, treated with DMSO, SGC-CBP30, or I-CBP112. 3 biol. replicates, indicated by shape.

g) Representative Replicate of organoids grown in ENR, or low growth factor conditions for 192h, treated with DMSO, SGC-CBP30 or I-CBP112.

h) mRNA sequencing of untreated vs. I-CBP112 treated organoids. GSEA of target genes of the convergence of murine and human transcription factors from the TRRUST transcriptional interactions database and proteins interacting with murine or human EP300 or CREBBP with a minimum medium experimental confidence level ($\geq 0.4$) from the STRING protein-protein interactions database. Enriched transcription factor target gene sets with a p-value $\geq 0.1$ are shown. All normalized enrichment score (NES) values are indicated.

i) mRNA sequencing of untreated vs. I-CBP112 treated organoids. GSEA for enterocyte, Paneth cell, goblet cell, enteroendocrine cell, tuft cell signatures from Haber et al. (GSE92332). j) GSEA for Gene Ontology biological process (GO:BP) terms. Normalized enrichment score (NES) is indicated by color. Top 50 categories are shown.

k) GSEA for Gene Ontology term "smoothened signaling pathway" (GO:0007224, NES=2.0406), and "tissue morphogenesis" (GO:0048729, NES=1.5148).

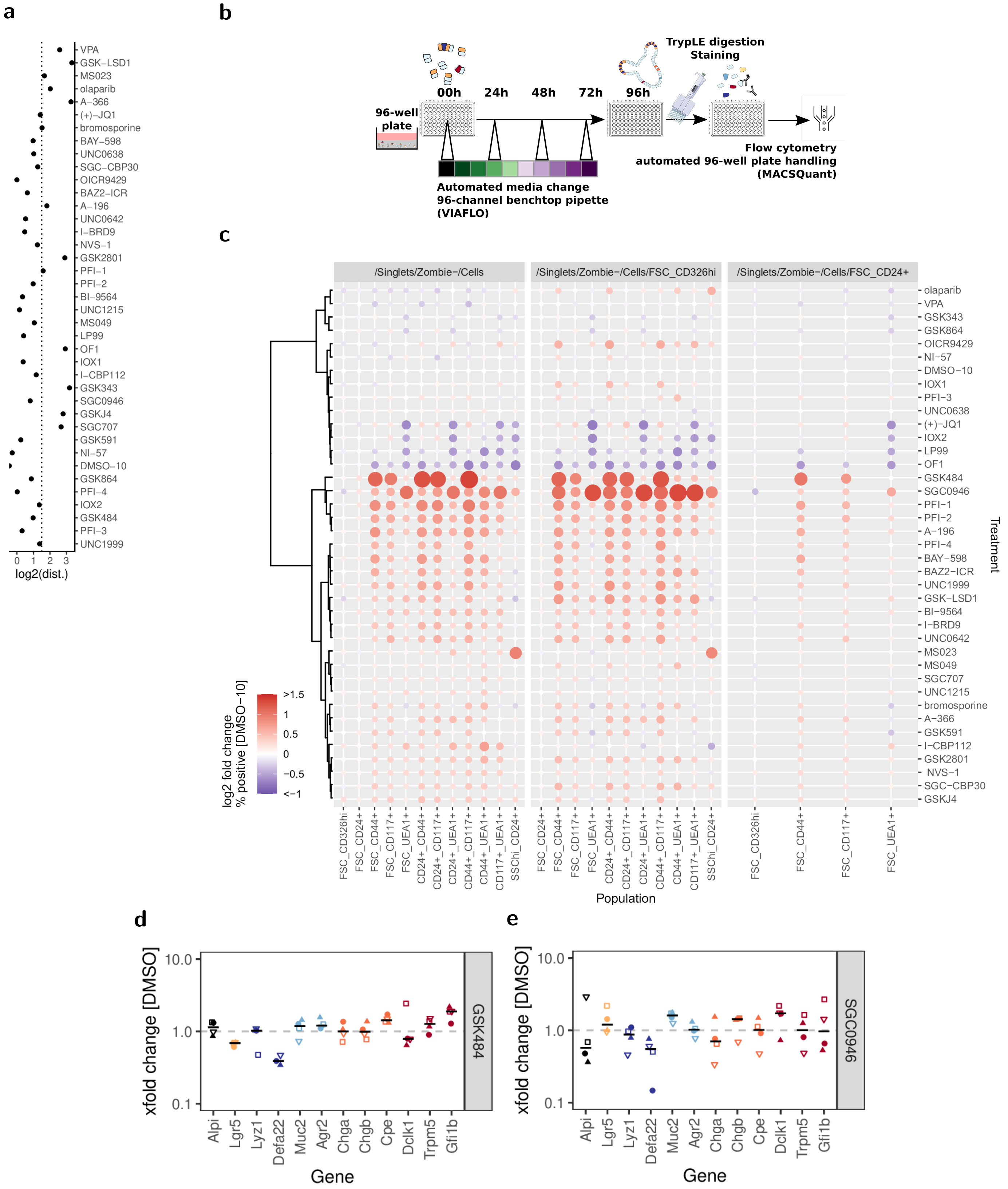


Supplementary Figure 4: continues next page

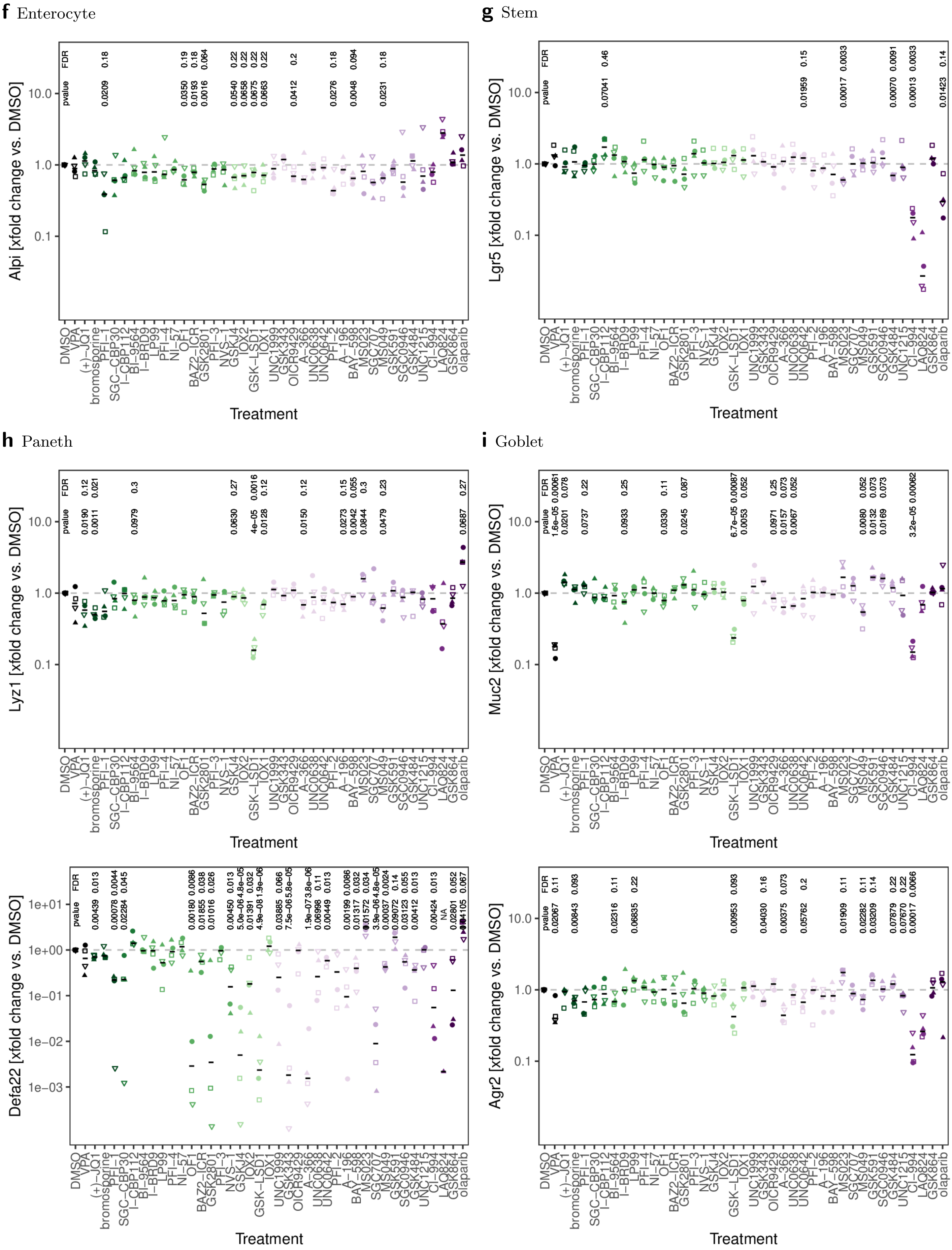


Supplementary Figure 4: continues next page

**j** Enteroendocrine

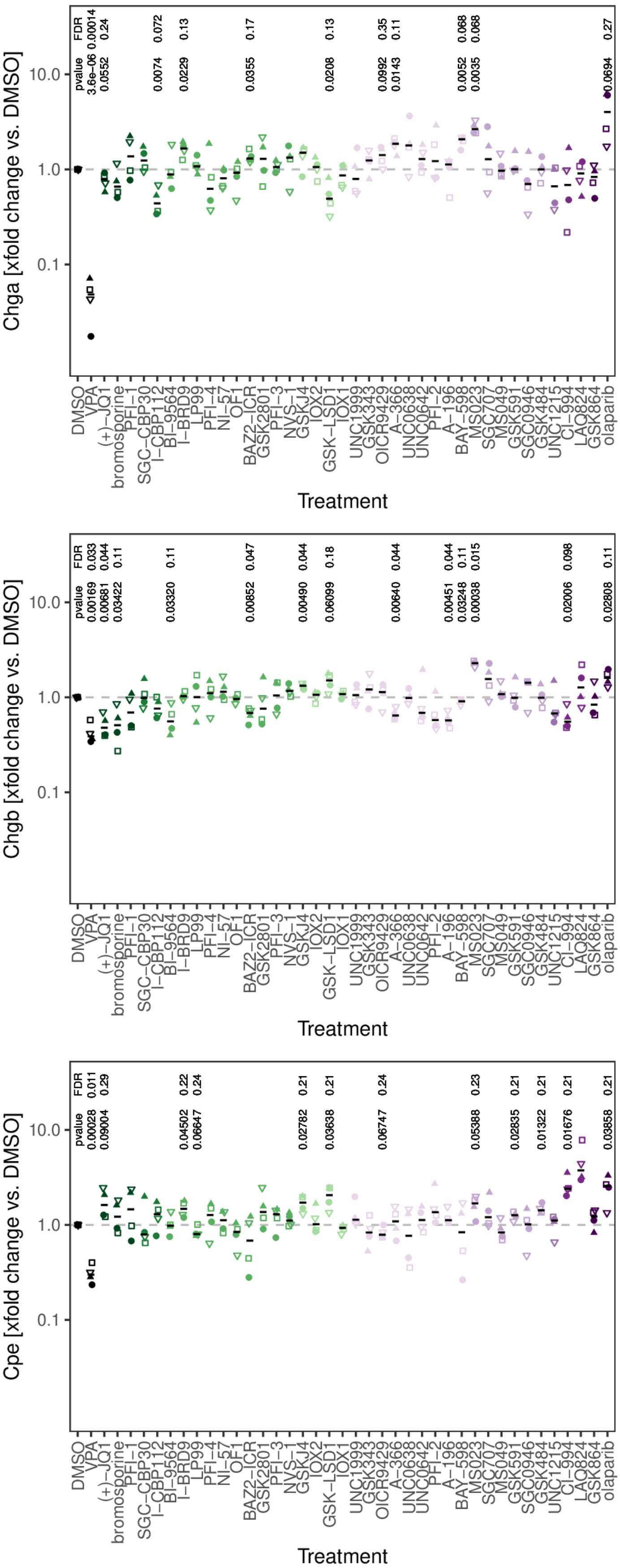


**k** Tuft

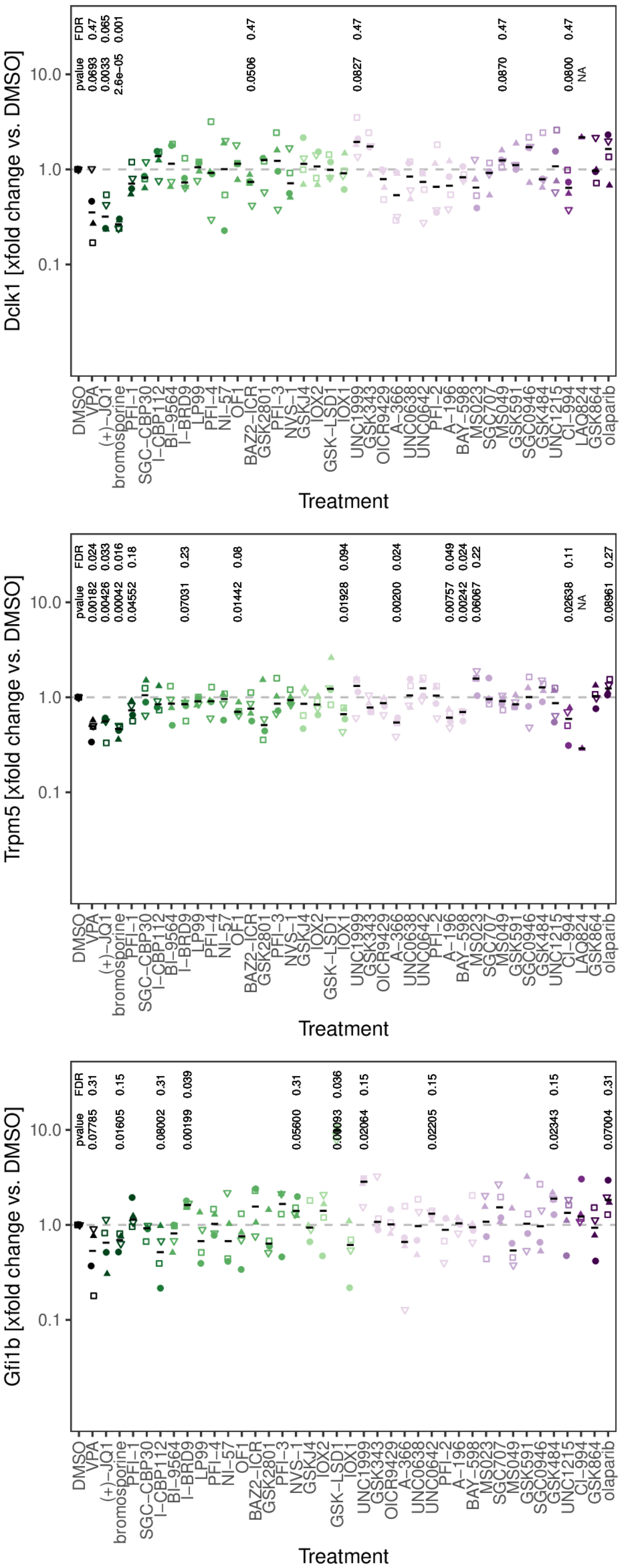


Supplementary Figure 4: continues next page

l

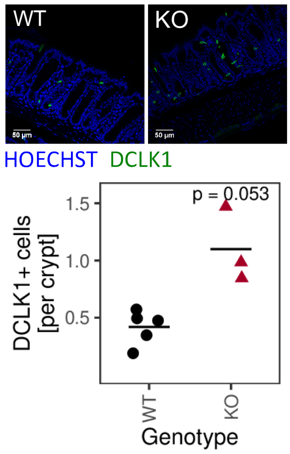


Supplementary Figure 4:
a) Log2 of the euclidean distance ("perturbation") of IEC lineage marker gene expression of organoids treated with DMSO or inhibitors for 96h, including gene *Defa22*. Samples treated with HDAC inhibitors were excluded from the analysis. Order is the same as in **Fig. 4a**.
b) Scheme of flow cytometry screen. Organoids were cultivated in 96-well plates, culture media was changed daily. All pipetting steps were carried out with automated pipettes or multichannel pipettes and samples were acquired on a flow cytometer with automated plate handling.
c) Flow cytometry screening of organoids, treated with DMSO or inhibitors for 96h. Population frequencies in Cells, FSC_CD326$^{hi}$ and FSC_CD24+ parent gate. Log2 fold change of gated populations normalized to DMSO treatment. Median of 4wells/3 biol. replicates, samples with $\leq$ 4000 cells and populations containing $\leq$40 cells were excluded. Dot size corresponds to absolute log2 fold change. Tree is based on euclidean distance clustering of populations in Cells parent gate.
d) Gene expression of organoids treated with DMSO or GSK484 for 96h, measured by qRT-PCR. GSK484 is an inhibitor of arginine deiminase (PADI4). 4 biol. replicates, indicated by shape. Median highlighted.
e) Gene expression of organoids treated with DMSO or SGC0946 for 96h, measured by qRT-PCR. SGC0946 is an inhibitor of H3 lysine-79 specific Histone-lysine N-methyltransferase (DOT1L). 4 biol. replicates, indicated by shape. Median highlighted.
f-k) Gene expression of *Alpi*, *Lgr5*, *Lyz1*, *Defa22*, *Muc2*, *Agr2*, *Chga*, *Chgb*, *Cpe*, *Dclk1*, *Trpm5* *Gfi1b* in organoids treated with DMSO or inhibitors for 96h, measured by qRT-PCR. xfold change relative to DMSO-treated organoids. 4 biol. replicates, indicated by shape. Median highlighted. Paired t-test and FDR shown for p-values $\leq$0.1.
l) DCLK1+ cells per crypt in colon of *Villin*-Cre+ *Lsd1*$^{f/f}$ (KO) mice with intestine-specific deletion of *Lsd1* and wild type (WT) littermates. Immunofluorescence staining of tissue sections. Representative staining (left) and quantification in 5/3 mice, mean highlighted. Unpaired t-test (right).

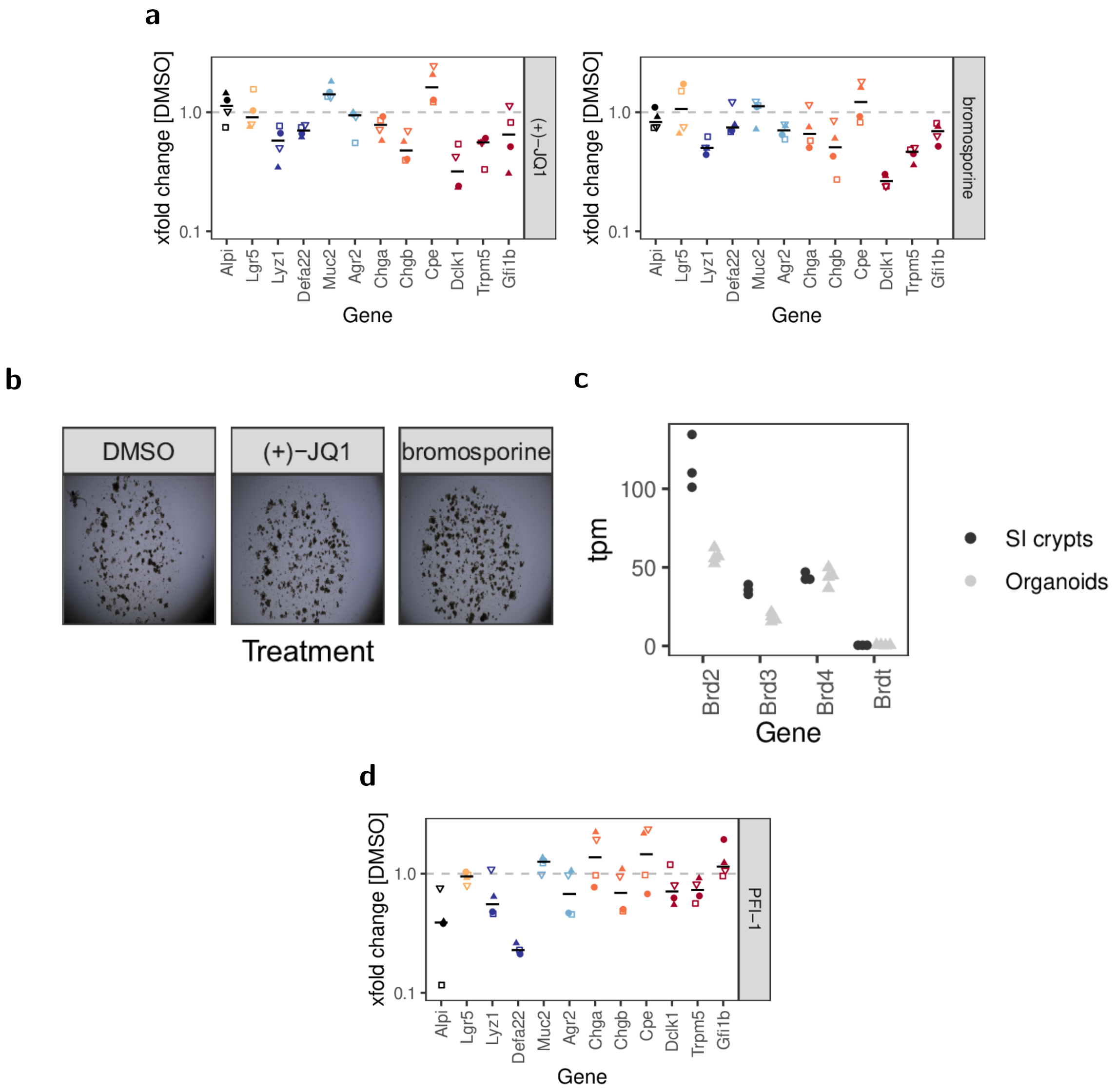


Supplementary Figure 5:

a) Gene expression of organoids treated with (+)-JQ1 (left) or bromosporine (right) for 96h, measured by qRT-PCR. xfold change relative to DMSO-treated organoids. 4 biol. replicates, indicated by shape. Median highlighted.

b) Organoids treated with (+)-JQ1 or bromosporine for 96h. Representative replicate from screen experiment.

c) BRD tpm values in mRNA sequencing datasets of small intestinal crypts (3 mice) and organoids (4 biol. replicates) from Zwiggelaar et al. (control groups from E-MTAB-9077, E-MTAB-7862).

d) Gene expression of organoids treated with PFI-1 for 96h, measured by qRT-PCR. PFI-1 is an inhibitor of BRD2/BRD4. xfold change relative to DMSO-treated organoids. 4 biol. replicates, indicated by shape. Median highlighted.

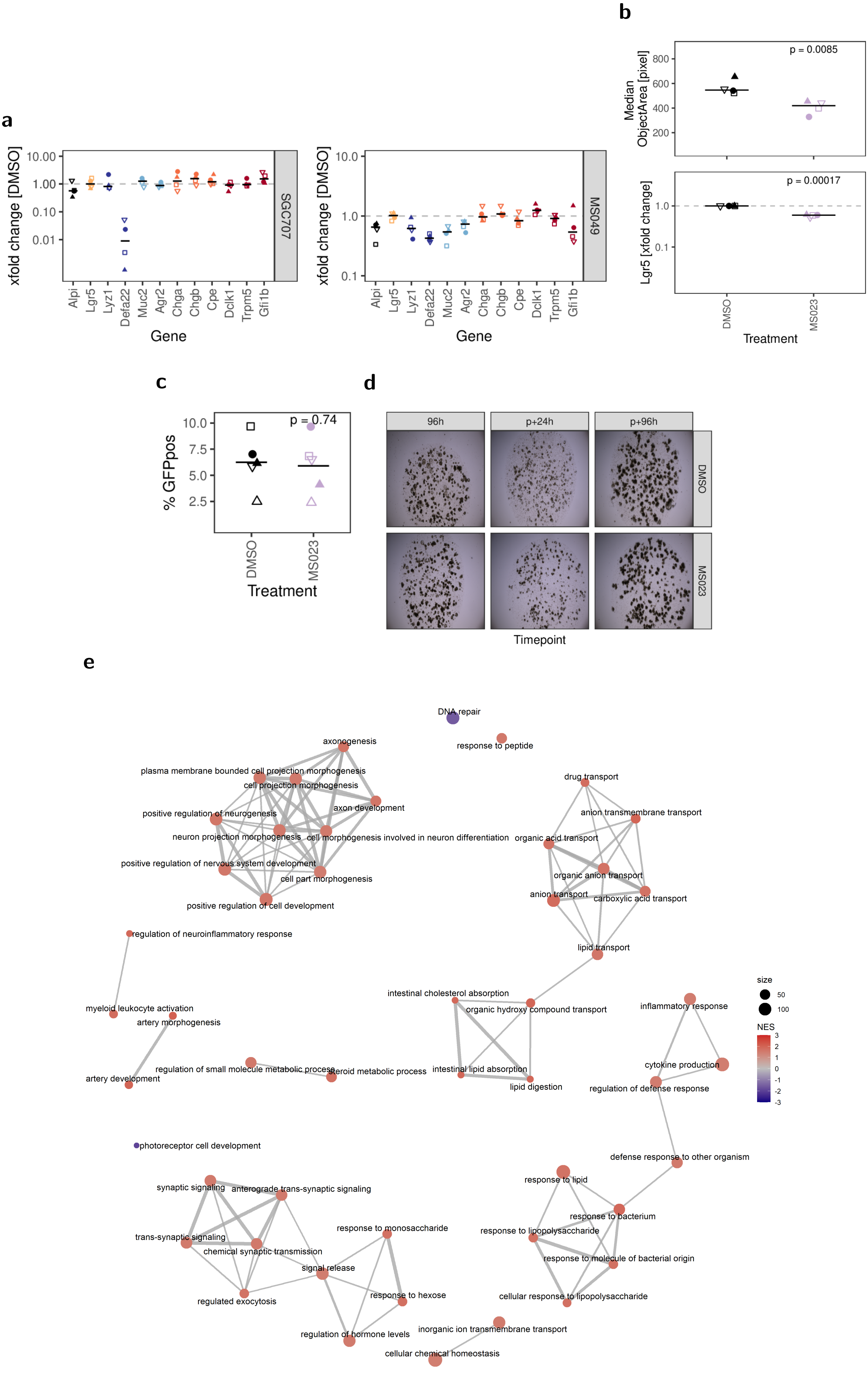


Supplementary Figure 6: continues next page

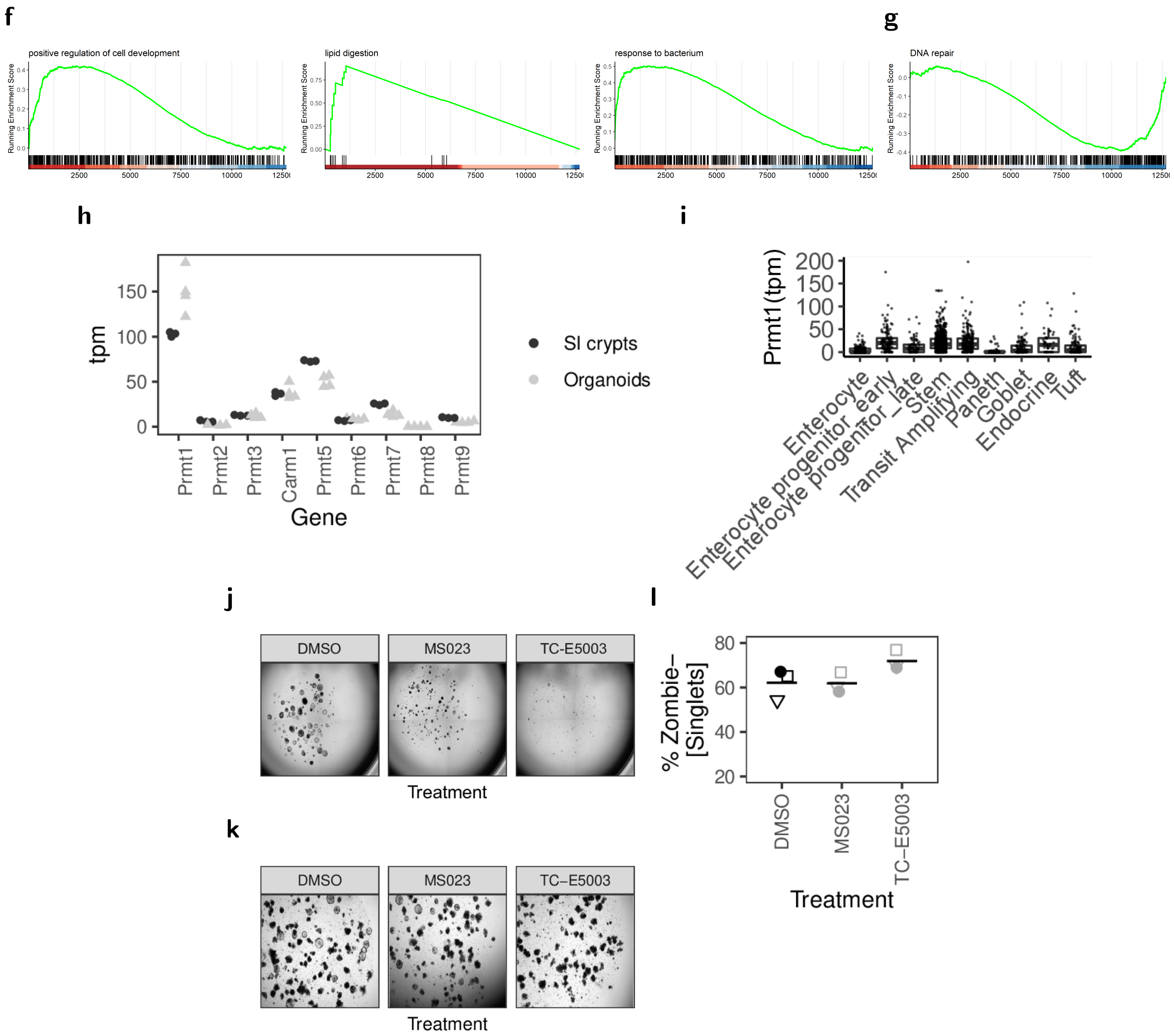


Supplementary Figure 6:

a) Gene expression of organoids treated with SGC707 (top) or MS049 (bottom) for 96h, measured by qRT-PCR. xfold change relative to DMSO-treated organoids. 4 biol. replicates, indicated by shape. Median highlighted.

b) Median organoid area and relative *Lgr5* gene expression in DMSO or MS023-treated organoids. 4 biol. replicates, indicated by shape. Median highlighted. Paired t-test.

c) Frequency of *Lgr5*-EGFP stem cells in reporter organoids treated with DMSO or MS023 for 96h, measured by flow cytometry. 5 biol. replicates, indicated by shape. Mean highlighted (bottom). Paired t-test.

d) Organoids treated with DMSO or MS023 for 96h, and passaged to ENR for additional 96h. Images taken at 96h timepoint and 24 and 96h after passaging.

e) mRNA sequencing of untreated vs. MS023 treated organoids. GSEA for Gene Ontology biological process (GO:BP) terms. Normalized enrichment score (NES) is indicated by color. Top 50 categories are shown.

f) mRNA sequencing of untreated vs. MS023 treated organoids. GSEA for GO:BP terms "positive regulation of cell development" (GO:0010720, NES=1.6325), "lipid digestion" (GO:0044241, NES=1.9547), and "response to bacterium" (GO:0009617, NES=1.9257).

g) mRNA sequencing of untreated vs. MS023 treated organoids. GSEA for GO:BP term "DNA repair" (GO:0006281, NES=-1.5531).

h) PRMT tpm values in mRNA sequencing datasets of small intestinal crypts (3 BiolRep) and organoids (4 BiolRep) from Zwiggelaar et al. (control groups from E-MTAB-9077, E-MTAB-7862).

i) Box plots showing tpm values from plate based scRNA sequencing from Haber et al. (GSE92332). Tuft cell clusters have been merged into one group.

j) *Apc*-deficient adenomas treated with DMSO, MS023, or TC-E5003 for 96h. Representative well.

k) Organoids treated with DMSO, MS023, or TC-E5003 for 96h. Representative replicate.

l) Viability of organoids treated with DMSO, MS023, or TC-E5003 for 96h, measured by flow cytometry as frequency of Zombie Aqua-negative cells in Singlet parent gate. 3 biol. replicates, indicated by shape.
