## Supplementary Table1 for "A semi-automated organoid screening method demonstrates epigenetic control of intestinal epithelial differentiation"

**Table 1:**
**Inhibitors**

| Source | ProbeName | InternalProbeID | ProbeSynonyme | PubChem_CID | ChEMBL | ChEBI | InChIKey | InhibitorClass | Target | TargetSynonyme | StockConc [mM] | WorkingConc [µM] |
|---|---|---|---|---|---|---|---|---|---|---|---|---|
| Sigma Aldrich | DMSO | | | 679 | CHEMBL504 | CHEBI:28262 | IAZDPXIOMUYVGZ-UHFFFAOYSA-N | Ctrl | | | 10 | 10 |
| Cayman Chemicals | VPA | | | 3121 | CHEMBL109 | CHEBI:83766 | NIJJYAXOARWZEE-UHFFFAOYSA-N | Ctrl | | | 100 | 1000 |
| SGC | (+)-JQ1 | 1 | | 46907787 | CHEMBL1957266 | CHEBI:137113 | DNVXATUJJDPFDM-KRWDZBQOSA-N | BRD_BET | BRD2, BRD3, BRD4, BRDT (BET) | | 10 | 0.2 |
| SGC | bromosporine | 7 | | 72943187 | CHEMBL3133807 | CHEBI:95080 | UYBRROMMFMPJAN-UHFFFAOYSA-N | BRD_BET | pan-Bromodomain | | 10 | 1 |
| SGC | PFI-1 | 10 | PF-6405761 | 71271629 | CHEMBL2179387 | | TXZPMHLMPKIUGK-UHFFFAOYSA-N | BRD_BET | BRD2, BRD4 | | 10 | 1 |
| SGC | SGC-CBP30 | 16 | | 72201027 | CHEMBL3622373 | CHEBI:95079 | GEPYBHCJBORHCE-SFHVURJKSA-N | BRD_HAT | CREBBP, EP300 | CBP, P300 | 10 | 1 |
| SGC | I-CBP112 | 28 | | 90488984 | CHEMBL3774655 | | YKNAKDFZAWQEEO-IBGZPJMESA-N | BRD_HAT | CREBBP, EP300 | CBP, P300 | 10 | 3 |
| SGC | BI-9564 | 6 | | 117072549 | CHEMBL3823101 | | BJFSUDWKXGMUKA-UHFFFAOYSA-N | BRD | BRD9, BRD7 | | 10 | 1 |
| SGC | I-BRD9 | 27 | GSK602 | 91668541 | CHEMBL3769507 | | WRUWGLUCNBMGPS-UHFFFAOYSA-N | BRD | BRD9 | | 10 | 1 |
| SGC | LP99 | 32 | | 91827372 | CHEMBL3753082 | | LVDRREOUMKACNJ-BKMJKUGQSA-N | BRD | BRD9, BRD7 | | 10 | 1 |
| SGC | PFI-4 | 13 | | 40642506 | CHEMBL3356143 | | QCIJLRJBZDBVDB-UHFFFAOYSA-N | BRD | BRPF1B | | 10 | 1 |
| SGC | NI-57 | 35 | | 91827373 | CHEMBL3752151 | | UEMQPCYDWCSVCU-UHFFFAOYSA-N | BRD | BRPF1, BRPF2, BRPF3 | | 10 | 1 |
| SGC | OF1 | 37 | | 35397514 | CHEMBL3770173 | | YUNQZQREIHWDQT-UHFFFAOYSA-N | BRD | BRPF1B, BRPF2 | | 10 | 5 |
| SGC | BAZ2-ICR | 5 | | 91654625 | CHEMBL4296718 | | RRZVGDGTWNQAPW-UHFFFAOYSA-N | BRD | BAZ2A, BAZ2B | | 10 | 1 |
| SGC | GSK2801 | 9 | | 73010930 | CHEMBL3739699 | | KHWCPNJRJCNVRI-UHFFFAOYSA-N | BRD | BAZ2A, BAZ2B | | 10 | 3 |
| SGC | PFI-3 | 12 | | 78243717 | CHEMBL3752911 | | INAICWLVUAKEPB-QSTFCLMHSA-N | BRD | SMARCA,PB1 | | 10 | 1 |
| SGC | NVS-1 | 36 | | 117072550 | CHEMBL3785432 | CHEBI:149519 | XVECNLUKQDKOST-UHFFFAOYSA-N | BRD | CECR2 | | 10 | 1 |
| SGC | GSKJ4 | 25 | | 71729975 | CHEMBL3183531 | | WBKCKEHGXNWYMO-UHFFFAOYSA-N | KDM | JMJD3 | KDM6B | 10 | 5 |
| SGC | IOX2 | 30 | | 54685215 | CHEMBL3186774 | CHEBI:95077 | CAOSCCRYLYQBES-UHFFFAOYSA-N | KDM | HIF-1α prolyl hydroxylase-2 (PHD2), JMJD2, JMJD3, FIH | HPH1, EGLN2, KMD4A | 10 | 1 |
| SGC | GSK-LSD1 | 26 | | 71522234 | CHEMBL4301645 | CHEBI:95087 | BASFYRLYJAZPPL-UONOGXRCSA-N | KDM | LSD1 | KDM1A | 10 | 1 |
| SGC | IOX1 | 29 | | 459617 | CHEMBL1230640 | | JGRPKOGHYBAVMW-UHFFFAOYSA-N | KDM | 2OG oxygenases, including the JmjC demethylases | | 10 | 10 |
| SGC | UNC1999 | 20 | | 72551585 | CHEMBL3414619 | CHEBI:93239 | DPJNKUOXBZSZAI-UHFFFAOYSA-N | PMT_KMT | EZH2 | | 10 | 3 |
| SGC | GSK343 | 21 | | 71268957 | CHEMBL2204995 | | ULNXAWLQFZMIHX-UHFFFAOYSA-N | PMT_KMT | EZH2 | | 10 | 3 |
| SGC | OICR9429 | 38 | | 91623360 | CHEMBL3798846 | CHEBI:95086 | DJOVLOYCGXNVPI-UHFFFAOYSA-N | PMT_KMT | WDR5 | MLL1 | 10 | 3 |
| SGC | A-366 | 3 | | 76285486 | CHEMBL3109630 | | BKCDJTRMYWSXMC-UHFFFAOYSA-N | PMT_KMT | EHMT1, EHMT2 | G9a, GLP | 10 | 1 |
| SGC | UNC0638 | 17 | | 46224516 | CHEMBL1231795 | | QOECJCJVIMVJGX-UHFFFAOYSA-N | PMT_KMT | EHMT1, EHMT2 | G9a, GLP | 10 | 1 |
| SGC | UNC0642 | 18 | | 53315878 | CHEMBL2441082 | CHEBI:95074 | RNAMYOYQYRYFQY-UHFFFAOYSA-N | PMT_KMT | EHMT1, EHMT2 | G9a, GLP | 10 | 1 |
| SGC | PFI-2 | 11 | | 71300326 | CHEMBL3414622 | | JCKGSPAAPQRPBW-OAQYLSRUSA-N | PMT_KMT | SETD7 | | 10 | 1 |
| SGC | A-196 | 2 | | 117072548 | | | ABGOSOMRWSYAOB-UHFFFAOYSA-N | PMT_KMT | SUV420H1/H2 | | 10 | 1 |
| SGC | BAY-598 | 4 | | 117072551 | CHEMBL3818617 | | OTTJIRVZJJGFTK-SFHVURJKSA-N | PMT_KMT | SMYD2 | | 10 | 1 |
| SGC | MS023 | 33 | | 92136227 | CHEMBL3901808 | | FMTVWAGUJRUAKE-UHFFFAOYSA-N | PMT_PRMT | PRMT1, PRMT3, PRMT4, PRMT6, PRMT8 | | 10 | 5 |
| SGC | SGC707 | 15 | | 90642938 | CHEMBL4072005 | | DMIDPTCQPIJYFE-UHFFFAOYSA-N | PMT_PRMT | PRMT3 | | 10 | 1 |
| SGC | MS049 | 34 | | 53868701 | CHEMBL3961701 | | HBOJWAYLSJLULG-UHFFFAOYSA-N | PMT_PRMT | PRMT4, PRMT6 | | 10 | 5 |
| SGC | GSK591 | 23 | | 117072552 | | | TWKYXZSXXXKKJU-FQEVSTJZSA-N | PMT_PRMT | PRMT5 | | 10 | 1 |
| SGC | SGC0946 | 14 | | 56962337 | CHEMBL3087498 | | IQCKJUKAQJINMK-HUBRGWSESA-N | PMT_PRMT | DOT1L | | 10 | 1 |
| SGC | GSK484 | 22 | | 86340151 | | | MULKOGJHUZTANI-ADMBKAPUSA-N | PMT_PRMT | PAD-4 | PADI4 | 10 | 10 |
| SGC | UNC1215 | 19 | | 57339144 | CHEMBL2426364 | | PQOOIERVZAXHBP-UHFFFAOYSA-N | MBT | L3MBTL3 | | 10 | 1 |
| SGC | CI-994 | 8 | N-acetyldinaline, Tacedinaline | 2746 | CHEMBL235191 | | VAZAPHZUAVEOMC-UHFFFAOYSA-N | HDAC | HDAC1, HDAC2, HDAC3, HDAC8 | | 10 | 10 |
| SGC | LAQ824 | 31 | Dacinostat | 6445533 | CHEMBL356066 | CHEBI:90195 | BWDQBBCUWLSASG-MDZDMXLPSA-N | HDAC | HDAC | | 10 | 1 |
| SGC | GSK864 | 24 | | 91864701 | | CHEBI:94063 | DUCNNEYLFOQFSW-PMERELPUSA-N | other | Mutant isocitrate dehydrogenase 1 | IDH1 | 10 | 0.3 |
| SGC | olaparib | 39 | AZD2281, KU0059436 | 23725625 | CHEMBL521686 | | FDLYAMZZIXQODN-UHFFFAOYSA-N | other | PARP | | 10 | 1 |
| SantaCruz | TC-E5003 | | | 87052 | CHEMBL1797447 | CHEBI:39867 | SHRCVZJKZJGIHQ-UHFFFAOYSA-N | follow-up | PRMT1 | | 50 | 50 |
