## Supplementary Table2 for "A semi-automated organoid screening method demonstrates epigenetic control of intestinal epithelial differentiation"

**Table 2:**
**qRT-PCR primers & probes**

| Gene | IEC_lineage | Used_in_screen | Primer_fwd | Primer_rev | Probe | Product_length |
|---|---|---|---|---|---|---|
| Hprt | no | yes | cctcctcagaccgcttttt | aacctggttcatcatcgctaa | UPL_#95 | 91 |
| Alpi | Enterocyte | yes | aggatccatctgtcctttggt | ttcagctgccttcttgttcc | UPL_#33 | 75 |
| Lgr5 | Stem | yes | cttcactcggtgcagtgct | gatcagccagctaccaaatagg | UPL_#60 | 75 |
| Defa21 | Paneth | no | gagcagccaggggaagat | cgatttctacaaaggcagatca | UPL_#64 | 110 |
| Defa22 | Paneth | yes | gatgaagagactaatactgaggagca | cgttttctacaaaggcagatca | UPL_#64 | 131 |
| Defa29 | Paneth | no | tgaagagactaaaactgaggagca | caccacggaaaccttctttg | UPL_#6 | 99 |
| Lyz1 | Paneth | yes | ggcaaaaccccaagatctaa | tctctcaccaccctctttgc | UPL_#46 | 104 |
| Mptx2 | Paneth | no | tccaagaagagcacaatgga | aaatgcctttcccttcatgtc | UPL_#105 | 99 |
| Agr2 | Goblet | yes | cctcaacctggtctatgaaaca | accgtcagggatgggtcta | UPL_#2 | 93 |
| Ccl9 | Goblet | no | caagggcttgaaattgaaatg | tcttgaacactgaatccgtgag | UPL_#27 | 90 |
| Muc2 | Goblet | yes | cacgagacccaggaagtacag | gcaaagccactaactgcttgt | UPL_#105 | 88 |
| Chga | Enteroendocrine | yes | aacttcaagacctggctctcc | ctcaaagctgctgtgttgct | UPL_#33 | 131 |
| Chgb | Enteroendocrine | yes | cctctcaaatgccctatcca | cacctttgacctcttttccact | UPL_#47 | 92 |
| Cpe | Enteroendocrine | yes | ccggaagagactctcaaaagc | cggacaaaccctttaacacc | UPL_#105 | 95 |
| Gfr3a | Enteroendocrine | no | tgatcctgctactggtgctg | ctctgtggcaagggagtttc | UPL_#5 | 65 |
| Dclk1 | Tuft | yes | agaaggcacagcttgcaga | gttattgtggcaggaatctgg | UPL_#25 | 74 |
| Gfi1b | Tuft | yes | gttgctgaaccagagccttc | ttggggtgtcacgagagg | UPL_#45 | 64 |
| Lrmp | Tuft | no | gctgcttatggagactacacga | tccgttttctgtggttctga | UPL_#55 | 90 |
| Pou2f3 | Tuft | no | caaagcagcaatgaactcctc | tgattccaacggtaccatga | UPL_#78 | 69 |
| Trpm5 | Tuft | yes | ttctagatggtgtccaccaaaa | gcagaggggtccctgaat | UPL_#33 | 80 |
