## Supplementary Table3 for "A semi-automated organoid screening method demonstrates epigenetic control of intestinal epithelial differentiation"

**Table 3:**
**Materials, Reagents, Software**

| Item | Supplier | Specification | Publication | Comment |
|---|---|---|---|---|
| **Mice** | | | | |
| C57BL/6JRj wild type | Janvier Labs | | | |
| C57BL/6 wild type | | | | |
| B6D2 (C57Bl/6 x DBA/2) *Villin* -Cre | | | elMarijou2004 | kind gift from Sylvie Robine, Institut Curie-CNRS, Paris, France |
| C57BL/6J *Lsd1* $^{f/f}$ | | | Kerenyi2013 | kind gift from Stuart Orkin, Harvard Medical School , Boston , USA |
| B6.129P2 *Apc* $^{tm1Rsmi}$/RfoJ | Jackson Laboratories | stock no: 029275 | | |
| *Lgr5* -EGFP-IRES-CreERT2 | Jackson Laboratories | stock no: 008875 | | |
| *Hpgds* -tdTomato | | | Bornstein2018 | |
| *Neurog3* -RFP | | | Kim2015 | kind gift from Anne Grapin-Botton, DanStem, University of Copenhagen, Copenhagen, Denmark |
| **Cell lines** | | | | |
| cell line producing Noggin | | | | kind gift from Hans Clevers, Hubrecht Institute, Utrecht, The Netherlands |
| cell line producing R-Spondin | | | | kind gift from Calvin Kuo, Stanford University School of Medicine, Stanford, USA |
| **Plates** | | | | |
| 24-well plate | Corning | #3523 | | |
| 96-well glass-bottom plate | Cellvis | #P-96-1N | | |
| 384-well optical plates | Applied Biosystems | #4309849 | | |
| 8-well microscopy slide | Ibidi | #80821 | | |
| **Reagents** | | | | |
| PBS | Sigma-Aldrich | #806552 | | |
| EDTA | Thermo Fisher Scientific | #15575020 | | |
| TrypLE Express | Thermo Fisher Scientific | #12605010 | | |
| Tamoxifen | Sigma-Aldrich | #T5648 | | |
| Corn oil | Sigma-Aldrich | #C8267 | | |
| FCS | Gibco | #10270106 | | |
| Saponin | Sigma-Aldrich | #8047-15-2 | | |
| Triton X-100 | Sigma-Aldrich | #T8787 | | |
| PFA | Alfa Aesar | #43368 | | |
| **Organoid culture** | | | | |
| Matrigel | Corning | #734-1101 | | |
| DMEM F12 | Gibco | #31330-038 | | |
| Penicillin-Streptomycin | Sigma-Aldrich | #P0781 | | |
| HEPES | Gibco | #35050061 | | |
| Glutamax | Gibco | #35050061 | | |
| B-27 supplement | Gibco | #35050061 | | |
| N2 supplement | Gibco | #17502001 | | |
| N-Acetylcysteine | Sigma-Aldrich | #A-7250 | | |
| EGF Recombinant Mouse Protein | Thermo Fisher Scientific | #PMG8041 | | |
| CHIR99021 | Sigma-Aldrich | #SML1046 | | |
| Valproic acid (VPA) | Cayman Chemicals | #13033 | | |
| DAPT | Cayman Chemicals | #13197 | | |
| IWP-2 | Cayman Chemicals | #13951 | | |
| **RNA isolation, qRT-PCR** | | | | |
| RNA-solv Reagent | Omega Bio-Tek | #R6830-02 | | |
| Direct-zol MiniPrep kit | Zymo Research | #R2051 | | |
| Direct-zol-96 | Zymo Research | #R2055 | | |
| High-Capacity RNA-to-cDNA Kit | Applied Biosystems | #4388950 | | |
| 2x Perfecta ROX,UNG Fast Mix | Quanta Biosciences | #733-1398 | | |
| Primers | Sigma-Aldrich | | | see Supplementary Table 2 |
| Universal ProbeLibrary probes | Roche | | | see Supplementary Table 2 |
| **Antibodies** | | | | |
| **IF & IHC staining reagents** | | | | |
| BD CompBead Anti-Mouse Ig | Becton Dickinson | #552843 | | |
| BD CompBead Anti-Rat and Anti-Hamster Ig | Becton Dickinson | #552845 | | |
| DAPI | Thermo Fisher Scientific | #62248 | | |
| Hoechst 33342 | Thermo Fisher Scientific | #62249 | | |
| Zombie Aqua | Biolegend | #423102 | | |
| TruStain FcX | Biolegend | #101320 | | |
| CD24-PerCp-Cy5.5, clone M1/69 | Biolegend | #101824 | | |
| CD24-AF647, clone M1/69 | Biolegend | #101818 | | |
| CD44-AF647, clone IM7 | Biolegend | #103018 | | |
| CD44-BV785, clone IM7 | Biolegend | #103041 | | |
| CD117-PE-Cy7, clone ACK2 | Biolegend | #135112 | | |
| CD326-BV421, clone G8.8 | Biolegend | #118225 | | |
| CD326-BV605, clone G8.8 | Biolegend | #118227 | | |
| UEA1-FITC, Ulex Europaeus Agglutinin I | Invitrogen | #L32476 | | |
| UEA1-Rhodamine, Ulex Europaeus Agglutinin I | Vector Laboratories | #RL-1062-2 | | |
| Goat Anti-Rabbit IgG-BV421 | Invitrogen | #A-31556 | | |
| Goat Anti-Rabbit IgG-AF488 | Invitrogen | #A-11034 | | |
| anti-DCLK1 | Abcam | #ab31704 | | |
| anti-MUC2 | Santa Cruz | #sc-15334 | | |
| anti-KI67 | Invitrogen | #MA5-14520 | | |
| Fluoromount G | Invitrogen | #00-4958-02 | | |
| EnVision-HRP | Dako | #K4063 & #K4061 | | |
| DAB | Dako | #K5007 | | |
| **mRNA sequencing** | | | | |
| Quick-RNA MicroPrep kit | Zymo Research | #R1050 | | |
| Illumina TruSeq Stranded Total RNA | Illumina | | | I-CBP112 study, carried out at Genomics Core Facility, NTNU |
| Illumina NS500 MO flow-cell | Illumina | | | I-CBP112 study, carried out at Genomics Core Facility, NTNU |
| NEB Next Ultra RNA Library Prep Kit | NEB | | | MS023 study, carried out at Novogene (UK) Co |
| **Instruments** | | | | |
| EVOS FL Auto2 microscope | Thermo Fisher Scientific | | | |
| Eclipse Ci-L microscope | Nikon | | | |
| LSM880 confocal microscope | Zeiss | | | |
| Axio Imager Z1 confocal microscope | Zeiss | | | |
| NanoDrop-1000 | NanoDrop | | | |
| QuantStudio 5 Real-Time PCR system | Thermo Fisher Scientific | | | |
| BD LSRII flow cytometer | Becton Dickinson | | | |
| BD FACSAria III flow cytometer | Becton Dickinson | | | |
| MACSQuant X flow cytometer | Miltenyi Biotec | | | |
| Viaflo 96-channel pipette | Integra Biosciences | | | |
| Qubit Fluorometric Quantitation system | Life Technologies | | | |
| 2100 Bioanalyzer | Agilent | | | |
| Ilumina NextSeq 500 | Ilumina | | | |
| Illumina NovaSeq 6000 | Illumina | | | |

| | | | |
|---|---|---|---|
| **Software** | | | |
| GNU R | | v3.6.3 | |
| Bioconductor | | v3.10 | |
| tidyverse | | v1.3.0 | Wickham2019 |
| ggpubr | | v0.3.0 | |
| ggimage | | v0.2.8 | |
| magick | | v2.3 | |
| pheatmap | | v1.0.12 | |
| ggtree | | v2.0.4 | Yu2018 |
| flowCore | | v1.53.17 | |
| CytoML/flowWorkspace | | v1.12.1 | |
| ggcyto | | v1.14.1 | Van2018 |
| flowViz | | 1.50.0 | |
| biomaRt | | v2.42.1 | Durinck2005 |
| ClusterProfiler | | v3.14.3 | Yu2012 |
| ImageJ/Fiji | | v1.52n | Schindelin2012 |
| Ilastik | | v1.3.2 | Berg2019 |
| FlowJo software | FlowJo, LLC | v10.6.2 | |
| QuantStudio Design & Analysis Software | Thermo Fisher Scientific | v1.5.1 | |
| FastQC | | v0.11.8 | |
| MultiQC | | v1.7 | Ewels2016 |
| featureCounts | | v1.6.4 | Liao2014 |
| GNU R | | v3.6.1 | |
| STAR | | v2.7.3a | Dobin2013 |
| DESeq2 | | v1.26.0 | Love2014 |
| **Datasets** | | | |
| Ensembl | | GRCh38.p13 | |
| GENCODE annotation | | M18 | Frankish2019 |
| STRING-DB | | v11 | Szklarczyk2019 |
| TRRUST | | v2 | Han2018 |
| IEC lineage signatures | | GSE92332 | Haber2017 |
| LGR5+ stem cells | | GSE33949 | Munoz2012 |
| GO:BP | | GO Consortium | TheGeneOntologyConsortium2019 |
| tpm BRDs, PRMTs | | E-MTAB-9077 | Zwiggelaar2020 |
| tpm BRDs, PRMTs | | E-MTAB-78 | Zwiggelaar2020 |
| scRNA tpm Prmt1 | | GSE92332 | Haber2017 |
